# Identifying LasR quorum sensors with improved signal specificity by mapping the sequence-function landscape

**DOI:** 10.1101/2023.08.21.554225

**Authors:** Min Zeng, Biprodev Sarker, Stephen N. Rondthaler, Vanessa Vu, Lauren B. Andrews

## Abstract

Programmable intercellular signaling using components of naturally-occurring quorum sensing can allow for coordinated functions to be engineered in microbial consortia. LuxR-type transcriptional regulators are widely used for this purpose and are activated by homoserine lactone (HSL) signals. However, they often suffer from imperfect molecular discrimination of structurally similar HSLs, causing misregulation within engineered consortia containing multiple HSL signals. Here, we studied one such example, the regulator LasR from *Pseudomonas aeruginosa*. We elucidated its sequence-function relationship for ligand specificity using targeted protein engineering and multiplexed high-throughput biosensor screening. A pooled combinatorial saturation mutagenesis library (9,486 LasR DNA sequences) was created by mutating six residues in LasR’s β5 sheet with single, double, or triple amino acid substitutions. Sort-seq assays were performed in parallel using cognate and non-cognate HSLs to quantify each corresponding sensor’s response to each HSL signal, which identified hundreds of highly specific variants. Sensor variants identified were individually assayed and exhibited up to 60.6-fold (*p* = 0.0013) improved relative activation by the cognate signal compared to the wildtype. Interestingly, we uncovered prevalent mutational epistasis and previously unidentified residues contributing to signal specificity. The resulting sensors with negligible signal crosstalk could be broadly applied to engineer bacteria consortia.

## INTRODUCTION

The use of engineered bacterial consortia to distribute tasks among different strains has gained great interest in various applications, such as biomanufacturing (1, 2), bioremediation (3, 4), therapeutics (5, 6), and agriculture (7, 8). This strategy can offer reduced cellular burden of microbial members and the ability for specialization that leverages the microorganism’s traits (9–11). For engineered bacterial consortia to function robustly, programmable cell-cell communication is required to coordinate processes in different cells and control their spatiotemporal dynamics, which has been achieved via diffusible chemical intercellular signaling (12–15), cell-cell adhesion (16, 17), and conjugal transfer of DNA (18–20). In nature, bacteria commonly communicate via diffusible quorum sensing signals to elicit population-level and cell density-dependent responses (21–23). Various quorum sensing signals have been used as chemical signals in engineered microbial consortia, including oligopeptides (24, 25), γ-butyrolactone (26), and homoserine lactones (27–37). Among these, homoserine lactones (HSLs) have been most widely utilized in synthetic biology due to the relative ease of signal production and sensing (38–40). In these systems, the canonical LuxR-type allosteric transcription factor binds to its cognate HSL ligand, and after complexation, activates transcription from its corresponding quorum sensing promoter (41, 42). HSLs for LuxR-type regulators contain a lactone ring and commonly an acyl chain that can vary in length (4 to 20 carbons), degree of saturation, and oxidation state at the third carbon (43, 44). Structural similarity can cause non-cognate HSL signals to activate LuxR-type quorum sensors resulting in signal crosstalk (45–49), which can be problematic for precise control of functions in microbial consortia when using multiple quorum sensing systems.

Various methods have been employed to alter ligand specificity of allosteric transcription factors. Directed evolution via random mutagenesis (by error-prone PCR or gene shuffling) of the ligand binding domain or protein coding sequence has been widely used to engineer ligand specificity, including for improved molecular discrimination and for preferential specificity for molecules other than the natural cognate ligand (50–61). While random mutagenesis can introduce numerous mutations and generate sequence diversity, it often suffers from mutational bias that hinders screening all amino acid substitutions and therefore cannot comprehensively determine sequence-function relationships (62–65). Another approach is designing transcription factors using rational design that applies structure-based computational modeling and has also been used to achieve desired ligand specificity (66–71), yet it requires a large amount of prior knowledge and predictions of biophysical interactions that are often not known (72). For this study, we chose the middle ground of semi-rational design, which utilizes available structural and functional information of the transcription factor to choose the target residues and amino acid diversity for protein engineering (73–76). Semi-rational design can narrow the designed sequence space to a subset within the limits of what can be experimentally tested and predicted to have a greater likelihood for a desired functionality (77).

Altering the ligand specificity of LuxR-type quorum sensing regulators has been achieved by directed evolution and site-directed mutagenesis. Directed evolution was used to tune the specificity of different LuxR homologs (50–53). For TraR and LasR quorum sensing regulators with structural information, site-directed mutagenesis has been applied to alter their specificity (78, 79). More recently, evolutionary information predicted the residues essential for specificity and guided the site-directed mutagenesis for specificity change (80). Those residues dictating specificity in LuxR homologs are often not highly conserved or essential in quorum sensing regulators (80, 81). Despite these insights, how mutations at different residues contribute to specificity of quorum sensing regulators remains unknown.

Here, we investigate the sequence-function relationship of signal specificity for the LasR quorum sensing regulator from *P. aeruginosa*. We chose LasR for this study because its protein structure has been well studied, and the protein-ligand interactions with its cognate signal molecule *n*-(3-oxododecanoyl)-HSL (C12-HSL herein) are known (82–84). Additionally, LasR is known to have high signal crosstalk with the non-cognate signal *n*-(3-hydroxytetradecanoyl)-HSL (C14-HSL) (46, 53). Using combinatorial saturation mutagenesis, we created a library of 9,486 LasR DNA designs targeting a region in its β5 sheet (L125 – L130) (Figure 1A). We then adapted a pooled high-throughput sort-seq approach to characterize the response of each corresponding LasR sensor to cognate C12-HSL and non-cognate C14-HSL, identifying 559 variants (5.6% of designs) with high C12-HSL specificity (>3.7-fold improved compared to the wildtype LasR protein) (Figure 1B). Separate individual assays of identified variants exhibited improvement in the ratio of response from C12-HSL to C14-HSL. From this dataset, we identified positions and amino acids that affect signal specificity and quantified epistatic interactions. We also trained a neural network model, which predicted new LasR protein sequences having high specificity. Furthermore, we investigated the trade-off between sensitivity and specificity of the LasR quorum sensor. This work elucidates the sequence-specificity relationship of LasR in a 6-amino acid region, thus providing an essential foundation for the engineering of highly specific quorum sensors for use in complex microbial consortia.

**Figure 1.**
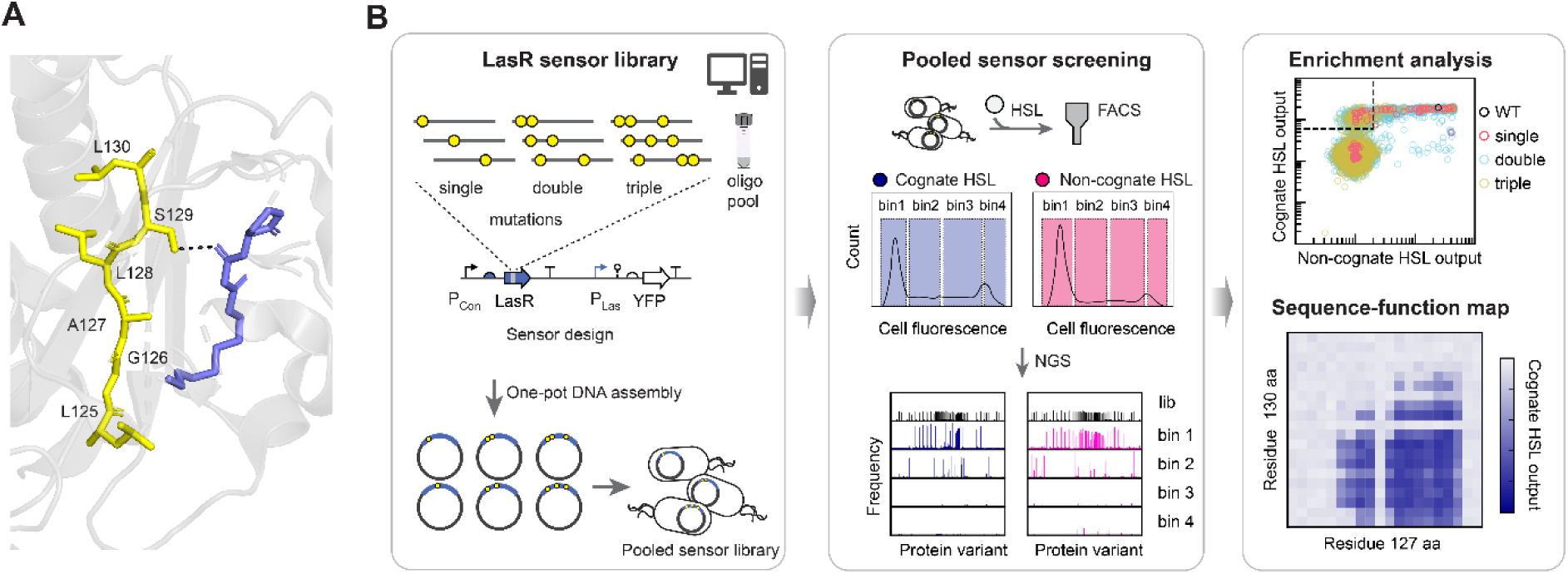
High-throughput engineering of signal specificity of LasR quorum sensor. **(A)** The residues in the β5 sheet (in yellow) of LasR (PBD: 3IX3 (83)) selected for mutagenesis are shown with the cognate signal molecule, N-3-oxododecanoyl homoserine lactone (C12-HSL, in blue). S129 forms a hydrogen bond with C12-HSL. Image was generated using The PyMOL Molecular Graphics System (Version 2.5.2 Schrödinger, LLC.). **(B)** Overview of the multiplexed approach for mapping the specificity landscape using combinatorial saturation mutagenesis. The LasR transcription factor library was constructed using an oligo pool to introduce single, double, and triple mutations, which was designed using a custom Python script (Methods). The oligo pool was amplified and assembled into the LasR sensor destination vector in a pooled Type IIS DNA assembly reaction. In the sensor construct, LasR is expressed by a constitutive promoter (P_Con_) and regulates the P_Las_ sensor output promoter that expresses a yellow fluorescent protein (YFP) reporter. The plasmid pool was transformed into *E. coli* for *in vivo* screening. After induction with the cognate HSL (blue filled circle) or non-cognate HSL (pink filled circle), cells were sorted using fluorescence activated cell sorting (FACS) into bins based on their fluorescence. The sensor plasmids were then extracted and sequenced by next-generation DNA sequencing (NGS) to quantify the frequency of each protein variant in each bin and the unsorted library (lib). Examples of data analysis are shown. Enrichment analysis plots compare the output of each sensor design (open circle) for the cognate and non-cognate HSLs. Heat map shows an example of the saturation mutagenesis data. The sequence-function relationship was mapped by plotting the sensor output for each combination of amino acids (aa) at different positions.

## RESULTS

### Design and construction of the LasR sensor library

In this study, we aimed to assess the protein sequence-function relationship for ligand specificity of LasR and to understand how problematic signal crosstalk for this biosensor could be mitigated. LasR is among a subset of well-studied LuxR-type quorum sensors that have been commonly used in combination to elicit multicellular bacterial responses (12, 32, 33, 98–100). The LasR symmetrical homodimer binds reversibly to its cognate ligand 3-oxo-C12-HSL (C12-HSL). However, ligand promiscuity and spurious activation of LasR by other long-acyl chain HSLs are well known and are especially prominent for 3-hydroxy-C14-HSL (C14-HSL), the cognate signal of the LuxR-type regulator CinR from *Rhizobium etli* (46, 49, 79) (Supplementary Figure S1). Importantly, the crystal structure of dimerized LasR bound to C12-HSL was previously determined (82). One molecule of C12-HSL binds to each LasR monomer in a ligand binding pocket comprised of a β-sheet (β5) and several α-helices (82). Therefore, we selected to study this system in this work. For the design of the library, we hypothesized that we could target mutations to residues in the reported ligand binding pocket to affect molecular interactions that would alter ligand discrimination and sensor activity.

For the design of our LasR mutagenesis library, we targeted mutations to six contiguous amino acid residues in the β5 sheet (L125 – L130) of LasR and designed 9,486 LasR regulator variants. We chose this region of LasR because it contained residues previously determined to interact with the acyl chain of HSL ligand (Figure 1A). Residue S129 forms a hydrogen bond with the acyl chain of C12-HSL, and residues L125, G126 and A127 have hydrophobic interactions with the acyl chain of C12-HSL (84, 101). Additionally, A127, S129 and L130 have been identified as essential for LasR specificity by covariation analysis (80). A central goal in this work was to assess the potential role and ability for mutational interactions to alter the differential activation of the sensor by two different HSLs, and therefore, we chose to apply combinatorial mutagenesis. With the aim of systematically quantifying the sequence-function relationship for the selected protein sequence space, we chose to apply saturation mutagenesis and utilized a chemically-synthesized pool of DNA oligonucleotides that allowed us to specify each mutation in the library.

A combinatorial saturation mutagenesis library was designed to contain all single and double amino acid substitutions in the LasR L125 – L130 region (Figure 2A). Each residue was mutated to one of 20 canonical amino acids, either individually to create 120 single mutation designs, or simultaneously pairwise to create 6,000 double mutation designs, including the wildtype protein sequence for both sets of designs. With space for additional designs on the oligonucleotide chip, we also included a subset of triple mutations. To limit the library size, only a small set of 4,000 LasR triple mutation variants were designed by mutating two residues simultaneously with the fixed mutation S129N. This S129N mutation was shown to improve the specificity of LasR in a previous study (79) and in our initial testing (Supplementary Figure S2). We chose this relatively moderate pool size with the aim of constructing a pooled library with representation of every design (i.e. the complete designed sequence space).

**Figure 2.**
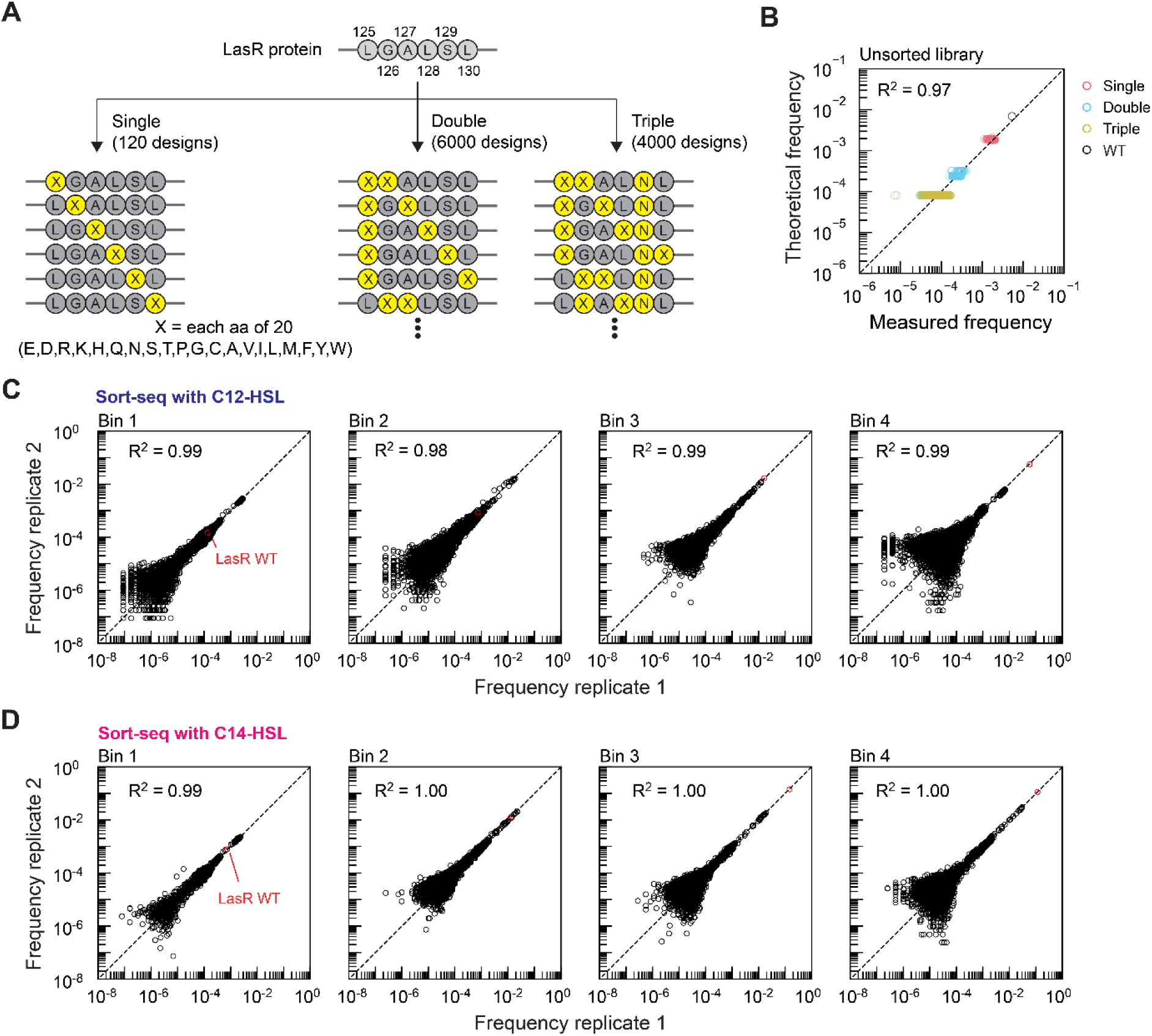
Sort-seq analysis of combinatorial saturation mutagenesis of LasR L125 – L130. **(A)** LasR protein sequences were designed to have each of the 20 natural amino acids (represented by X in yellow) at positions 125 to 130. For single amino acid substitution, each residue was mutated to each of the 19 other amino acids at each position, and the wildtype was included in pool. Double amino acid substitutions were designed by mutating each pair of residues simultaneously to achieve all combinations of amino acids. All triple mutants contained the S129N mutation while mutating two other residues simultaneously. The number of unique LasR DNA designs in the library is 9,486. **(B)** The measured frequency of each design in the constructed library was compared to the theoretical frequency, which was calculated as the frequency of each oligo in the designed oligo pool. Two aliquots of the library prior to sorting (unsorted) were sequenced and analyzed (Methods), and the average measured frequency of each variant is plotted. All 9,486 variants were identified by NGS in the constructed library. **(C)** Duplicate sort-seq assays with addition of 1 μM 3OC12-HSL (C12-HSL) were performed in two identical experiments to screen the pooled LasR sensor library. **(D)** Duplicate sort-seq assays with addition of 2 μM 3OHC14-HSL (C14-HSL) were performed. Cells were sorted into four bins, and each bin was sequenced and analyzed (Methods). The frequency of each variant in each experiment is plotted (open black circle). For comparison, the frequency of wildtype LasR (open red circle) in each bin is shown. The coefficient of determination (R^2^) was determined by linear least squares regression with a zero intercept. Dotted lines indicate the 1:1 relationship.

We wrote and used a custom Python script to automate the design of the set of oligos to create these LasR mutations (Methods). This script generates a CSV file of the oligo pool using an input configuration file specifying codon usage in the host organism, the number of point mutations, the residues to mutate, and specified amino acids at each position for mutations. For amino acid substitutions, the codon having the highest genomic codon usage frequency in *E. coli* was selected, except in cases when insertion of that codon would introduce a BsaI restriction site. In those cases, codon substitutions were prioritized in order of decreasing usage frequency. The script allows the user to specify disallowed DNA sequences (e.g. BsaI recognition sequence here) and other common features of the oligo, such as the oligo amplification sequences. For this study, each 80 nt single-stranded oligo contained the 18 nt LasR variable region flanked by BsaI recognition sequences along with orthogonal DNA linker sequences and outer 20 nt oligo amplification sequences (Supplementary Figure S3). In the specified oligo chip design, the number of single mutation designs was intentionally enriched 20-fold (2,400 oligo spots on the chip) with the aim of ensuring representation of all single mutation designs. To quantify sensor activation relative to the wildtype DNA sequence in the pooled assays, the oligo for the wildtype parent sequence was also included and designed to have the highest frequency of any oligo in the pool (enriched 86-fold on the chip). Overall, the oligo pool contained 12,400 oligos (0.99 Mbp total, 9,486 unique DNA sequences, 9,140 unique protein sequences for LasR) and was ordered as a 12K oligo chip.

Using the PCR amplified oligo pool and a multiplexed one-pot Type IIS DNA assembly reaction, we constructed the plasmid pool of sensor characterization constructs containing the designed LasR variants. The destination vector for the DNA assembly contained everything of LasR sensor except the LasR variable region (Supplementary Figure S3). The design of the sensor characterization construct used here follows previously established standardized genetic architectures for assaying the output of the sensor promoter in relative promoter units (RPU), which are relative to the reference plasmid (pAN1717) designated as 1 RPU (88, 89). By doing this, the sensor response measured on different instruments can be quantitatively compared. The low-copy (p15A origin) sensor characterization plasmid constitutively expressed LasR, and the activity of the P_Las_ sensor output promoter was measured using an insulated yellow fluorescent protein reporter (ribozyme, RBS, eYFP CDS, and terminator identical to the pAN1717 reference plasmid, Supplementary Figure S3). The assembled plasmid pool was electroporated into *E. coli* NEB 10-beta to screen the LasR sensors *in vivo*.

We evaluated the composition of the constructed LasR sensor library by sequencing two aliquots of the unsorted library after molecular barcoding and next-generation sequencing preparation. Given the low nucleotide diversity at unmutated residues of LasR, which can largely reduce Illumina sequencing output and read quality (102, 103), we designed a set of custom primers for each sample to append 3 – 4 nt to increase the nucleotide diversity (Methods). A region of LasR containing the 18-bp variable region, was sequenced using 2 x 75 bp paired-end DNA sequencing. We then determined the number of perfect-match read counts for the sequenced LasR region (34 bp), defined as those having the exact sequence for the designed variant. At least 6,200 perfect-match read counts were identified for each design of the LasR single mutation variants (Supplementary Figure S4A). At least 170 perfect-match read counts were identified for 99.98% of the LasR double and triple mutation variants (9,367 out of 9,369), except for two LasR designs (G126D-A127G: 44 read counts and G126D-A127G-S129N: 39 read counts) (Supplementary Figure S4B). We observed each design in the unsorted library with all LasR designs represented in the pool (100% coverage for the library). The frequency of each variant was calculated as the ratio of its perfect-match read counts to the total number of perfect-match read counts for the sample. The average frequency of variants in the constructed pooled library strongly correlated with the frequency of the corresponding oligos in the designed oligo chip (R^2^ = 0.98) (Figure 2B). The frequency of variants in two unsorted libraries was highly correlated (R^2^ = 0.997, Supplementary Figure S5).

### Pooled screening of LasR sensor variants for signal specificity

Next, we screened the functional activity of the constructed LasR sensor library using a high-throughput approach known as sort-seq that utilized fluorescence-activated cell sorting (FACS), molecular barcoding, and next-generation sequencing (NGS). The sort-seq approach is most commonly used for screening biosensor activation by one ligand. However, measuring signal crosstalk and ligand specificity requires quantifying the activation of the sensor variants to each of multiple signals. To achieve this, we performed pooled sort-seq experiments to measure the activity of each LasR variant to cognate and non-cognate HSLs in parallel, which circumvents performing two sequential rounds of dual selections in directed evolution approaches used for identifying protein variants with improved specificity (50, 52). With the aim of assessing specificity, here we selected one concentration for each ligand, which was specified to be the typical concentration for activation of the cognate sensor based on previous reports and measurements of the wildtype LasR dose-response curves (45, 46) (Supplementary Figure S1).

Pooled sensor characterization was performed for the LasR sensor library in *E. coli* cells with addition of either 1 µM C12-HSL or 2 µM C14-HSL. For each experiment, cells were sorted into four bins by FACS based on cell fluorescence from YFP expression, which is the fluorescent reporter to measure the P_Las_ sensor promoter output. The fluorescence threshold for the first bin was set to the autofluorescence of *E. coli* 10-beta cells (without plasmid). The thresholds for the three remaining bins were specified to split the rest of the library into bins of roughly equal proportions of cells. The sort-seq experiments were performed in two replicates to assess reproducibility, and at least 3.5 million cells were sorted for each sample. Cells containing the RPU standard plasmid (pAN1717) were also analyzed on the cell sorter so that the sensor output could be converted to RPU. Cells collected in each bin were cultured, and then plasmids were extracted, barcoded, and pooled together for next-generation Illumina sequencing (Methods). A total of 102.5 million perfect-match reads were obtained (92.4% of all reads), which provided 5.6-fold average sequencing coverage for cells in all 16 sorted bins. For the replicate experiments, the frequency of the variants in each bin was strongly linearly correlated for both the C12-HSL sort-seq (R^2^ = 0.98 – 0.99, Figure 2C) and C14-HSL sort-seq (R^2^ = 0.99 – 1.00, Figure 2D) experiments. By including the original wildtype LasR sensor in the pool, the frequency of each variant can be directly compared to the frequency of the wildtype LasR sensor (red markers in Figure 2C, 2D).

### Validation of the pooled sensor screening results by individual assays of 41 LasR variants

Next, we analyzed the enrichment of LasR variants from the sequencing data. Due to cell-to-cell variability and imperfect accuracy of cell sorting, not every cell containing an identical genetic design is expected to have the exact same fluorescence and be found in one single bin. We sought to use the distribution of each sensor variant across the bins to infer the sensor output for each ligand assayed (1 µM C12-HSL or 2 µM C14-HSL). To do this, the output of the sensor was calculated as the weighted sum of the median cell fluorescence in the bin multiplied by the proportion of the adjusted perfect-match read counts of each LasR variant in the bin, respectively (Methods). The bin having lowest fluorescence (bin 1) contained most of the cells during sorting (67.4 - 84.5% in each sorting experiment), and only up to 2 million cells per bin could be collected. Therefore, we applied a scaling factor to normalize the read counts for this bin. To be able to quantitatively compare the sensor response measured on the cell-sorter to an analytical flow cytometer (86, 87), the measured cell fluorescence in arbitrary units was converted to relative promoter units (RPU) for the P_Las_ sensor output promoter (Methods). To identify variants with high specificity for C12-HSL, we defined limits for the sensor output to C12-HSL and C14-HSL (output with C12-HSL > 6.0 RPU and output with C14-HSL output < 0.2 RPU).

By applying this criterion, sixteen single-mutation variants having high specificity to C12-HSL were identified from the library (13.8% of single mutant designs) (Figure 3A). This set of sensor variants were individually constructed and assayed by flow cytometry (Figure 3B). The mutations among this set were at all amino acid positions, except position 126. One mutation at positions 125 and 129 were included in the identified variants (L125E and S129N), and multiple mutations at positions 127, 128, and 130 were identified (4 mutations at 127, 5 mutations at 128, and 5 mutations at 130). The S129N mutation, which was previously identified to improve C12-HSL specificity, was among the single mutations identified here (79). Among the 16 single-mutation variants, the output to C14-HSL for all variants was less than the wildtype LasR sensor (*p* = 0.0039 – 0.0048), ranging from 14.6-fold to 65.5-fold lower output. Whereas the wildtype LasR sensor had 422.9-fold induction to C12-HSL and 62.9-fold induction to C14-HSL, these 16 sensor variants had 146.7-fold to 541.9-fold induction to C12-HSL and only 1.1-fold to 4.3-fold induction to C14-HSL. To quantify the sensor specificity, we defined specificity as the difference between the natural logarithm of the C12-HSL output relative to the C14-HSL output for the variant and wildtype LasR (Methods). Therefore, the specificity for the sensor containing wildtype LasR is equal to Zero. Variants having a greater ratio of output to C12-HSL relative to C14-HSL have a positive specificity score (i.e. improved specificity for C12-HSL). The specificity (*S*) for these 16 single-mutation variants ranged between 1.95 ± 0.16 to 3.75 ± 0.26 with highest specificity for A127N. Interestingly, four of these mutations at position 128 (hydrophobic residues cystine, isoleucine, valine, tryptophan, and tyrosine), which to our knowledge has not previously been reported as contributing to C12-HSL molecular recognition.

**Figure 3.**
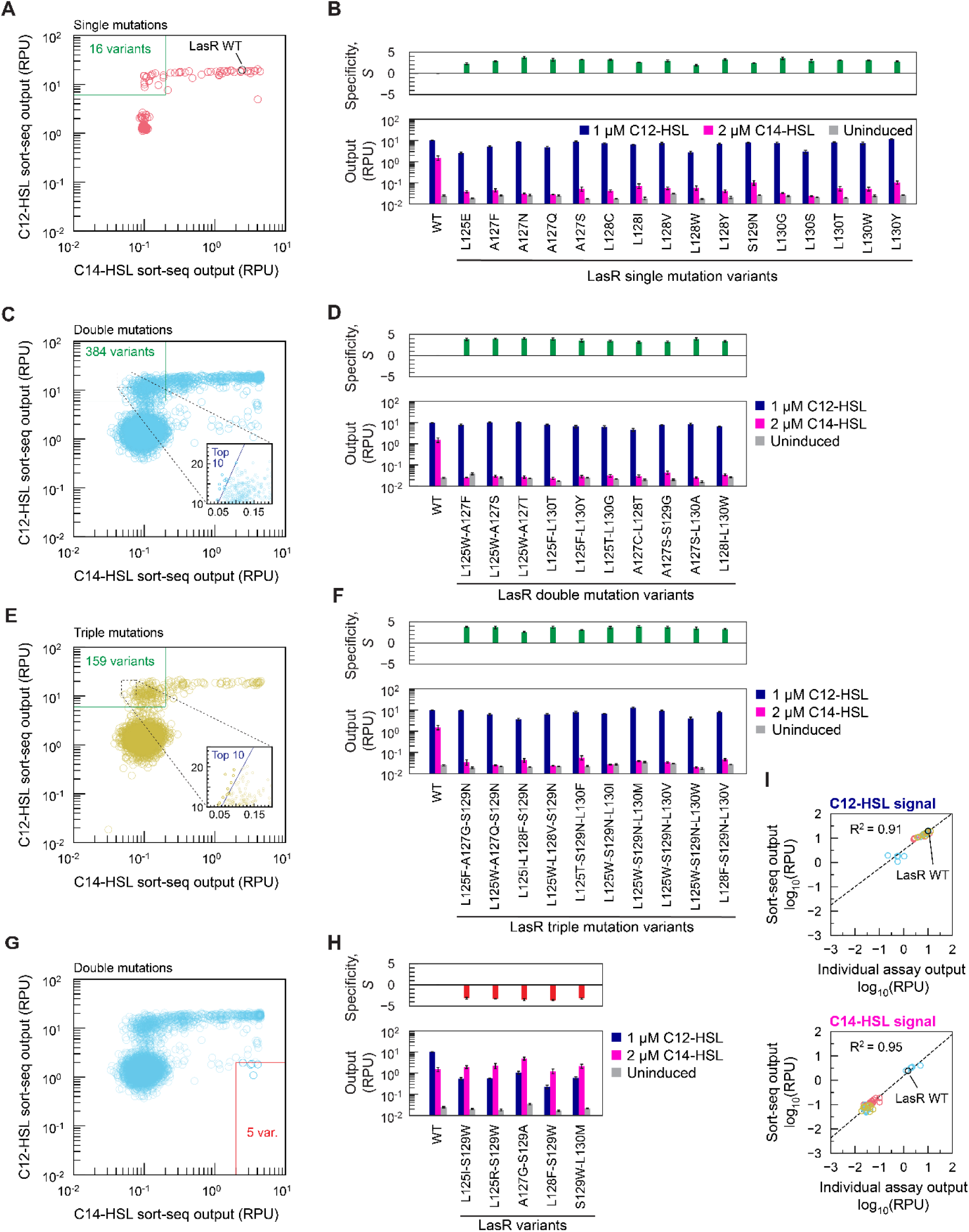
Validation of sort-seq results by individual assays of biosensors containing LasR variants. **(A)** The average sort-seq output of sensors containing LasR with one amino acid substitution is plotted for sort-seq assays with 1 μM 3OC12-HSL (C12-HSL) or 2 μM 3OHC14-HSL (C14-HSL), respectively. The wildtype LasR sensor (WT) is plotted in black for comparison. Output for each LasR variant was computed from the distribution of read counts in the sorted bins, and fluorescence was converted to relative promoter units (RPU) (Methods). Variants with improved C12-HSL specificity were determined by specifying average output thresholds of C12-HSL output > 6 RPU and C14-HSL output < 0.2 RPU (green box), and 16 single mutations variants met this criterion. **(B)** Sensors containing each of the 16 LasR single mutations identified were individually constructed. Sensor characterization assays using flow cytometry were performed individually for each construct on three separate days. Bars indicate the average output for a population of at least 10,000 cells, and error bars are the s.d. (*n* = 3). The specificity (*S*) of each sensor was calculated for each day as defined in the methods. Bars indicate the mean ± s.d. (*n* = 3). **(C)** The average sort-seq output of sensors containing LasR with two amino acid substitutions is plotted for sort-seq assays with 1 μM C12-HSL or 2 μM C14-HSL, respectively. Using the criterion for improved C12-HSL specificity (green box), 384 double mutation variants were identified. Of those, 10 top variants having the highest specificity were selected, indicated by a blue line in the inset plot. **(D)** Sensors containing each of those 10 LasR double mutations were individually constructed and assayed by flow cytometry on three separate days without inducer or with addition of C12-HSL or C14-HSL. The average output and specificity are plotted. Bars indicate the mean ± s.d. (*n* = 3). **(E)** Average sort-seq output of sensors containing LasR with three amino acid substitutions is plotted. Using the criterion for improved C12-HSL specificity (green box), 159 triple mutation variants were identified. Of those, 10 top variants were selected, indicated by a blue line in the inset plot. **(F)** Sensors containing each of those 10 LasR triple mutations were individually constructed and assayed by flow cytometry on three separate days without inducer or with addition of C12-HSL or C14-HSL. The average output and specificity are plotted. Bars indicate the mean ± s.d. (*n* = 3). **(G)** From the subset of double mutation LasR variants, a few surprisingly displayed preferential activation by C14-HSL. Five variants with specificity for C14-HSL were identified by applying thresholds of C12-HSL output < 2 RPU and C14-HSL output > 2 RPU (red box). **(H)** Each of these five LasR sensors were individually constructed and assayed without inducer (light grey), with 1 µM C12-HSL (dark blue) or 2 µM C14-HSL (magenta). **(I)** For all 41 sensors individually assayed (panels B, D, F, H), the measured output with C12-HSL (top panel) or C14-HSL (bottom panel) is compared to the prediction from the corresponding sort-seq dataset in log scale. The coefficient of determination (R^2^) was determined by least squares regression with intercepts. Dotted lines indicate the resulting fit.

Using the same criterion for high C12-HSL specificity, 384 double-mutation LasR variants in the library (6.85% of double mutant designs) were found to have high specificity (Figure 3C). We chose the ten of them having the highest *S* scores (Figure 3C inset), and these variants were individually constructed and assayed (Figure 3D). For these 10 sensor variants, the output with 2 µM C14-HSL was less than the wildtype LasR sensor (*p* = 0.0039 – 0.0041) and reduced by 35.1-fold to 66.7-fold, while the C12-HSL output remained high (7.62 ± 1.83 RPU for these 10 sensors). The specificity (*S*) of these double-mutation variants ranged from 3.14 ± 0.24 up to 4.08 ± 0.20. In some cases, each of the two combinatorial mutations in LasR were found to improve C12-HSL specificity in the single-mutation subset (e.g. L128I and L130W). In other cases, one mutation that decreased C12-HSL specificity when introduced alone augmented signal discrimination in combination with another mutation. For example, LasR L125W decreased C12-HSL specificity (*S* = −0.38 ± 0.12, Supplementary Figure S6), yet combined with the mutation A127F (*S* = 2.89 ± 0.11) or A127S (*S* = 3.27 ± 0.07), specificity increased more than purely additively (L125W-A127F, *S* = 3.85 ± 0.30 or L125W-A127S, *S* = 3.95 ± 0.19).

Of the triple-mutation variants, 159 were identified (4.23% of triple mutant designs) that have high C12-HSL specificity, again applying the same criterion (Figure 3E). Of these, the ten triple-mutation variants having the highest specificity were individually assembled and sensor characterization assays were performed (Figure 3F). The specificity (*S*) of these triple-mutation variants ranged from 2.58 ± 0.13 up to 3.90 ± 0.24. The mutation L125W, which is deleterious alone (Supplementary Figure S6), was present in six of the top ten triple-mutation variants. Interestingly, this mutation was also found in three of the top double-mutation variants above. Among all individually assayed sensor variants, four LasR double-mutation variants (A127S-L130A, L125F-L130T, L125W-A127S, and L125W-A127T) exhibited the highest measured C12-HSL specificity (*S* > 3.9).

Reversing ligand specificity of an allosteric transcriptional factor can require cooperative interactions of mutations in different regions of the ligand binding pocket (66, 70, 73, 104), yet here, we identified a small subset of designs with engineered preference for C14-HSL. By setting thresholds for ligand specificity for C14-HSL (C12-HSL output < 2 RPU and C14-HSL output > 2 RPU), we identified 5 variants in the library (0.11% of library) that were inferred to maintain high activation by C14-HSL and have a lower activation by C12-HSL (Figure 3G). All 5 variants contained double mutations in LasR, and we individually constructed and assayed each of them (Figure 3H). All variants have significantly decreased output (*p* < 0.001) with C12-HSL (0.60 ± 0.28 RPU) compared to wildtype LasR. Relative to the uninduced basal promoter activity, these five sensors have higher induction by C14-HSL (78.6 ± 18.6-fold to 142.8 ± 11.6-fold) than C12-HSL (14.3 ± 2.7-fold to 33.7 ± 3.4-fold). Indeed, the measured specificity scores (*S*) of these variants were highly negative, ranging from −3.61 ± 0.15 to −3.18 ± 0.22, showing engineered specificity for the C14-HSL ligand relative to wildtype LasR. Four of five variants contained S129W (from pooled sort-seq assays, *S* = −1.9), and this mutation was reported to alter specificity and preferentially bind HSL ligands having longer acyl chain length (79).

To further test the inferred sensor performance from the sort-seq assays, we compared the measured output from the individual assays to the sort-seq output using linear regression analysis for the 41 sensor variants and the wildtype LasR sensor. Overall, they exhibited strong linear correlation for both the C12-HSL assays (R^2^ = 0.91) and C14-HSL assays (R^2^ = 0.95) (Figure 3I). These results support that the pooled screening by high-throughput sort-seq accurately determined the sensor responses. While each design was not individually assayed, in total 559 designs (5.7% of the whole library) exhibited comparable C12-HSL output with the wildtype LasR sensor and low C14-HSL output (shown as green boxes in Figure 3 scatter plots). Next, the sequence-function relationship for this variable region of LasR was analyzed to comprehensively understand how protein sequence mutations affected ligand specificity (82, 84).

### Sequence-function relationships for LasR specificity and sensor activation

In our sort-seq experiments, we assayed sensors containing all single and double amino acid substitutions in the LasR variable region (positions 125-130) and a subset of triple mutations (i.e. all containing S129N). From the sort-seq data, we assessed the sequence-function relationships for sensor activation by C12-HSL, sensor activation by C14-HSL, and ligand specificity (*S*). We mapped each functional attribute of the sensor against the identities of the amino acids at each pair of positions in the protein sequence, and we placed amino acids with similar physicochemical properties next to each other in the heat maps (Figure 4A, and Supplementary Figure S7 – S9). In the heat maps, we indicate the wildtype amino acid (outlined in red) for each position, and therefore, we can analyze the single and double mutations from these plots. LasR variants with high activation to either C12-HSL or C14-HSL (defined here as ≥ 50% output of wildtype LasR, which is ≥ 9.85 RPU for C12-HSL and ≥ 1.22 RPU for C14-HSL) often clustered together around the wildtype amino acid in each position, yet the distribution of amino acids in these clusters varied based on the position (Figure 4A, and Supplementary Figure S7 – S9). For example, 15 amino acid substitutions (all except proline, glycine, aspartic acid, and glutamic acid) at position 125 resulted in high activation to C12-HSL (≥ 9.85 RPU) (Figure 4A and Supplementary Figure S10A). In contrast, none of the amino acid substitutions at position 126 maintained high activation to C12-HSL, and on average, the output decreased 15-fold for any mutation at this position (1.34 ± 0.16 RPU for the 19 substitutions) (Supplementary Figure S10A). All amino acid substitutions at position 126 also eliminated sensor activation by C14-HSL (4 – 5% of wildtype LasR) (Figure 4A and Supplementary Figure S10B). For both C12-HSL and C14-HSL, we observed that proline substitution at any position decreased sensor activation and eliminated activation for most designs (17-fold reduced sensor output on average) (Figure 4A and Supplementary Figure S10, S11), which is not surprising. Proline substitution increases conformational rigidity in a protein’s structure and has inactivated other proteins (105, 106). For all positions, more mutations resulted in high sensor activation by C12-HSL (42.2% of single mutation and 12.9% of double mutations on average) than by C14-HSL (17.2% of single mutation and 4.2% of double mutations on average) (Supplementary Figure S10, S11). In many cases, amino acid residues contributing to specificity resulted from mutation of amino acid identities at a position that affected the activation by C12-HSL and C14-HSL differently. For example, single substitutions with polar amino acids (glutamine, asparagine, threonine, and serine) at position 127 increased LasR specificity (*S* = 2.54 ± 0.37 for 4 designs), which resulted from decreasing the sensor activation by C14-HSL 18.5-fold on average while only decreasing the sensor activation by C12-HSL 1.4-fold on average (Figure 4A).

**Figure 4.**
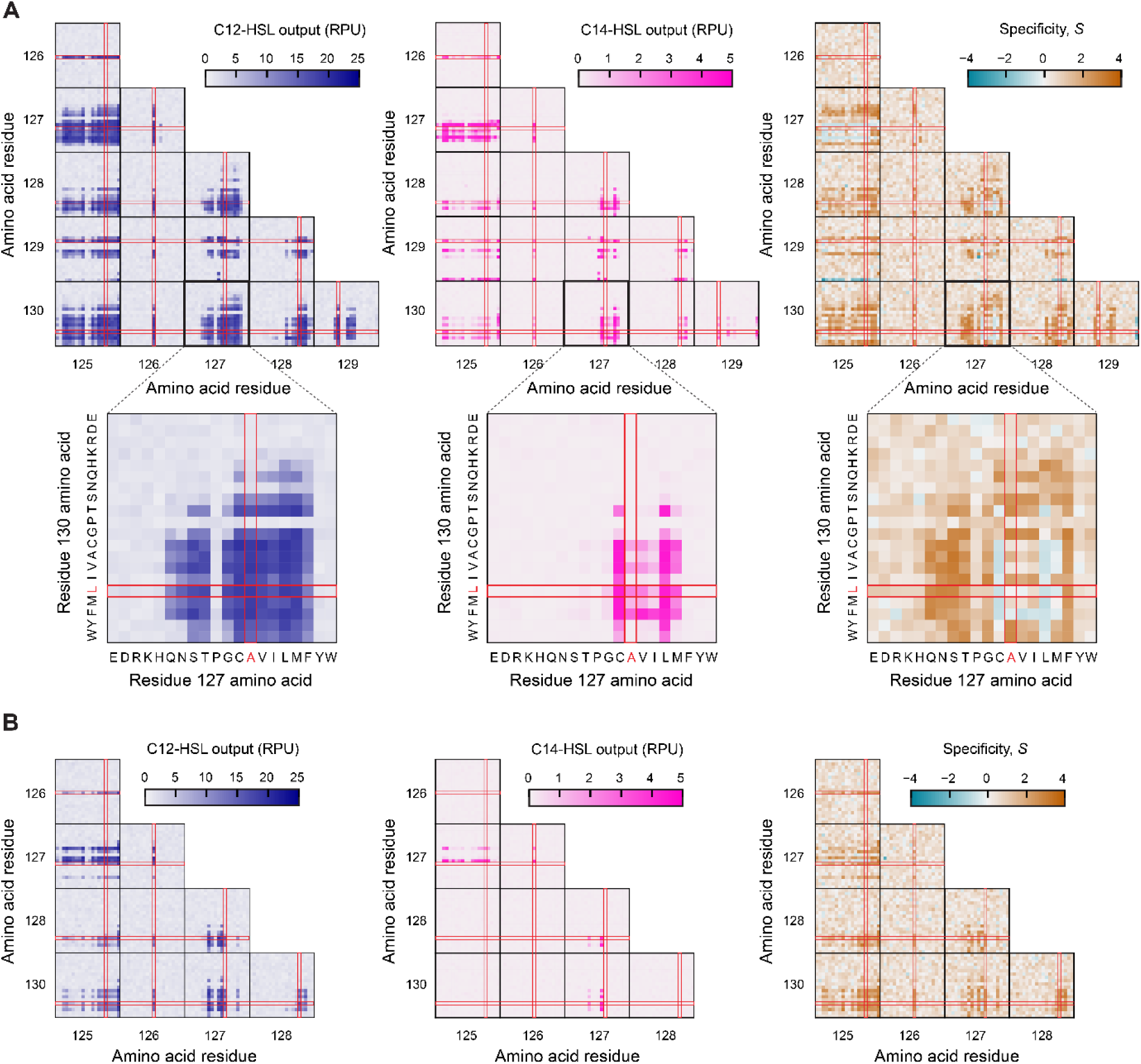
Sequence-function relationships for ligand specificity of LasR from combinatorial saturation mutagenesis. **(A)** Single and double mutations LasR sort-seq data depicted in heat maps. Each heat map shows the C12-HSL output (left), C14-HSL output (middle) and specificity (right) as determined by sort-seq assays for each combination of amino acids (20) at each position 125-130. Heat maps are arrayed to show all combinations of amino acid positions 125 – 130. Plots for positions 130 versus 127 are enlarged as an example. Amino acids with similar physicochemical properties were placed together in the heat maps, and identical ordering is used in all heat maps. The wildtype amino acid at each position is indicated (red lines). Larger images of each plot are in Supplementary Figure S7 – S9. **(B)** LasR sort-seq data for variants containing three amino acid substitutions depicted in heat maps. All LasR designs contained mutation S129N by design, so position 129 is not depicted. The C12-HSL output (left), C14-HSL output (middle) and specificity (right) of variants with triple mutations are shown. Larger versions of each plot are in Supplementary Figure S12 – S14. C12-HSL output and C14-HSL output were determined by the average of two replicate sort-seq assays using 1 µM 3OC12-HSL (C12-HSL) or 2 µM 3OHC14-HSL (C14-HSL), respectively (Methods). Specificity was calculated from the average C12-HSL output and average C14-HSL output using the function defined in the Methods.

For the triple-mutation LasR variants, we also mapped the functional attributes of the sensor against the identities of the amino acids at each position (Figure 4B and Supplementary Figure S12 – S14). Since all triple mutants contained the S129N mutation, it is not shown in the heat maps, and all other pairs of positions were analyzed (Figure 4B). Compared to the double-mutation variants, far fewer triple-mutation variants maintained high sensor activation by both HSL species. For C12-HSL, only 4.7% of all triple-mutants maintained sensor activation (≥ 9.85 RPU) (Supplementary Figure S15A), compared to 12.9% of the double-mutation variants (Supplementary Figure S11A). For C14-HSL, only 0.7% of all triple-mutants maintained sensor activation (≥ 1.22 RPU) (Supplementary Figure S15B), compared to 4.2% of the double-mutation variants (Supplementary Figure S11B). However, interestingly, many triple-mutation variants containing L125W maintained high sensor activation by C12-HSL (28.6%, 22 out of 77 variants) (Supplementary Figure S15A), and these sensors all had low activation by C14-HSL (0.12 ± 0.07 RPU) (Supplementary Figure S15B). By comparing the sensors having triple and double mutations, we see that the mutational tolerance at position 127 was changed by the S129N mutation. For example, none of the substitution with a subset of nonpolar amino acids (valine, isoleucine, leucine; 234 variants total) at position 127 in the triple mutants resulted in high sensor activation by C12-HSL, whereas 41.5% of the corresponding double mutants without the S129N mutation (291 variants total) achieved high sensor activation by C12-HSL. For the sensor activation by C14-HSL, only triple-mutants containing serine or cysteine at position 127 had high activation (Supplementary Figure S15B), while double-mutation variants containing A127S had low activation (0.11 ± 0.03 RPU for 95 variants) yet substitutions with 5 other amino acids (glycine, valine, isoleucine, leucine, methionine) maintained high activity (Supplementary Figure S11B). These sequence-function heat maps present visually how each functional attribute of the sensor is affected by the amino acid substitutions in LasR.

### Developing neural network models for the sequence-function relationship for LasR specificity

While combinatorial saturation mutagenesis can exhaustively determine the sequence-function landscape, relationships can often be determined and modeled using sparser data and supervised machine learning. We next wanted to investigate models relating the LasR protein sequence to C12-HSL ligand specificity that could be developed from our data. Here, we used a supervised deep-learning platform for protein sequence-function mapping developed by Gelman, Romero, Gitter, et al (94) to compare linear and nonlinear models to predict the specificity of LasR from its protein sequence. First, we reduced our library dataset to 9,140 unique protein sequences by removing synonymous codons, and then, randomly split the dataset into training (60% library), tuning (20% library), and testing (20% library) subsets. The inputs to the software were the whole LasR protein sequence and a file listing for each protein variant the number of mutations, identity of the mutations, and the functional score, which was here specified to be the calculated specificity (*S*) (Supplementary Spreadsheet, Tab 1). We generated and compared the linear regression, fully connected network, and convolutional neural network models. For predicting the specificity of the testing set variants, the convolutional neural network model showed a strong correlation (Pearson coefficient, *ρ* = 0.72 ± 0.01, *n* = 3) and greater agreement than both the linear regression model (*ρ* = 0.45 ± 0.004, *n* = 3) and the fully connected network model (*ρ* = 0.58 ± 0.005, *n* = 3) (Supplementary Figure S16). This agrees with previous studies that have shown that convolutional neural networks can capture nonlinear interactions of mutations and learn weights of amino acids to outperform other models (94, 107–114). To assess the tradeoff between accuracy and size of the dataset for generating the convolutional neural network model, we next tested the effect of decreasing the size of the training and tuning datasets using a 3:1 proportionality for all. With decreasing the size of the training and tuning sets 10-fold (6% training, 2% tuning), the correlation for the test set was largely reduced (Pearson *ρ* = 0.46 ± 0.01, *n* = 3), yet a 2-fold decrease in the size of the training and tuning sets (30% training, 10% tuning) only reduced the model agreement by 9.7% (*ρ* = 0.65 ± 0.01, *n* = 3) (Supplementary Figure S17).

We then used the convolutional neural networks to design new LasR variants predicted to have a desired specificity (Figure 5A). A random-restart hill-climbing algorithm in the platform was used to search LasR designs with maximal specificity for mutations constrained to positions 125 – 130 (94). We applied this searching method to identify three LasR variants with three, four, or five mutations, respectively, predicted to have the greatest C12-HSL specificity (*S*). Given that we also identified variants with engineered preference for C14-HSL in the library, we also constructed a convolutional neural network for the reversed specificity (−1 × *S)* output function (Supplementary Figure S18), which corresponds to preferential specificity for C14-HSL relative to C12-HSL. Then, we used the same algorithm and identified two designs predicted to have maximal reversed specificity. Sensors containing each of the five LasR variants identified by the neural networks were constructed and individually assayed. All three variants predicted to have maximum specificity (*S*) had significantly decreased C14-HSL output relative to the wildtype (*p* = 0.031), showing 59.6 - 68.5-fold reduction and approaching the uninduced output of each sensor (Figure 5B). Among those three, the LasR triple mutation variant L125W-A127T-L128M showed the highest specificity (*S* = 4.16 ± 0.44), yet this was not a significant improvement in specificity (*p* = 0.84) compared to the LasR variant L125W-A127T that demonstrated the highest C12-HSL specificity in the designed library (*S* = 4.08 ± 0.20). The sensor output with C12-HSL was significantly less for the LasR variants with four or five mutations than the wildtype LasR (*p* < 0.001), which is not surprising given that the neural network used the specificity as the functional score and not the promoter activity for each signal. The two LasR variants predicted to have the maximum reversed specificity exhibited specificity less than zero (Figure 5B). However, while one variant (L125R-L128F-S129W) maintained an equivalent promoter response to C14-HSL as the wildtype LasR, the other had 48.3-fold lower promoter output. The predictions from the convolutional neural networks had much greater agreement with experimental measurements for the triple mutants (root mean square error, RMSE = 0.12) than variants containing more than three mutations (RMSE = 1.36).

**Figure 5.**
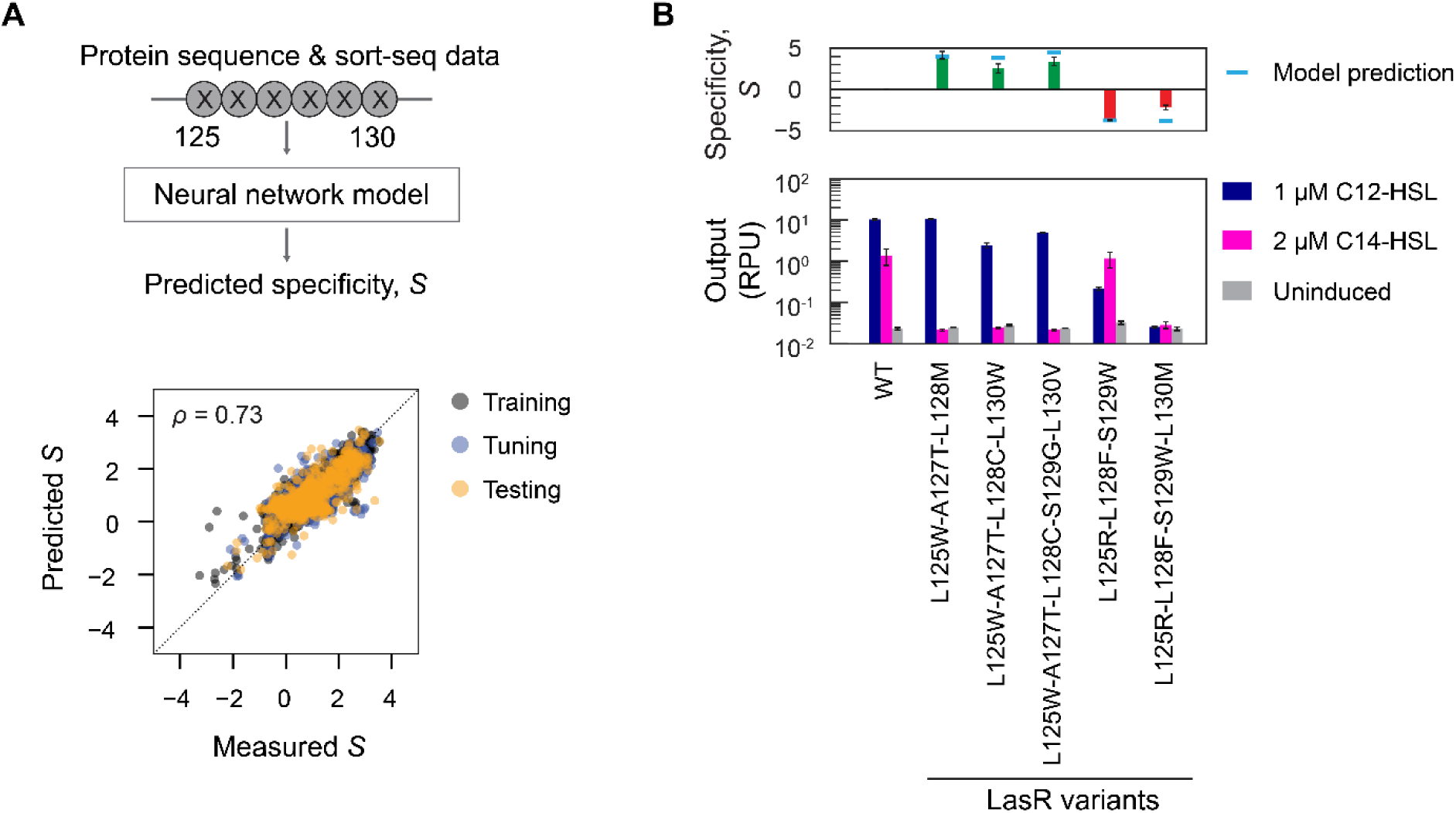
Machine learning model for protein sequence-function for LasR ligand specificity. **(A)** Using the sort-seq data for 9,140 protein variants, a machine learning model was developed to predict the ligand specificity (*S*) from the LasR protein sequence input. The convolutional neural network model for *S* was constructed using a published platform that used supervised deep learning (94). The sort-seq data was randomly split into a model training set (gray, 60% of variants) and tuning set (blue, 20% of variants) to generate the model. The model was evaluated using a testing dataset (orange, 20% of variants), and the Pearson correlation coefficient (*ρ*) was determined. Dotted line shows the 1:1 relationship. Based on this convolutional neural network model, we applied a random-restart hill climbing searching algorithm to design LasR variants with desired specificities. Convolutional neural network model for negative of specificity (−1 × *S*) in Supplementary Figure S18 was utilized for prediction of LasR designs with reversed specificity using the same searching algorithm. **(B)** From the model, the protein sequences with the maximum *S* and containing three, four, or five mutations were identified. To evaluate whether preferential activation by C14-HSL could be predicted, protein sequences with minimum S and containing three or four mutations were also identified. The corresponding LasR sensors were constructed and then assayed individually without inducer (light grey), with 1 µM 3OC12-HSL (C12-HSL, dark blue) or 2 µM 3OHC14-HSL (C14-HSL, magenta) on 3 separate days. The cell fluorescence was measured by flow cytometry for a population of at least 10,000 cells, and output was converted to RPU (Methods). The specificity of each sensor variant in each day’s experiment was calculated (Methods) and compared to the model prediction (light blue line). Bars indicate the mean ± s.d. (*n* = 3).

### Mapping the landscape for mutational epistasis

Mutational epistasis, in which the phenotypic effect of a mutation can widely vary depending on the genetic background that it is introduced into, can make a protein sequence-function landscape rugged (70, 91). From our combinatorial saturation mutagenesis assays, we next sought to examine epistatic interactions of mutations in LasR for the variable region (position 125 –130). Various models can be used to evaluate whether mutations display epistasis (90, 115, 116). Here, we applied the relative epistasis model of Khan et al (90). Pairwise epistasis was quantified as the difference between the specificity of a double mutation variant and the sum of the specificity of the two corresponding single mutation variants (Methods).

Among the exhaustive set of double mutation variants, both negative and positive pairwise epistasis were identified, yet the large majority (83%) had a negative epistatic effect (Figure 6 and Supplementary Figure S19). Such pervasive negative epistasis may be due to the indirect or direct physical interactions caused by the spatial proximity of the amino acid residues in the β sheet (117, 118). Prevalent negative epistasis in β sheets has been identified in other proteins, such as TEM-1 β-lactamase (119). Notably, eleven mutations (L125R, L125K, L125W, A127C, A127V, A127L, A127M, L128F, S129G, S129W, L130F) exhibited positive epistasis with many other mutations, yet the specificity score for each individual mutation was less than zero (Supplementary Table S1). One of these mutations (L130F) was reported to enhance the stability of LasR (79). Some LasR variants with improved specificity emerged out of positive epistasis. Among the selected ten LasR double mutation variants with the highest specificity from the library, six variants had positive epistasis (Supplementary Table S2). From a practical point of view, those variants with improved specificity would not have been identified if residues at those positions were fixed during an earlier round of mutagenesis, and this supports the use of combinatorial mutagenesis.

**Figure 6.**
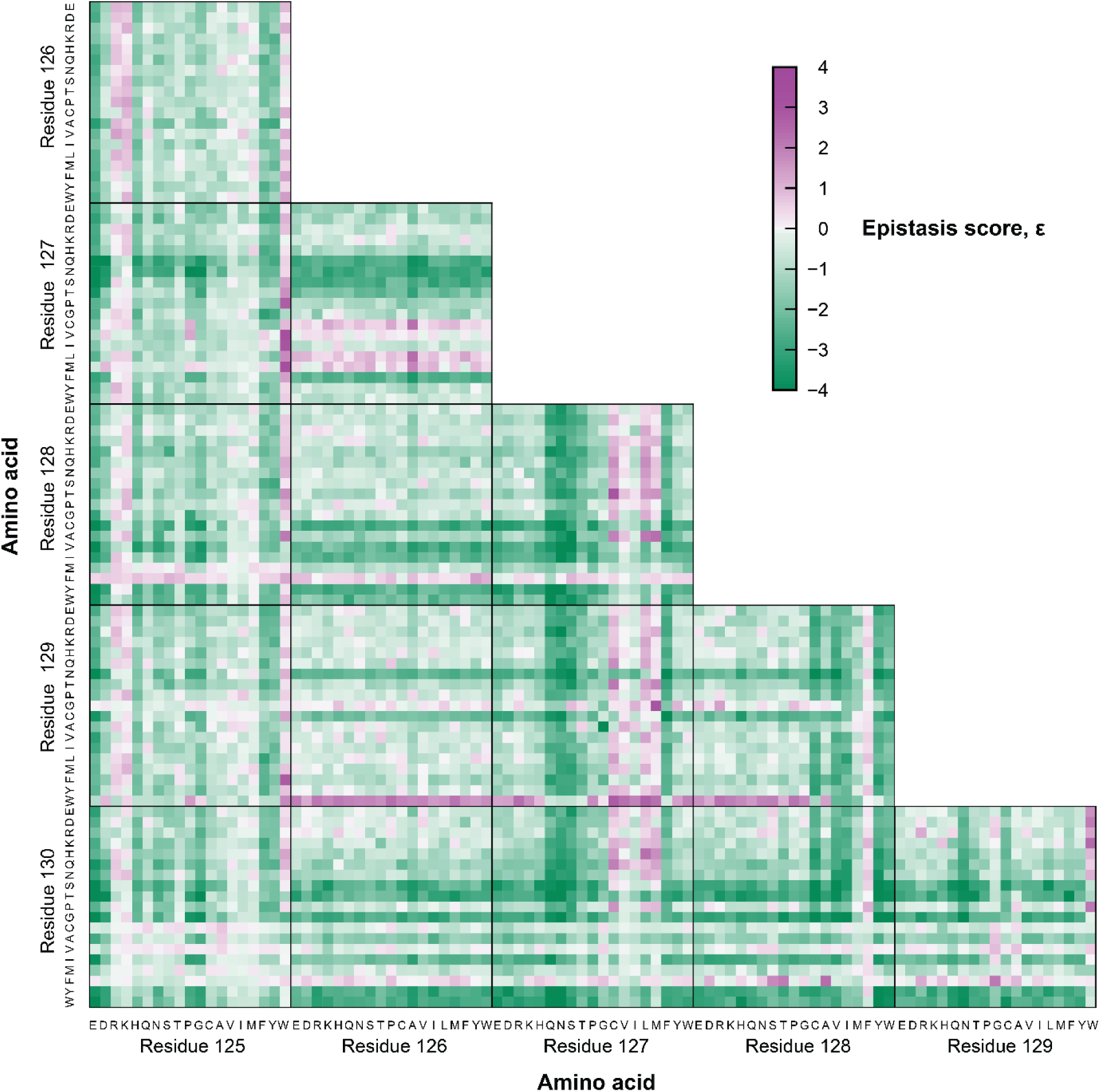
Pairwise mutational epistasis of LasR signal specificity. Pairwise epistasis scores (*ε*), defined as the non-additive effect of two mutations on sensor specificity (*S*), were calculated from the sort-seq dataset for LasR double mutation variants (Methods). Each heat map shows the epistasis for each combination of amino acids mutations (19 with exception of wildtype) at each position 125-130. Positive epistasis (purple) and negative epistasis (green) are colored. Heat maps are arrayed to show all combinations of amino acid mutations at positions 125 – 130. Amino acids are ordered based on their properties, and identical ordering is used in all heat maps after the removal of the wildtype amino acid for each position, respectively.

We also quantified the third-order epistasis in the background of S129N for our set of LasR triple mutation variants. Again, we used the relative epistasis model and calculated the epistasis as the difference between the pairwise epistasis score in the presence of S129N background (third mutation) and in its absence (Methods) (90). Pairwise epistasis in the S129N genetic background was dominantly positive (Supplementary Figure S20) and led to pervasive third-order positive epistasis (83.3% of protein sequences) (Supplementary Figure S21). However, those regions with positive epistasis in the third-order epistatic heatmap mostly contain variants that have low signal activation to the HSLs (Supplementary Figure S12, S13). Among the selected ten LasR triple mutation variants with greatest specificity in the library, all demonstrated positive third-order epistasis in the background of S129N, although most were relatively weak (Supplementary Table S3).

### Investigation of proposed trade-off between sensitivity and specificity

Previous studies have shown that increasing specificity of LuxR-type quorum sensing regulators can come at the expense of decreasing its sensitivity, meaning that as the regulator better discriminates the cognate ligand, it requires a higher concentration of the ligand to reach the same level of activation (79, 81). Therefore, we next investigated whether this trade-off between sensitivity and specificity was observed for our set of LasR variants with increased C12-HSL specificity. Among the sensor variants identified above (Figure 3), we selected variants with improved C12-HSL specificity whose output to 2 μM C14-HSL was less than 0.1 RPU (8 single, 10 double, and 9 triple mutation variants) and characterized the sensor response to C12-HSL for each (Figure 7A and Supplementary Figure S22)

**Figure 7.**
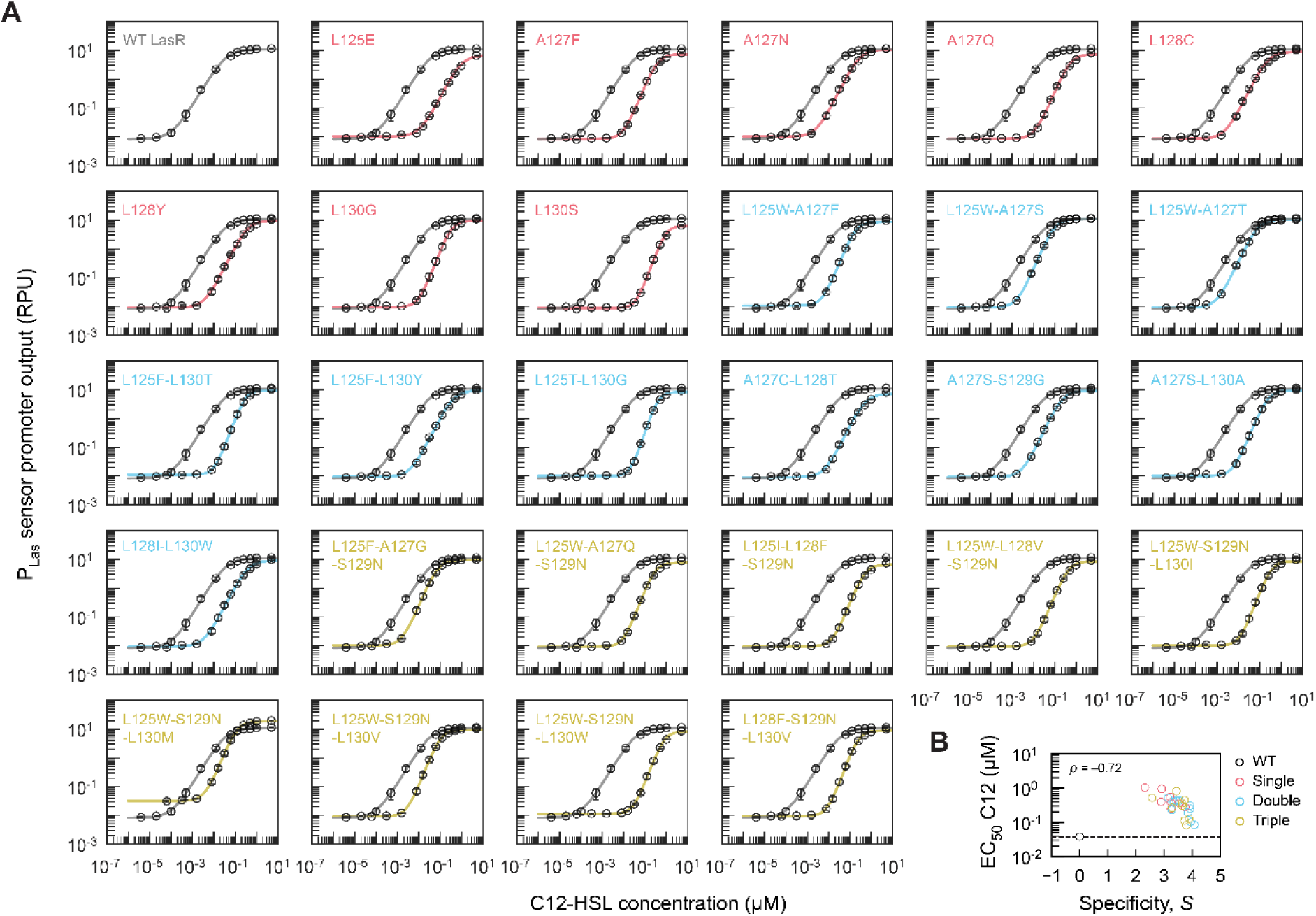
Response curves of engineered biosensors with LasR variants to investigate the relationship between sensitivity and specificity. **(A)** Twenty-seven LasR variants with single (red), double (blue), or triple (yellow) mutations were selected (from variants in Figure 3) to assay their dose-response curves to 3OC12-HSL (C12-HSL) relative to the sensor containing wildtype (WT) LasR. Sensor characterization assays were performed with addition of 0 – 5 μM C12-HSL on three separate days. The geometric mean fluorescence of at least 10,000 cells was measured by flow cytometry, and output was converted to RPU. Markers indicate the mean ± s.d. (*n* = 3 biological replicates). Representative histograms are shown in Supplementary Figure S22. Each solid line indicates the sensor response function, which is the Hill equation fitted by least squares regression to the experimental measurements. The wildtype LasR sensor (gray line) is shown in each panel for comparison. **(B)** To assess the relationship between sensitivity and ligand specificity for these sensor variants, the half maximal effective concentration (EC_50_) of C12-HSL was determined from the fitted response function for each sensor and is plotted as a function of the average specificity *S,* as measured in Figure 3. The Pearson correlation coefficient was determined for all 27 LasR variants and does not include the wildtype. The dashed line indicates EC_50_ of the wildtype LasR sensor.

(Methods). The measurements from the dose-response experiments were fit to the Hill equation for cooperative ligand binding to determine the sensor response function, and this function describes the relationship between the ligand concentration and the sensor promoter output (P_Las_) (Supplementary Table S4). Compared to the wildtype LasR sensor, all assayed sensor variants were less sensitive to C12-HSL. The ligand concentration for half-maximal sensor output (EC_50_) for the least sensitive variant (1.04 ± 0.08 µM C12-HSL for L125E) was 28 times greater than the EC_50_ of the wildtype LasR sensor (0.037 ± 0.004 µM C12-HSL for wildtype). However, two LasR variants had EC_50_ values less than 2.3-fold increase compared to the wildtype (0.086 ± 0.009 µM C12-HSL for L125W-A127T and 0.083 ± 0.008 µM C12-HSL for L125F-A127G-S129N). Interestingly, one variant (the L125W-S129N-L130M) had increased basal promoter activity (*p* = 0.0093), despite mutations being located outside the helix-turn-helix DNA binding domain, yet similar effects have been observed for engineered LacI as well (120). Overall, EC_50_ values showed a strong negative correlation with specificity for the characterized set of 28 LasR sensor variants (Pearson coefficient, *ρ* = −0.73) (Figure 7B). More sensitive (i.e. having smaller EC_50_) engineered LasR variants in this set had greater specificity, suggesting that some amount of improved ligand discrimination can be afforded without requiring a loss of sensitivity.

## DISCUSSION

The specificity of quorum sensing regulators is important for programming intercellular signaling in microbial consortia. Here, we demonstrated a pooled sort-seq approach that enables the estimation of specificity by quantifying a quorum sensor’s response to cognate and non-cognate HSLs for 9,486 variants in parallel. By combining a combinatorial saturation mutagenesis library and high-throughput sort-seq assay, we identified engineered sensors that eliminated problematic signal crosstalk along with providing insights on the sequence-function relationship for ligand specificity of this widely used quorum sensing regulator, LasR.

The hit rate of LasR variants with high C12-HSL specificity (5.7%, 559 variants) in our study indicates the functional plasticity of this protein. Functional plasticity for the specificity of LasR homologs has been reported (81), and therefore, it is not entirely surprising that we identified remarkable functional plasticity in the short LasR β5 sheet (L125-L130). Notably, certain mutations at non-contacting residues (≥ 5 Å) in the vicinity of the ligand binding pocket (residue L128 and L130) are beneficial for LasR specificity, expanding the repertoire of LasR variants with improved target specificity. Previous studies have highlighted the involvement of residue L130 in ligand selection by LasR (79, 80). Additionally, residues outside of the β5 sheet have been shown to be essential for LasR specificity through covariation analysis (80). We conducted site-directed mutagenesis at several of these essential residues (i.e. LasR G38, V76, and S91) and identified additional beneficial mutations (Supplementary Figure S23). Future studies could further expand the residues to target for improved LasR specificity beyond the β5 sheet.

Typically, evolutionarily conserved residues of LasR homologs are less tolerant to mutations as they are essential for function (42, 81, 84, 121). Residue conservation can be assessed from sequence alignments of functionally related protein homologs (122–124). Previous studies based on a small set of sequence alignments of LasR homologs (≤ 11) revealed that LasR G126 has lower evolutionary conservation compared to S129 (41, 125). To comprehensively assess residue conservation within our targeted region, we quantified amino acid frequencies at each position from an alignment of over 3,000 LasR homologs (80), uncovering lower G126 frequency relative to S129 in this larger set (Figure 25). Nevertheless, in our study, all mutations at position 126 abolished sensing of both HSLs (i.e. C12-HSL and C14-HSL). Conversely, four single amino acid substitutions (asparagine, glycine, cystine, or alanine) maintained 50% of the wildtype LasR activation by C12-HSL at position 129. Our dataset experimentally validates the importance of G126 for LasR sensing function, confirming a previous inference from LasR’s codon usage frequency in clinical isolates of *P. aeruginosa* (126). Other than residue G126, our study corroborates correlations between the mutational tolerance of residues within the β5 sheet of LasR and their level of evolutionary conservation in LasR homologs (41, 125) (Supplementary Figure S24). Notably, position 125, exhibiting the highest tolerance for mutations within our dataset, also has the greatest diversity of amino acids among natural LasR homologs (Supplementary Figure S24). Similar correlations have been observed for a wide range of proteins (127, 128) and could be harnessed to reduce the size of a designed mutagenesis library for functional screening of a protein, as demonstrated in previous studies (76, 110).

The presented pooled sort-seq method offers a promising approach for effectively engineering the specificity of LuxR-type quorum sensors. Whereas natural bacterial quorum sensing is believed to utilize promiscuity of LuxR-type regulators to mediate interspecies interactions via signal crosstalk (e.g. eavesdropping), protein engineering offers the opportunity to leverage their mutational tolerance to obtain optimal ligand specificity for a given application (49, 129–131). We selected the residues for mutagenesis based on the protein crystal structure of LasR and known ligand-protein interactions, which likely contributed to the success of this approach. For LuxR-type regulators without known crystal structures, the mutagenesis sites may be alternatively selected based on covariation analysis of naturally evolved protein sequences or perhaps by inference from this LasR library dataset and protein sequence homology. Our study’s use of a combinatorial saturation mutagenesis library allowed for the identification of many LasR variants with improved C12-HSL specificity, while also systematically elucidating the relationship between ligand specificity and the sequence of the targeted region of the β5 sheet. Engineering the specificity of quorum sensors has the potential to provide a toolset for designing and prescribing cell-cell signaling interactions to bring about multicellular bacterial processes and advance the applications of engineered bacterial consortia.

## MATERIAL AND METHODS

### Reagents and strains

*E. coli* NEB 10-beta from New England Biolabs (NEB) was used for sensor assays. *E. coli* NEB 5-alpha (NEB) was used for standard cloning of other plasmids. All sort-seq and individual assays were performed in M9 minimal media (Sigma-Aldrich; 6.78 g/L Na_2_HPO_4_, 3.0 g/L KH_2_PO_4_, 1.0 g/L NH_4_Cl, 0.5 g/L NaCl final concentration) with 0.34 g/L thiamine hydrochloride (Sigma-Aldrich), 0.2% w/v casamino acids (Acros), 2 mM MgSO_4_ (Sigma-Aldrich), 0.1 mM CaCl_2_ (Sigma-Aldrich), and 0.4% w/v D-glucose (Sigma-Aldrich). Luria-Bertani (LB) Miller media (Fisher BioReagents) was used for cloning and plasmid propagation. The antibiotic used for sensor plasmid selection was kanamycin (50 μg/ml, GoldBio). The inducers used for sensors were *n*-(3-oxododecanoyl)-L-homoserine lactone (C12-HSL, Sigma-Aldrich, #O9139) and *n*-(3-hydroxytetradecanoyl)-DL-homoserine lactone (C14-HSL, Sigma-Aldrich, #51481). For blue-white screening, 40 μl of 0.1 M isopropyl β-D-1-thiogalactopyranoside (IPTG, GoldBio) and 120 μl of 20 mg/ml 5-bromo-4-chloro-3-indolyl β-D-galactopyranoside (X-gal, GoldBio) were overlaid on LB agar plates. All primers and oligos were ordered from Integrated DNA Technologies (IDT) dried and resuspended in 10 mM Tris buffer (pH=8, Fisher Scientific).

### Design of oligos for LasR mutagenesis

The LasR mutations were encoded in the oligos designed by a customized Python script (https://github.com/AndrewsLabSynBio/directed_mutagenesis_library). All oligos in the pool have sequences for PCR amplification, flanking BsaI recognition sites and the resulting overhangs for one-pot Type IIS DNA assembly surrounding the LasR variable region (Supplementary Figure S3A). The orthogonality of the two DNA linker sequences was examined by a previously published Python script (85). The LasR variable region contains 18 nucleotides encoding the codons for residues 125-130 of LasR. Oligo designs were generated for saturation mutagenesis of all single and double mutations. A subset of all possible triple mutations was designed. The triple mutations were designed by including the S129N mutation in all designs and mutating two other residues simultaneously. The Python script by default selects codons with the highest usage frequency for the input organism (*E. coli* NEB 10-Beta in this study), unless specified problematic sequences are introduced (Supplementary Figure S3B). For example in this work, if a BsaI recognition site would be introduced by selecting the highest frequency codon, the codon with the next highest frequency was chosen instead. The Python script generates combinatorial saturation mutagenesis libraries for up to 3 amino acid mutations. An option to include an oligo containing the wildtype DNA sequence and eliminate repeated oligo designs is provided in the script. A combinatorial saturation mutagenesis library can be generated for a specified protein sequence and input organism. The codon frequency for the organism can be input in a CSV file or predicted based on a FASTA file of coding sequences in the organism. The positions to be mutated and amino acid preferences can be specified by the user. The flowchart of the customed Python script is detailed in Supplementary Figure S25. Our 12400-member oligo pool contained 20 copies of each single mutation design as an added measure to ensure full representation of all single mutation variants. The oligo pool in this work was ordered as a 12K chip from GenScript (NJ, USA).

### Library preparation using one-pot Type IIS assembly

The LasR sensor backbone was built by removing the BsaI recognition site on plasmid pMZ103-rbs1 (Supplementary Figure S3C) and by inserting the LacZ gene into the LasR variation region (Supplementary Figure S3D). The BsaI recognition site on pMZ3-rbs1 was removed by introducing a mutation in BsaI recognition site via PCRs. Two purified PCR products were joined by BbsI Type IIS Assembly reaction using 5 fmol of each purified PCR fragments, 0.5 µl of 10X T4 ligase buffer, 250 U T4 ligase (2,000,000 U/ml, NEB, M0202T), 5 U BbsI (NEB, R0539L) and nuclease-free water for a total of 5 µl reaction mix. The reaction mix was incubated in a thermocycler with the following protocol: alternating steps of 37 °C for 2 min and 16 °C for 2 min for 36 cycles, followed by 50 °C for 30 min and deactivation at 80 °C for 20 min. Transformation was performed by adding 2 µl reaction mix into 5 µl chemical competent *E. coli* NEB 5-alpha cells and following the manufactured recommended protocols. Transformations were plated on LB agar with kanamycin. A colony from the overnight plate was selected for an overnight liquid culture, a plasmid DNA miniprep (Qiagen, 27106), and Sanger sequencing (Azenta) to verify the DNA sequence. In the subsequent assembly to construct LasR sensor backbone, the LacZ gene was then inserted in place of the LasR mutated region (LasR L125 – L130) to evaluate the efficiency of library cloning using blue-white screening. The assembly reaction contained 5 fmol of a purified PCR fragment of LasR sensor backbone, 10 fmol of a purified PCR product of the LacZ gene, 0.5 µl of 10X T4 ligase buffer, 250 U T4 ligase, 5 U BbsI and nuclease-free water for a total of 5 µl reaction mix. The reaction mix was incubated in a thermocycler and transformed into *E. coli* NEB 5-alpha using the protocols above. This resulting sensor backbone was sequenced by Sanger sequencing.

The single stranded oligos in the pool were amplified using KAPA HiFi HotStart DNA polymerase (KAPA BIOSYSTEMS). The PCR reaction contained 1 µl of oligo pool resuspended in TE buffer (20.87 ng/µl), 0.75 µl of 10 µM Amp-Fw primer (TCAGTCCATCCCACCTTGCC), 0.75 µl of 10 µM Amp-Rv primer (ACCACACAGCCATAGAGTCG), 0.75 µl of 10 mM dNTP, 5 µl of 5X KAPA reaction buffer, 0.5 U KAPA polymerase and nuclease-free water for a total of 25 µl reaction mix. Two reaction mixes were incubated in a thermocycler with the following protocol: 95 °C for 3 min, 98 °C for 20 s, 68.8 °C for 15 s, 72 °C for 15 s, repeating the last three steps for 6 cycles, and 72 °C for 1 min. The amplified oligo pool was then purified using GenCatch^TM^ PCR Cleanup Kit (Epoch Life Science) and eluted in Tris buffer (pH = 8).

The LasR sensor library was constructed using one-pot BsaI Type IIS Assembly reactions, containing 50 fmol of LasR sensor backbone, 80 fmol of amplified oligo pool, 0.5 µl of 10X T4 ligase reaction buffer, 5 U BsaI-HFv2 (NEB, R3733L), 250 U T4-ligase and nuclease-free water for a total of 5 µl reaction mix. A negative control with only 50 fmol of LasR sensor backbone was set up simultaneously. The reaction mixes were incubated at 37 °C for 5 h, 30 °C for 30 min and then 80 °C for 20 min. Ultra electrocompetent *E. coli* NEB10-beta was prepared as per Supplementary Note S1. Transformations were performed by adding 4 µl of reaction mix into 100 µl of electrocompetent *E. coli* NEB 10-beta in 1 mm electroporation cuvettes (Fisherbrand, #FB101) using a BIORAD MicroPulser (Setting: Ec1, 1.8 kV) and subsequent mixing with 500 µl of prewarmed SOC. Next, 250 µl of the resuspended cells were transferred to two aliquots of 750 µl prewarmed SOC and incubated at 37 °C, 250 rpm for 1 h. Then 90 µl of the recovered cell media were plated on each LB agar plate (2.25% agar) with kanamycin using a cell spreader and an inoculating turn table. Additionally, 5 µl of recovered cell media was plated on LB agar plates containing IPTG and X-gal for the blue-white screening and coverage estimation. Two transformations were carried out to ensure 40X coverage of the library (40 X 12,400 = 480,000 colonies). The reaction mix of the negative control was also electroporated to *E. coli* NEB 10-beta as described above and were plated on a LB agar plate containing X-gal and IPTG.

The agar plates were incubated for 12 h until the formation of singular colonies without merging to avoid the competition of colony growth. The total number of colonies was estimated to be 1,025,000 (82.7-fold of 12,400 oligo designs). The plate of the negative control had 1,103 colonies (769 blue and 334 white). The estimated error rate of the library was 0.22%. The colonies were harvested from all plates using a cell lifter, resuspended into LB and vortexed vigorously for uniform suspension. The suspension was centrifuged at 3,900 rpm for 10 min, the supernatant was discarded, and the cell pellet was resuspended in 20 ml fresh LB. An equal volume of 50% v/v glycerol was added to make aliquots of glycerol stocks and stored at −70 °C freezer. Plasmids and genetic part sequences used in this work are listed in Supplementary Table S5 and S6, respectively.

### Pooled sensor characterization by FACS

The entire LasR sensor library was sorted into four bins based on the fluorescence response of each variant to cognate HSL (C12-HSL) or non-cognate HSL (C14-HSL) using Florescence-Activated Cell Sorting (FACS). Two 1-ml glycerol stocks of library aliquots and a 1-ml LasR wild-type (WT) sensor glycerol stock were thawed on ice and inoculated in 29 ml of M9 supplemented with 50 μg/ml kanamycin in 250-ml flasks. To convert the YFP expression to RPU, 0.2-ml glycerol stock of wild-type *E.coli* NEB10-beta and the RPU standard strain (pAN1717) were thawed and added to 5.8 ml of M9 and M9 supplemented with 50 μg/ml kanamycin, respectively. After 3 h incubation at 37 °C with shaking at 250 rpm (New Brunswick Innova 44 shaker), the OD_600_ of all library aliquots/strains were measured (NanoDrop OneC, Thermo Scientific). The cell media of LasR WT sensor was spiked into the library as a reference to 1% OD_600_ of the LasR sensor library. The LasR sensor library was then diluted to an OD_600_ of 0.05 and induced with either 1 µM 3OC12-HSL or 2 µM 3OHC14-HSL in a sterile 96-well U bottom plate and incubated for 5 h at 37 °C with shaking at 1000 rpm (ELMI TRMS-04 DTS-4 thermostat microplate shaker). Each library sample was then diluted using sterilized PBS and sorted by BD FACSAria Fusion Sorter into 4 separate bins. Two aliquots of LasR sensor library under two different sorting conditions yielded 16 total bins. Bin 1 contains sensors with background fluorescence based on the fluorescence level of wild-type *E. coli* NEB 10-Beta. Remaining cells in the library with higher fluorescence were separated into bin 2, bin 3 and bin 4 with roughly even proportions (424 V of FSC, 430 V of SSC, 493 V of FITC, and a threshold of 1500 au for FSC). At least 3.5 million cells were sorted per sample (Supplementary Figure S3E). Sorted cells were then cultured in LB supplemented with 50 μg/ml kanamycin for 7 h to amplify cells. The LasR sensor plasmids from the 16 bins were then extracted. To determine the distribution of designs in the constructed LasR sensor library, two unsorted library aliquots were directly miniprepped from 1ml glycerol stocks.

### Next generation sequencing

Each bin was sequenced to map the fluorescence level to the LasR design. The unsorted library was sequenced to determine LasR design distribution within constructed LasR sensor library. The LasR variable region for each sample was amplified with a set of custom designed primers (Supplementary Table S7) that varied the DNA sequence at the 5’ end of the amplicon to increase the nucleotide diversity of amplicons, which aids Illumina sequencing of low nucleotide diversity amplicons. PCR reactions were performed using KAPA HiFi HotStart DNA polymerase for 22 cycles (98 °C for 20 s, 68 °C for 15 s, and 72 °C for 15 s). The PCR products were purified and eluted in 0.1 X TE buffer. The concentration of purified PCR products was measured using a Nanodrop for amplicon library preparation. The amplicon library, containing adaptor sequences for Illumina NextSeq500, a unique barcode for each sample, and the PCR product, was created using the NEBNext Ultra II DNA library prep kit for Illumina (NEB #7103S) with index primer set 1 (NEB #7335S) and set 2 (NEB #7500S) according to the manufacturer’s protocol with minor modifications as noted below. A quantity of 600 ng of PCR product was used as the starting material, and clean-up of adaptor-ligated DNA (with 0.8X resuspended beads) instead of size selection was performed because the length of the PCR amplicon was uniform (113 bp). Four PCR cycles were chosen for PCR enrichment of adaptor-ligated DNA. The resulting amplicon libraries were purified using NEBNext® Sample Purification Beads in the kit and a magnetic stand (EpiMag HT 96-Well Magnetic Separator, Epigentek). For pooling, the DNA concentration of each sample was measured by Qubit 3.0 fluorometer. The relative amount of each barcoded DNA sample was pooled according to the number of cells in the sample (i.e. cell number in each bin) with the ratios listed in Supplementary Table S8. The quality of the pooled amplicon library was assayed on an Agilent 2100 Bioanalyzer prior to sequencing.

The pooled DNA sequencing library was sequenced on an Illumina NextSeq500 with a 50% PhiX spike-in to increase the diversity of sequence and sequence quality given the regions of low nucleotide diversity for our amplicons. A NextSeq500 Mid-150 cycle kit was used for pair-end read (2 X 75 base pairs). A total of 8,500,000,000 bp (8.5 Gbp) of NGS data were collected, yielding 5.6X read coverage on average for the cells in each bin.

### Next generation sequencing data analysis

The FASTQ file for each sample was demultiplexed on Illumina platform. The perfect-match read counts for each design in each sample were quantified using a custom Python script that counts the reads that perfectly match the reference sequences or reverse complement thereof. The reference sequences contain 9-bp constant upstream sequence (CGCGGCGAA), 18-bp DNA sequence of LasR designs, and 7-bp constant downstream sequence (AGCGTGG).

The assessment of design y’s frequency across various samples served to evaluate the agreement among unsorted samples and the reproducibility between two sorted sample replicates. The frequency was determined using the following equation:

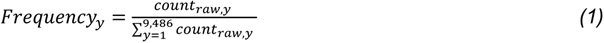

where *count_raw,y_* is the raw count or perfect-match read counts design *y*, and the denominator is the sum of raw counts of all 9,486 LasR designs in a sample.

Most LasR designs were present in multiple bins. Due to the expansive upper limit of bin 4, we chose median fluorescence as fluorescence measurement for both bin 4 and the other three bins. The inferred fluorescent output of each LasR sensor design (*FL_y_*, representing the fluorescence of design *y* in arbitrary units) of each replicate was quantified by assuming the median fluorescence of the bin for the proportion of cells contained in the corresponding bin, which was calculated by the following equation:

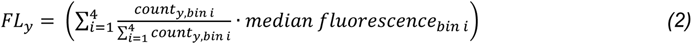

In this equation, *count_y_*_,*bin i*_ is the actual read counts of design *y* in bin *i*, and the value of *median fluorescence_bin i_* was determined during FACS.

Given that bin 1 contained more than 60% of the entire library and was stopped earlier during FACS (stopping threshold: 2,000,000 cells), the raw counts in bin 1 are lower than its actual counts. To estimate the actual cell counts in bin 1, scaling factor was calculated using the following equations:

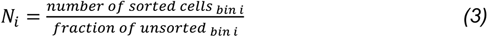

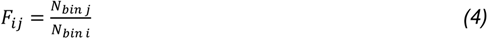

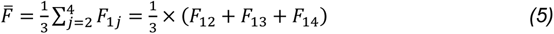

In equation (3), *N_i_* is estimated total unsorted cell counts from bin *i*. Both the number of sorted cells in bin *i* and fraction of bin *i* within unsorted library were obtained through FACS. In equation (4), *F_ij_* is the scaling factor for cell counts in bin *i* derived from bin *j*. In equation (5), *F̄* is the average scaling factor for cell counts in bin 1, derived from bin 2, 3, and 4. The calculations of *F̄* for each sorted sample are shown in Supplementary Table S9.

Additionally, the determination of the pool ratio for each sample was based on sorted cells count within each bin and then slightly adjusted to ensure enough sequencing coverage of bin 4, which had the lowest cell count. Thus, the actual counts were subsequently corrected using the coverage correction factor *f_bin i_* (Supplementary Table S8). The actual counts of LasR design *y* in bin *i* were calculated using the following equation:

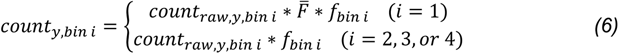

In equation (6), *count_raw_*_,*y*,*bin i*_ is the raw count of design *y* in bin *i*, *F̄* is the average scaling factor for the cell counts within bin 1, and *f_bin i_* is the coverage correction factor. Parameters for each bin are summarized in Supplementary Table S10.

Fluorescence in arbitrary units cannot be compared directly for measurements collected using different flow cytometers, such as between a cell sorter and analytical cytometer (86, 87). However, by using an insulated fluorescent reporter and standard reference plasmids (ribozyme, RBS, and gene sequence for eYFP are held constant), the cell fluorescence in arbitrary units can be converted to relative promoter units (88, 89). Therefore, here we similarly converted fluorescence in arbitrary units to relative promoter units (RPU):

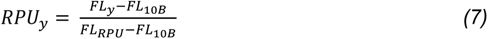

where *FL_RPU_* is the median fluorescence of the RPU strain harboring a plasmid (pAN1717) with a standard constitutive promoter and sensor characterization cassette, and *FL*_10_*_B_* is the median fluorescence of wild-type *E. coli* NEB 10-Beta (autofluorescence of cells), measured on the flow cytometer instrument for a population of at least 10,000 cells. The RPU of each variant is provided in Supplementary Spreadsheet, Tab 2.

### Specificity and epistasis score calculation

Specificity (*S*), the target function of the LasR protein in this study, was defined as the difference between the natural logarithm of C12-HSL output to C14-HSL output for the variant and wildtype LasR.

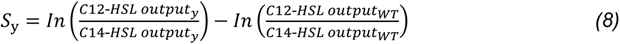

In equation (8), the C12-HSL output and C14-HSL output were the fluorescence outputs of LasR sensors containing LasR design y or WT LasR to C12-HSL and C14-HSL in RPU, respectively.

Pairwise mutational epistasis for LasR mutation variants was calculated from the average specificity as determined by the sort-seq experiments using the relative epistasis model (90). Using this model, the epistasis (*ε*) for a fitness function (*W*) for a genotype containing two mutations (*a* and *b*) is the log ratio of observed and predicted fitness values and applies the multiplicative null hypothesis for the predicted fitness [*ε* = ln(*W_ab_*) − ln(*W_a_*) − ln(*W_b_*)]. Our specificity function here serves as the fitness function and includes the log transformation as defined above. The wildtype LasR sensor was used as the genetic background reference (*S_WT_*), where its relative specificity was set to zero by definition of the function (*S_WT_ =* 0). The specificity of LasR containing two amino acid mutations (*S_ij_*) at positions *i* and *j* (e.g. *S_AB_* for amino acid A in position *i* and amino acid B in position *j*) and the specificity for LasR containing each mutation individually (e.g. *S_A_* for amino acid A in position *i* and *S_B_* for amino acid B in position *j*) were used. Each pairwise epistasis score (*ε_ij_*) for each combination of amino acid mutations in each position (*ε_AB_* for amino acid A in position *i* and amino acid B in position *j*) from the double mutation variants was calculated as follows using the average specificity of the variants (91, 92):

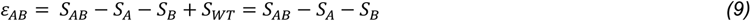

Similarly, the third-order epistasis scores were calculated for LasR variants containing three mutations using the relative epistasis model. Using this model, the third-order epistasis (*ε*) for a fitness function (*W*) for a genotype containing three mutations (*a*, *b,* and *c*) is the extent to which the pairwise epistasis of any two mutations differs in the background of the third mutation (92, 93). We only calculate the third-order epistasis in the background of S129N as our library only contained a small set of triple mutation variants. In the background of mutation C at position *k* (i.e. S129N), the pairwise epistasis score (*ε_ij|_*_k_) for each combination of amino acid mutations in each position (*ε_AB|C_* for amino acid A in position *i* and amino acid B in position *j*) from our triple mutation variants. Subsequently, we calculated the third-order epistasis score (ε_ABC_) as follows:

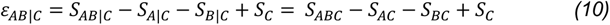

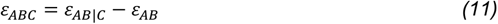

In equation (10), *S_AB|C_* is the specificity of LasR containing amino acid A in position *i* and amino acid B in position *j* in the presence of S129N, *S_A|C_* is the specificity of LasR containing amino acid A in position *i* in the presence of S129N, *S_B|C_* is the specificity of LasR containing amino acid B in the position *j* in the presence of S129N, and *S_C_* is the specificity of LasR containing S129N. In equation (11), *ε_AB|C_* is the pairwise epistasis score in the background of S129N, and ε_AB_ is the pairwise epistasis score without S129N.

### Machine learning model training and testing

Machine learning models for protein sequence-function relationships in this work were trained using the platform built by Gelman, et al. for supervised deep learning (94). We ran the scripts with the default learning rate and batch size. The input to the script contained the mutation(s) in each variant and its specificity score (Supplementary Spreadsheet, Tab 1), as determined by sort-seq experiments. In the script, the input was transformed to feature vectors containing mutation positions and physicochemical and biochemical properties of the amino acids. Some LasR designs included in our library had an identical protein sequence (such as S129N and S129N-L130L) due to codon degeneracy and the fact that we selected the highest frequency codon when designing the pool. To eliminate any overlap between different datasets, the 9,486 unique DNA sequences were reduced to 9,140 unique protein sequences by removing variants containing L125L and L130L before splitting the dataset. The LasR library dataset was randomly divided into a training, tuning, and testing dataset. The platform uses the tuning dataset to optimize the hyperparameters of the machine learning model during the run. The software can fit the following model types evaluated in this work: linear regression, fully connected network, and convolutional neural network models. We used Python v3.6 and TensorFlow to run the code in a central processing unit (CPU) environment. A script in this platform uses a random-restart hill climbing algorithm to identify variants predicted to have the highest output (function) it can find. For the convolutional neural network model, we ran the script with the objective of finding the highest specificity score and specified to identify sequences with three, four, and five mutations in the LasR L125 – L130 region. To identify variants with reversed specificity for C14-HSL (originally the non-cognate signal), we constructed a convolutional neural network model for the negative specificity (−1 x S) and otherwise carried out an identical procedure.

### Construction of individual sensor variants

Variants with improved C12-HSL specificity were selected from the LasR sensor library to validate the sort-seq assay. The individual variants were constructed in a similar way to the LasR sensor library. The single stranded DNA oligos containing the LasR variable region, enzyme recognition sites, and overhangs were ordered from IDT. These oligos were then annealed in a reaction mix containing 8.5 µl of the top strand (100 µM), 8.5 µl of the complementary bottom strand, and 2 µl of 10X ExoI buffer (NEB, B0293S). The annealing was performed in a thermocycler by gradually decreasing the temperature from 98 °C to 60 °C, with 1 decrease °C every 2 minutes. Unannealed oligos were digested by adding 10 U ExoI (NEB, M0293S) to the reaction mix, incubated at 37 °C for 1 h, and then purified using a PCR cleanup kit. BsaI Type IIS Assembly reactions were performed in 5 µl total volume containing 80 fmol of purified annealed oligos and 50 fmol of LasR sensor backbone, 0.5 µl of 10 X T4 ligase buffer, 250 U T4 ligase, 5 U BsaI-HFv2 and nuclease-free water. Next, 2 µl of reaction mix was transformed into 5 µl chemical competent *E. coli* NEB 10-beta. Transformed cells were plated on agar plates containing IPTG, X-gal, and kanamycin for blue-white screening. White colonies were selected for LB liquid culture, extraction of plasmid DNA, and Sanger sequencing to verify the DNA sequence of the construct.

### Sensor characterization assay for individual designs

A single colony of individual designs, wild-type *E. coli* NEB 10-beta, and the RPU strain were picked from overnight agar plates and inoculated in a 96-well U-bottom microtiter plate with 200 µl M9 with appropriate antibiotics. The plate was sealed with a breathable seal (AeraSeal, Sigma-Aldrich) and incubated at 37 °C, 1000 rpm for 16 h in a digital microplate shaking incubator (Elmi DTS-4). The cells were then diluted 178-fold through two serial dilutions, with 15 µl of cell culture transferred to 185 µl of fresh M9 with appropriate antibiotics in each dilution. The resulting plate was incubated at 37 °C, 1000 rpm for 3 h. After 3 h, cell culture from each well was diluted 658-fold through two serial dilutions, one transferring 15 µl of cell culture into 185 µl fresh M9 with appropriate antibiotics and the next transferring 3 µl cell culture to 145 µl M9 with antibiotics and inducers as needed. The resulting induction plate was incubated at 37 °C, 1000 rpm for 5 h. Before assaying, 2-5 µl of cells were transferred to 200 µl PBS with 2 mg/ml kanamycin (to halt cell growth) and incubated at room temperature for 30 min. For experiments assaying specificity, inducer conditions were no inducer, 1 µM C12-HSL, and 2 µM C14-HSL. For experiments characterizing the response function of LasR sensor variant, initial C12-HSL concentrations were 5 µM, and 1 µM, subsequently diluted by 2-fold for each concentration. For experiments characterizing the response function of the wildtype LasR sensor, C12-HSL concentrations were 5 µM, 1 µM, 0.5 µM, 0.25 µM, 0.125 µM, and 0.0625 µM, subsequently diluted by 5-fold for each concentration.

### Flow cytometry analysis

Cell fluorescence was assayed on a BD Accuri C6 flow cytometer equipped with a 20 mW 480 nm solid state blue laser (BD Biosciences, San Jose, CA) or BD LSR Fortessa flow cytometer (BD Biosciences, San Jose, CA) with 340 V of FSC voltage, 700 V of SSC voltage and 400 V of blue laser (488 nm) voltage. Different thresholds were applied: for the BD Accuri C6 flow cytometer (FSC-H > 25000 and SSC-H > 1000) and for the BD LSR Fortessa flow cytometer (FSC > 5000 and SSC > 2000). At least 10,000 gated events were collected for each sample. Samples were analyzed using FlowJo software. A cell gate was applied to distinguish *E. coli* cells from electronic noise or debris. The geometric mean of the fluorescence for the gated events was calculated and then converted to relative promoter units (RPU) using geometric mean fluorescence instead of median fluorescence as described in equation (7).

To determine the response function for selected sensor variants, the sensor outputs were fitted to the Hill equation. The curve fit was performed in Python using the least squares method to minimize the sum of the error between the log10-transformed measured outputs and the fitted output in RPU. The form of the Hill equation used to fit the data is as follows (95):

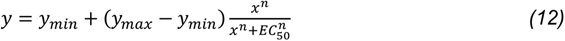

In this equation, the sensor output *y* is a function of the signal concentration (*x*) and Hill parameters are *y_min_* (minimum output), *y_max_* (maximum output), *EC_50_* (signal concentration for half maximum output), and *n* (Hill coefficient).

### Statistical analysis

The coefficient of determination (R^2^) was determined by linear least-squares regression using the scikit-learn module in Python (96). Data was analyzed using independent two-tailed Student’s *t*-tests using the SciPy package (97), and results were considered statistically significant if the *p-*value < 0.05. Pearson correlation coefficients were determined using the SciPy package. Pearson correlation coefficients for the machine learning models were determined by the published scripts that were used for modeling (94).

## DATA AVAILABILITY

The custom Python script that generated oligos designs for combinatorial saturation mutagenesis library is available on GitHub at https://github.com/AndrewsLabSynBio/directed_mutagenesis_library. All NGS files and raw counts have been deposited in the NCBI SRA database via Bioproject (PRJNA985756). Plasmids for reported sensors and LasR variants are available via Addgene (#204463-#204470).

## SUPPLEMENTARY DATA

Supplementary Data are available.

## ACKNOWLEDGEMENT

The authors thank Dr. Amy S. Burnside for assistance with cell sorting. The BD FACSAria Fusion Sorter at the UMass Amherst Flow Cytometry Core Facility was supported by a grant from the Massachusetts Life Sciences Center. The authors thank Dr. Ravi Ranjan at the Genomics Resource Laboratory Core Facility at UMass Amherst for assistance with next generation sequencing. The authors thank the Andrews group members for their feedback on experimental design and manuscript preparation.

