## Supplementary Information for "Identifying LasR quorum sensors with improved signal specificity by mapping the sequence-function landscape"

<u>**Supplementary Materials for:**</u>

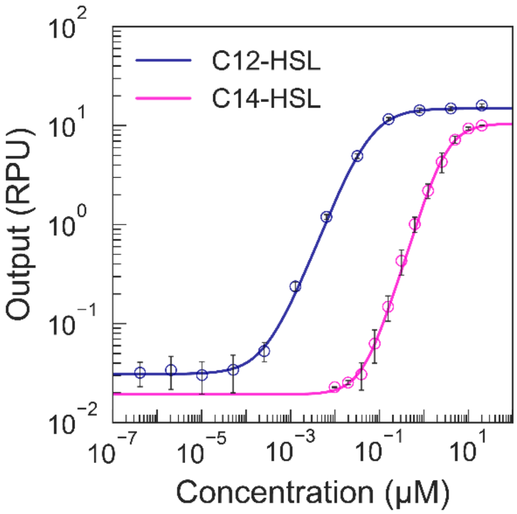


**Figure S1. Sensor response funtions for wildtype LasR with cognate C12 HSL and noncognate C14-HSL.** Sensor characterization assays were performed for the sensor construct with the wildtype (WT) LasR protein sequence with addition of 0-20 μM C12-HSL (dark blue) or C14-HSL (magenta) on three separate days. The geometric mean fluorescence of at least 10,000 cells was measured by flow cytometry, and output was converted to RPU. Markers indicate the mean ± s.d. (*n* = 3 biological replicates). Each solid line indicates the sensor response function, which is the Hill equation fitted by least squares regression to the experimental measurements.

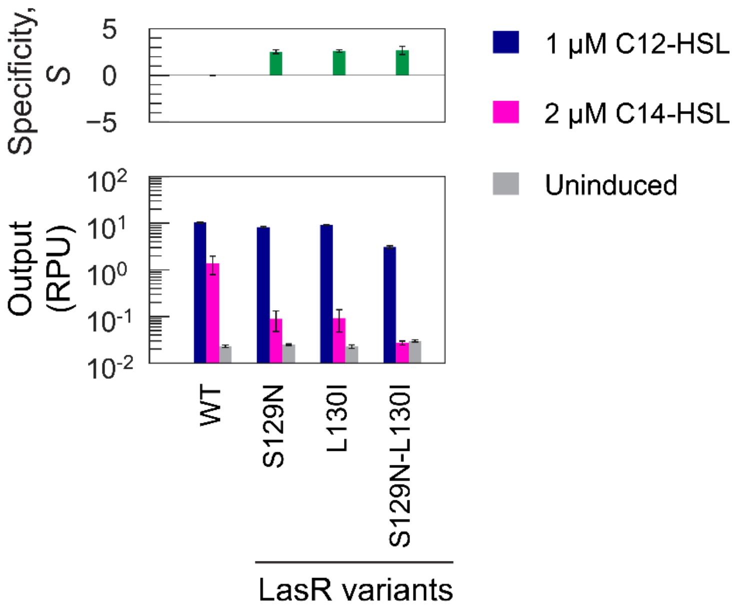


**Figure S2. LasR variants tested prior to library design.** In our selected region of LasR (L125-L130), mutations S129N and L130I were reported to reduce activation by homoserine lactones with longer acyl chain length (1, 2). We constructed sensors with each single mutation and both mutations in LasR. Sensor characterization assays were performed for these LasR variants and WT LasR on three separate days without inducer (light grey), with 1 µM C12-HSL (dark blue), or with 2 µM C14-HSL (magenta). The cell fluorescence was measured by flow cytometry. Cell fluorescence was converted to relative promoter units (RPU), and specificity (*S*) of the sensor variant on each day was calculated (Methods). Bars indicate the mean ± s.d. ($n$ = 3 biological replicates).

A

TCAGTCCATCCCACCTTGCCGGTCTCACGAANNNNNNNNNNNNNNNNNNAGCGAGAGACCCGACTCTATGGCTGTGTGGT

Sequence for PCR amplification    Overhang    **LasR variable region**    BsaI recognition site

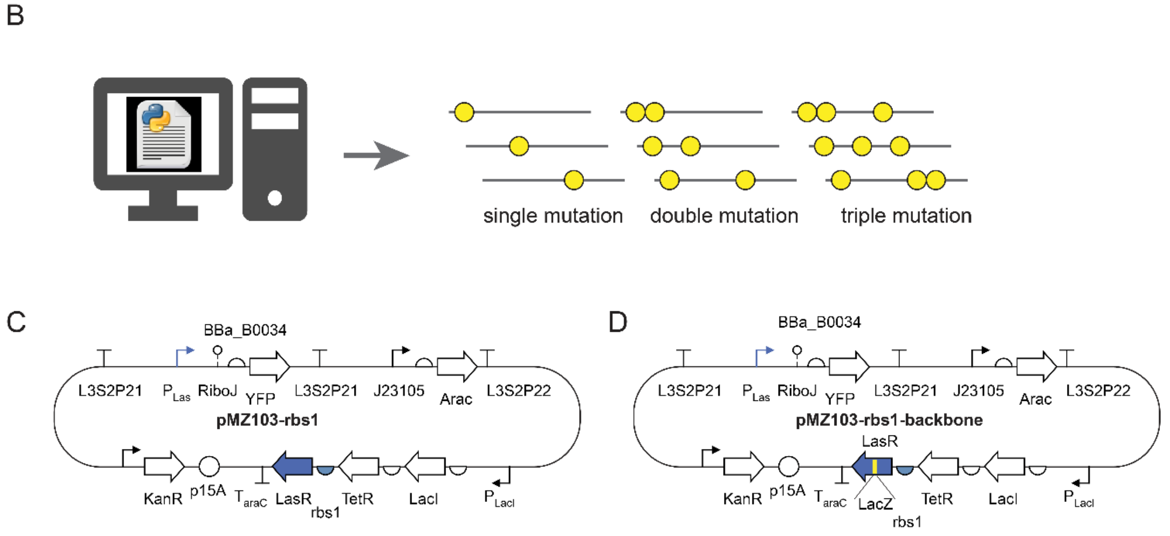


E

| replicate | condition | | Cells per bin<br>Bin 1 | Bin 2 | Bin 3 | Bin 4 |
|---|---|---|---|---|---|---|
| A | 1 μM | C12-HSL | 2,000,000 | 495,948 | 581,304 | 390,870 |
| B | 1 μM | C12-HSL | 2,000,000 | 705,053 | 786,267 | 500,752 |
| A | 2 μM | C14-HSL | 2,000,000 | 581,325 | 530,747 | 427,597 |
| B | 2 μM | C14-HSL | 2,000,000 | 565,332 | 491,980 | 405,524 |

**Figure S3. Summary of library construction and cell sorting. (A)** The sequence elements of designed oligos. The designed LasR variable region was flanked by Type IIS BsaI recognition sites and linker sequences for Type IIS assembly into the LasR sensor constructs. Outside of these sequences are amplification sequences to PCR amplify the oligo pool before DNA cloning. In the LasR variable region, N represents A, T, C, or G as specified by each design. **(B)** The oligo pool included single, double, and triple mutations and was designed using a custom Python script. The flow chart of Python script is detailed in Figure S25. **(C)** The plasmid map of the LasR sensor plasmid, pMZ103-rbs1 is shown. **(D)** The plasmid map of the LasR sensor plasmid, pMZ103-rbs1-backbone is shown. A LacZ marker (yellow) for blue-white screening was placed at the LasR 125 – 130 insertion site. **(E)** The number of cell events sorted in each bin for each of the replicate sort-seq assays with C12-HSL or C14-HSL are listed.

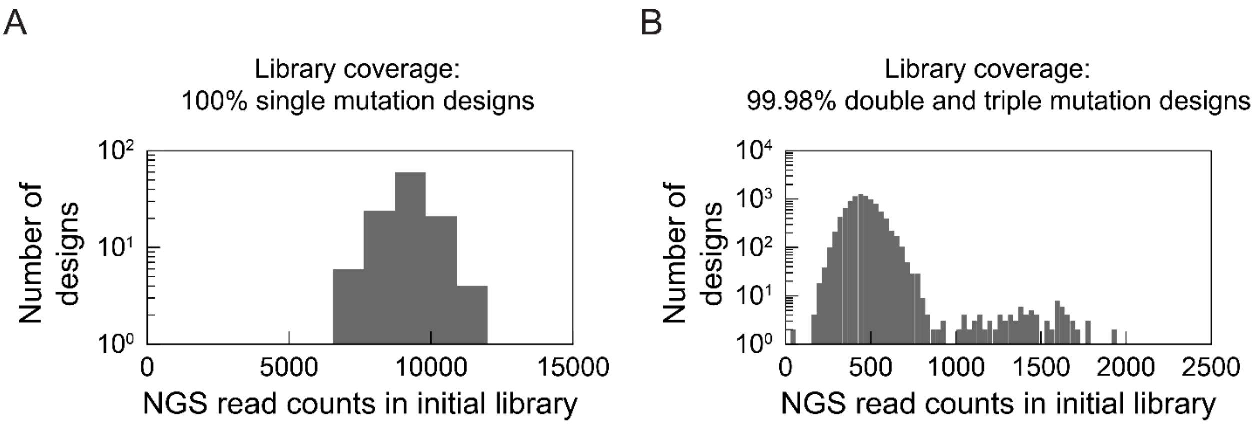


**Figure S4. Composition of the combinatorial saturation mutagenesis library analyzed by next-generation sequencing.** We sequenced two aliquots of unsorted library and quantified the perfect-match reads for each LasR design using a custom Python script. A 34-bp sequence of LasR (including the 18 bp variable region) was the reference (Methods). Read count of each design is the average of two unsorted library samples. **(A)** Perfect-match read count distribution of LasR single mutation designs in aliquots of the unsorted library. **(B)** Perfect-match read count distribution of LasR double and triple mutation designs in aliquots of unsorted library. Only two LasR designs (LasR G126D-A127G and LasR G126D-A127G-S129N) had counts lower than 100.

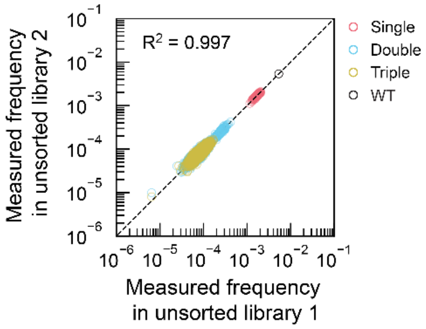


**Figure S5. Frequency of each LasR design in aliquots of the unsorted library.** The frequency of each LasR design in two unsorted libraries was determined and are compared. Markers are colored to indicate if the design contains one (red), two (blue), or three (yellow) amino acid substitutions. The wildtype LasR (WT) sequence was included in the library (black). The coefficient of determination ($R^2$) was determined by least squares regression. Dotted line indicates the 1:1 relationship.

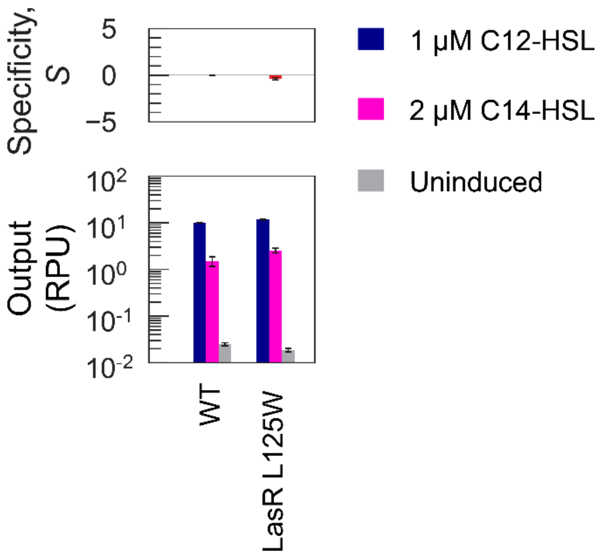


**Figure S6. Sensor characterization of LasR L125W.** The mutation L125W was identified in LasR double and triple variants with improved C12-HSL specificity, yet the sort-seq results for the variant with the L125W single mutation showed low specificity. Here, we individually assayed the sensor containing the LasR L125W without inducer (light grey), with 1 μM 3OC12-HSL (dark blue) or with 2 μM 3OHC14-HSL (magenta). The cell fluorescence was measured by flow cytometry, and the arbitrary unit was converted to standard RPU (Methods). The specificity of LasR sensor variants on each day was calculated as described in the Methods. Bars indicate the mean ± s.d ($n$ = 3 biological replicates).

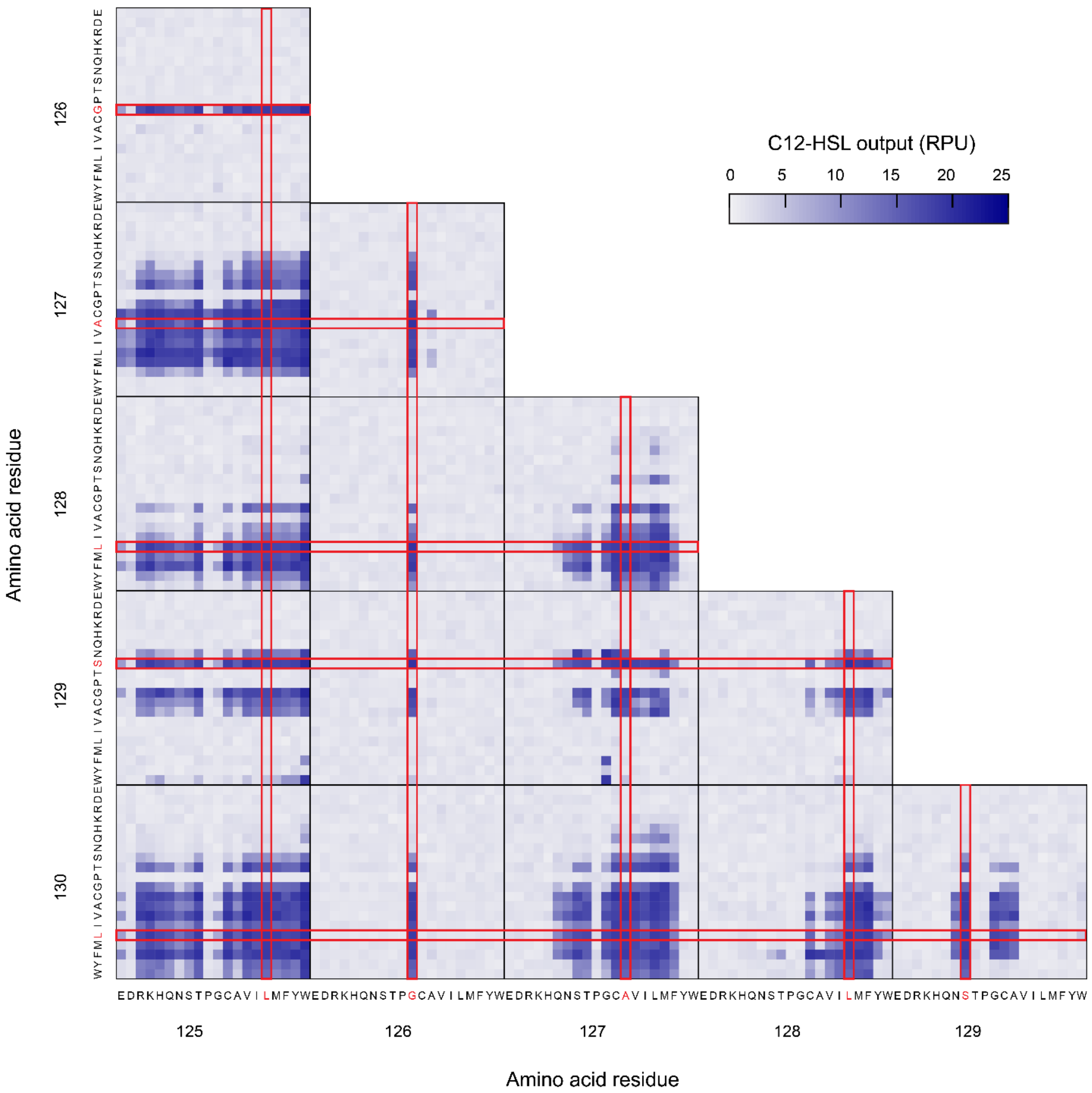


**Figure S7. Sequence-function relationship for activation by C12-HSL and LasR protein sequence (up to 2 amino acid substitutions).** An enlarged version of the left panel of Figure 4A is shown to indicate amino acids in each position using 1-letter abbreviations. Sensor output with C12-HSL was mapped against amino acids. The amino acid in wildtype LasR is shown (red). Output was determined by the average of two replicates of sort-seq using 1 µM C12-HSL (Methods).

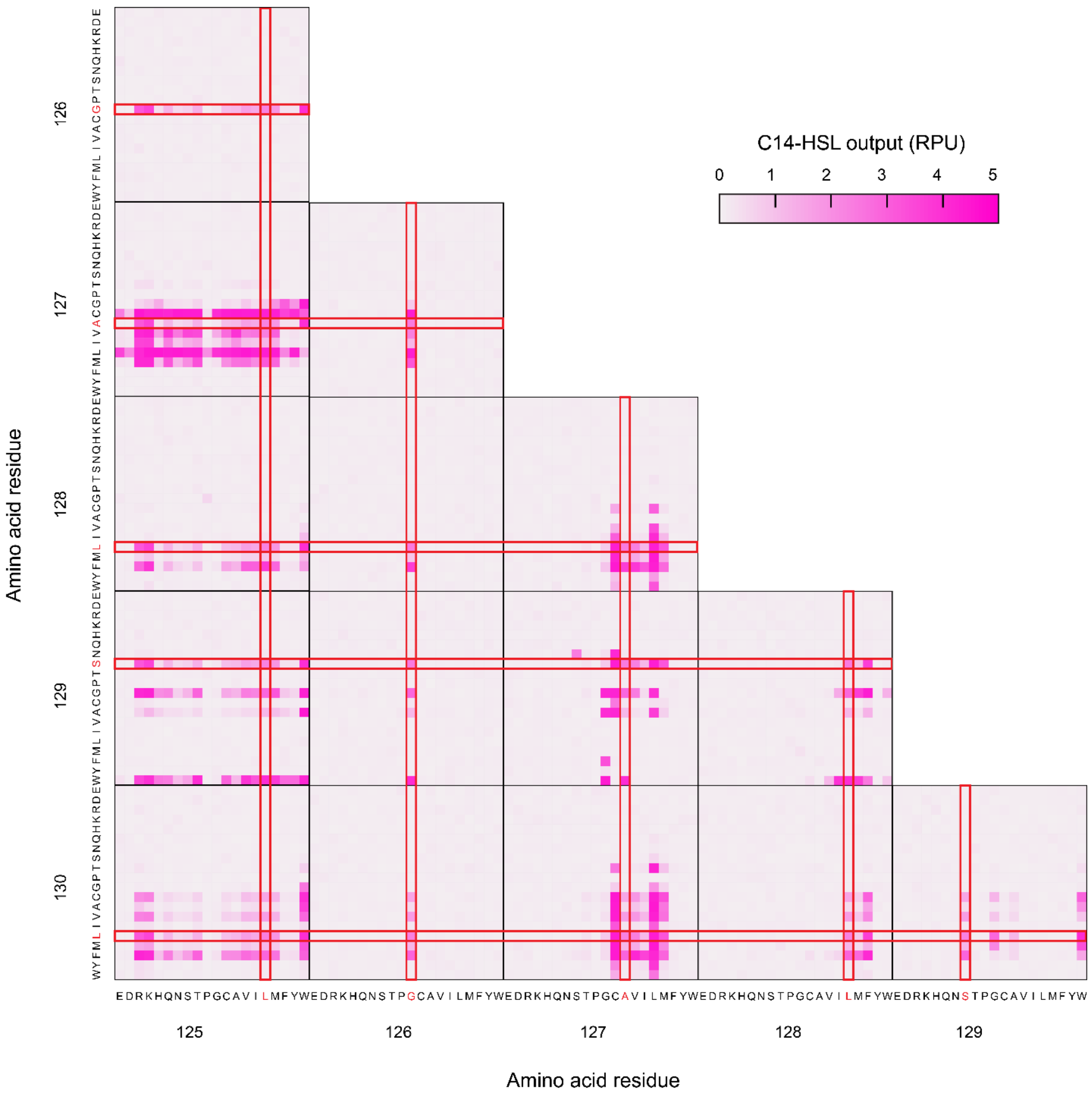


**Figure S8. Sequence-function relationship for activation by C14-HSL and LasR protein sequence (up to 2 amino acid substitutions).** An enlarged version of the middle panel of Figure 4A is shown to indicate amino acids in each position using 1-letter abbreviations. Sensor output with C14-HSL was mapped against amino acids. The amino acid in wildtype LasR is shown (red). Output was determined by the average of two replicates of sort-seq using 2 μM C14-HSL (Methods).

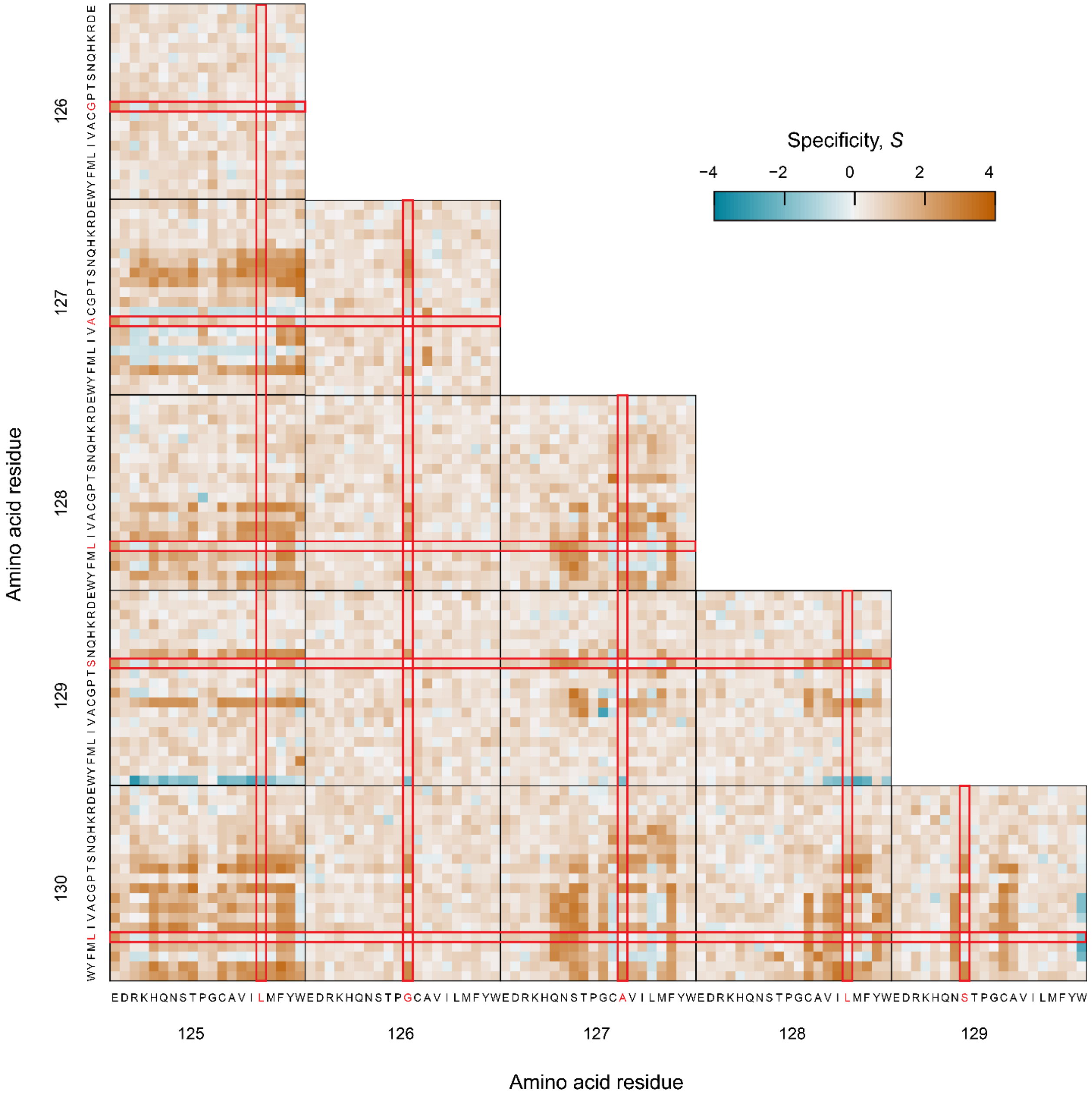


**Figure S9. Sequence-function relationship for specificity and LasR protein sequence (up to 2 amino acid substitutions).** An enlarged version of the right panel of Figure 4A is shown. Amino acid identity is listed using its 1-letter abbreviation. Specificity (*S*) was determined by the average of two replicates of sort-seq and is plotted for each amino acid in each position of the LasR variable region. The amino acid in wildtype LasR is shown (red).

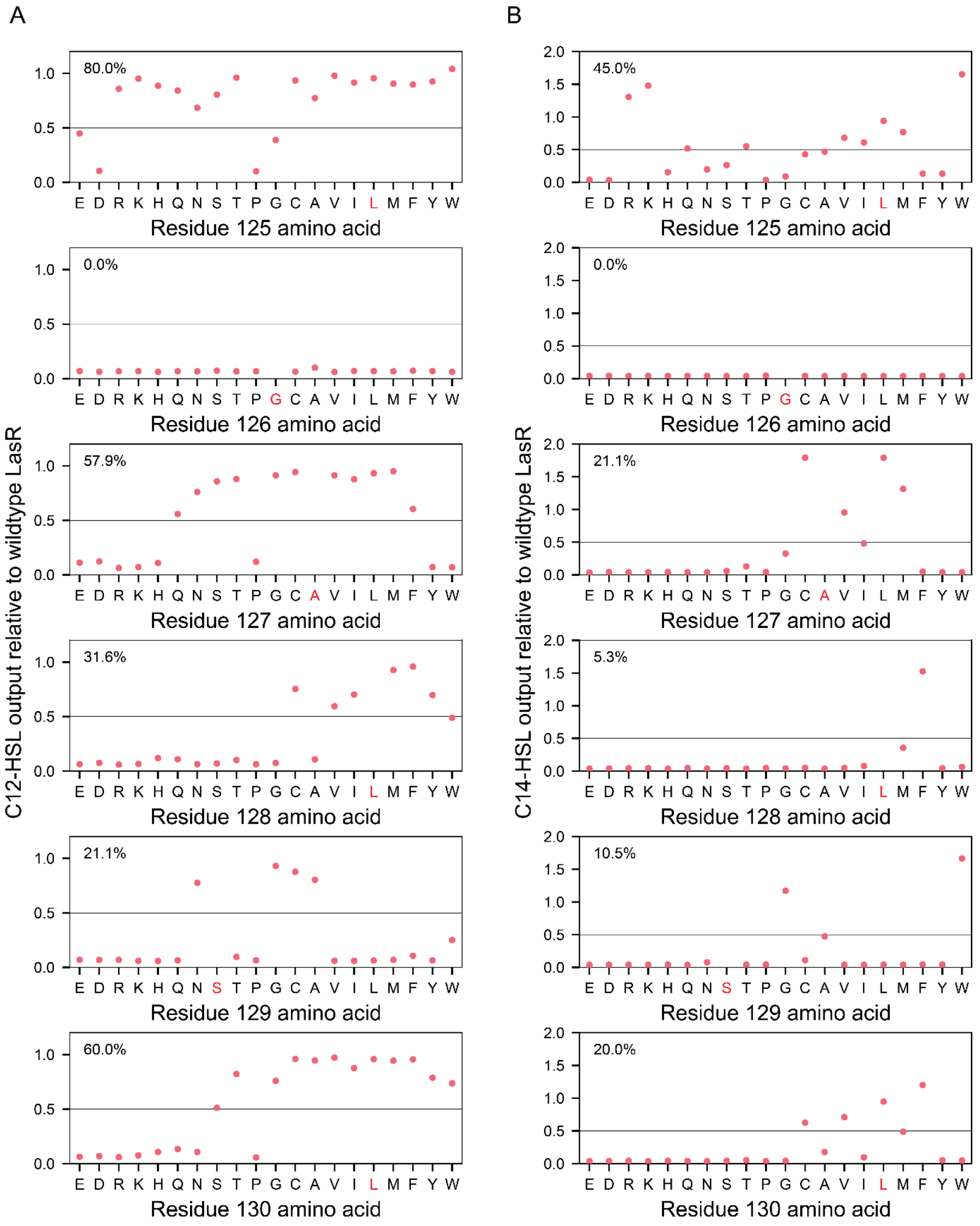


**Figure S10. Sensor output for single mutations relative to wildtype LasR.** The average sort-seq output for each LasR design was used to calculate its relative output (output for variant / output for wildtype LasR). **(A)** Activation by C12-HSL and **(B)** C14-HSL for each single mutation design is plotted. The percent of designs having at least 50% of the sensor output (black horizontal line, 9.85 RPU for C12-HSL and 1.22 RPU for C14-HSL) with wildtype LasR (amino acid in red font) is shown (top left).

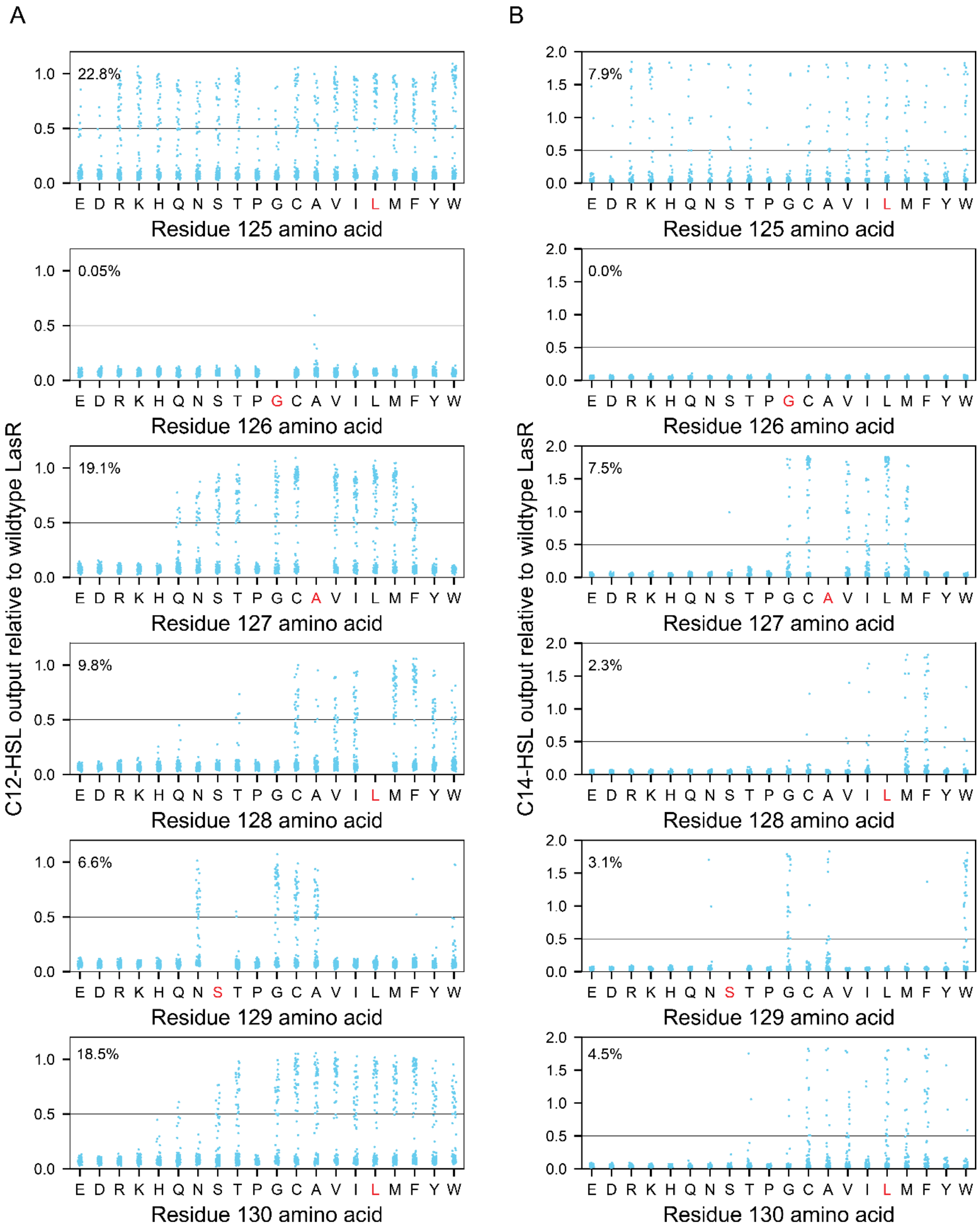


**Figure S11. Sensor output for double mutations relative to wildtype LasR.** The average sort-seq output for each LasR design was used to calculate its relative output (output for variant / output for wildtype LasR). **(A)** Activation by C12-HSL and **(B)** C14-HSL for each double mutation design is plotted. The percent of designs having at least 50% of the sensor output (black horizontal line, 9.85 RPU for C12-HSL and 1.22 RPU for C14-HSL) with wildtype LasR (amino acid in red font) is shown (top left).

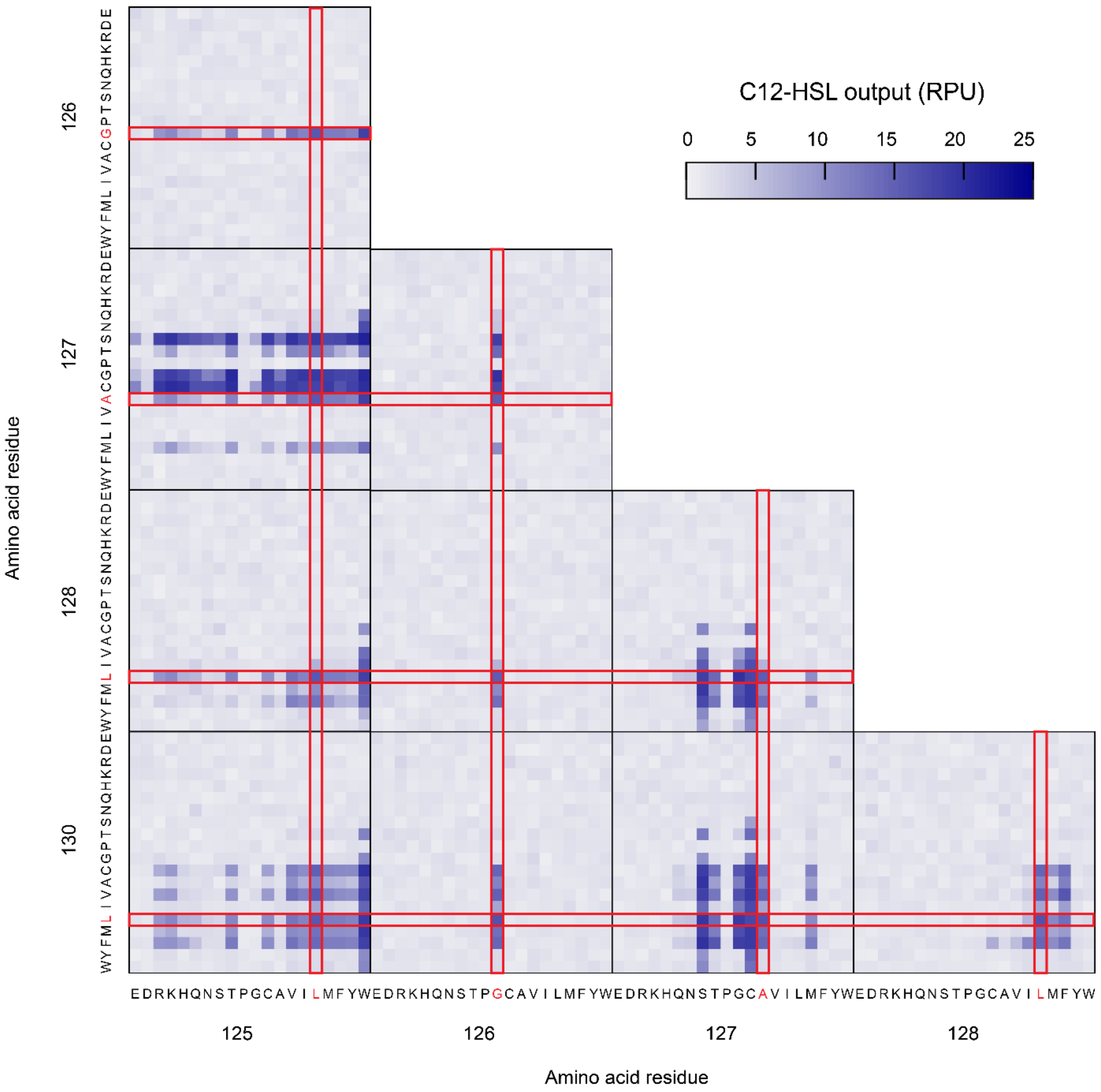


**Figure S12. Sequence-function relationship for activation by C12-HSL and LasR protein sequences with 3 amino acid substitutions.** An enlarged version of the left panel of Figure 4B is shown to indicate amino acids in each position using 1-letter abbreviations. Sensor output with C12-HSL was mapped against amino acids. The amino acid in wildtype LasR is shown (red). Output was determined by the average of two replicates of sort-seq using 1 μM C12-HSL (Methods).

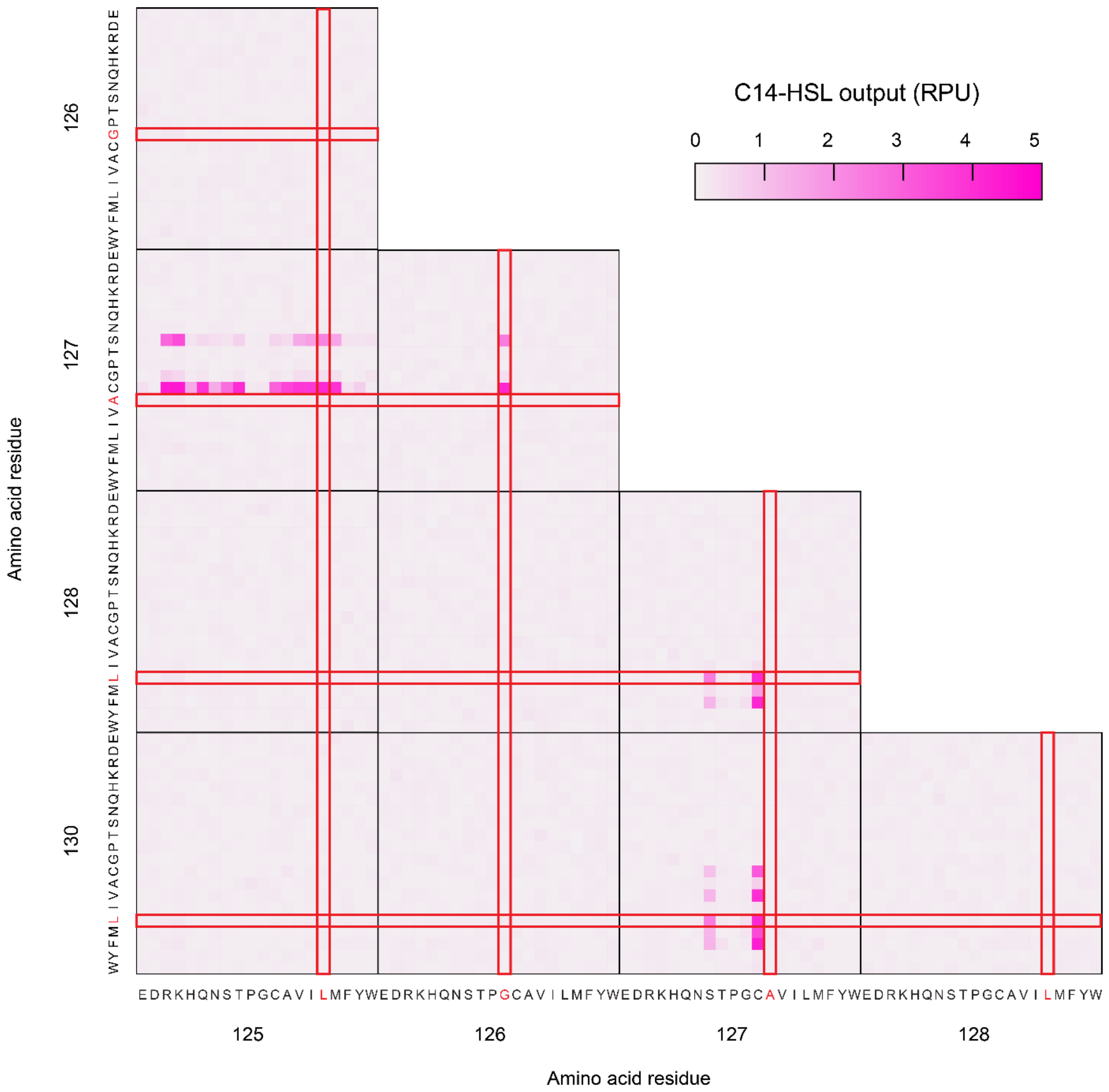


**Figure S13. Sequence-function relationship for activation by C14-HSL and LasR protein with 3 amino acid substitutions.** An enlarged version of the middle panel of Figure 4B is shown to indicate amino acids in each position using 1-letter abbreviations. Sensor output with C14-HSL was mapped against amino acids. The amino acid in wildtype LasR is shown (red). Output was determined by the average of two replicates of sort-seq using 2 μM C14-HSL (Methods).

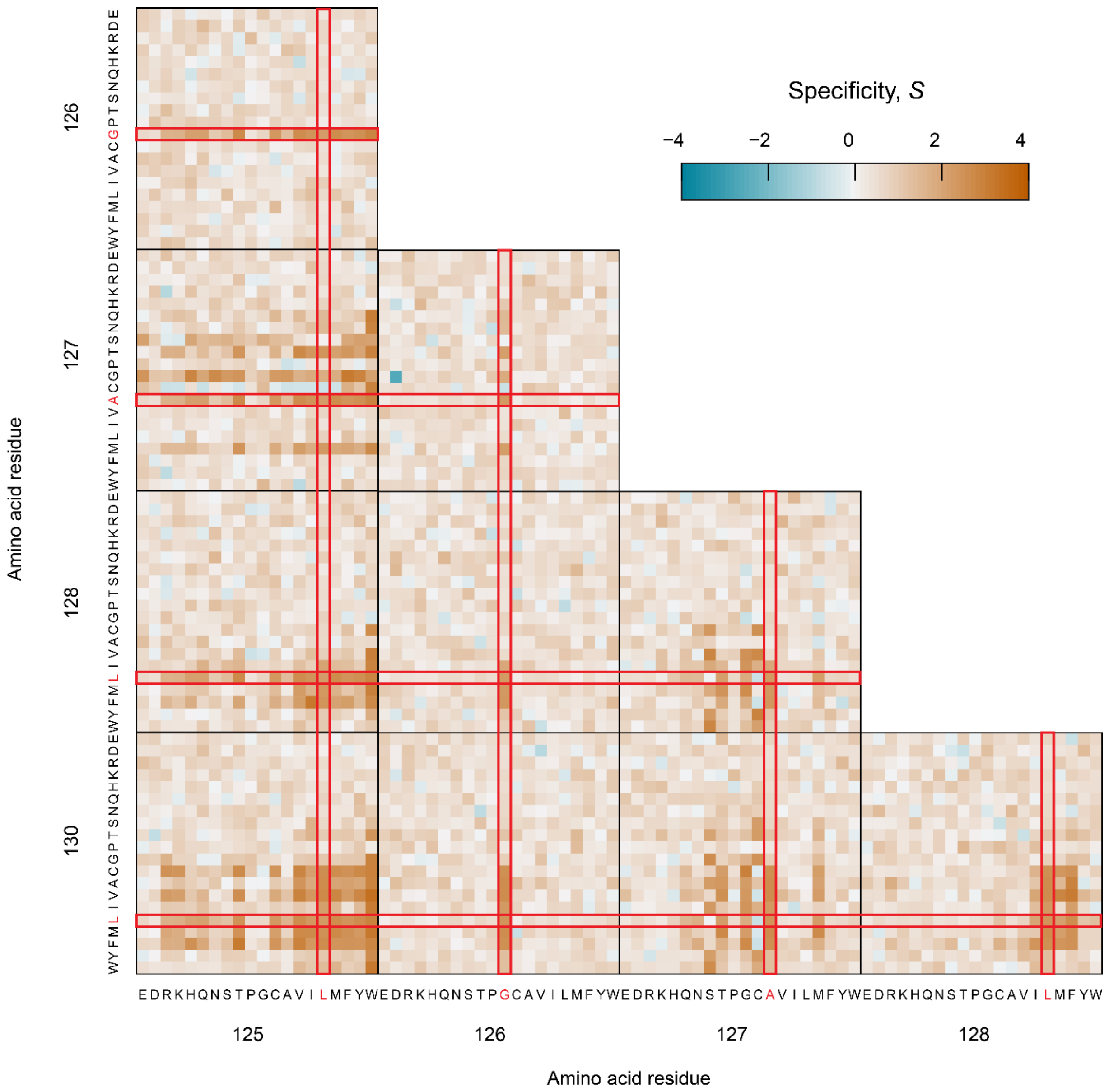


**Figure S14. Sequence-function relationship for specificity and LasR protein sequence with 3 amino acid substitutions.** An enlarged version of the right panel of Figure 4B is shown. Amino acid identity is listed using its 1-letter abbreviation. Specificity (*S*) was determined by the average of two replicates of sort-seq and is plotted for each amino acid in each position of the LasR variable region. The amino acid in wildtype LasR is shown (red).

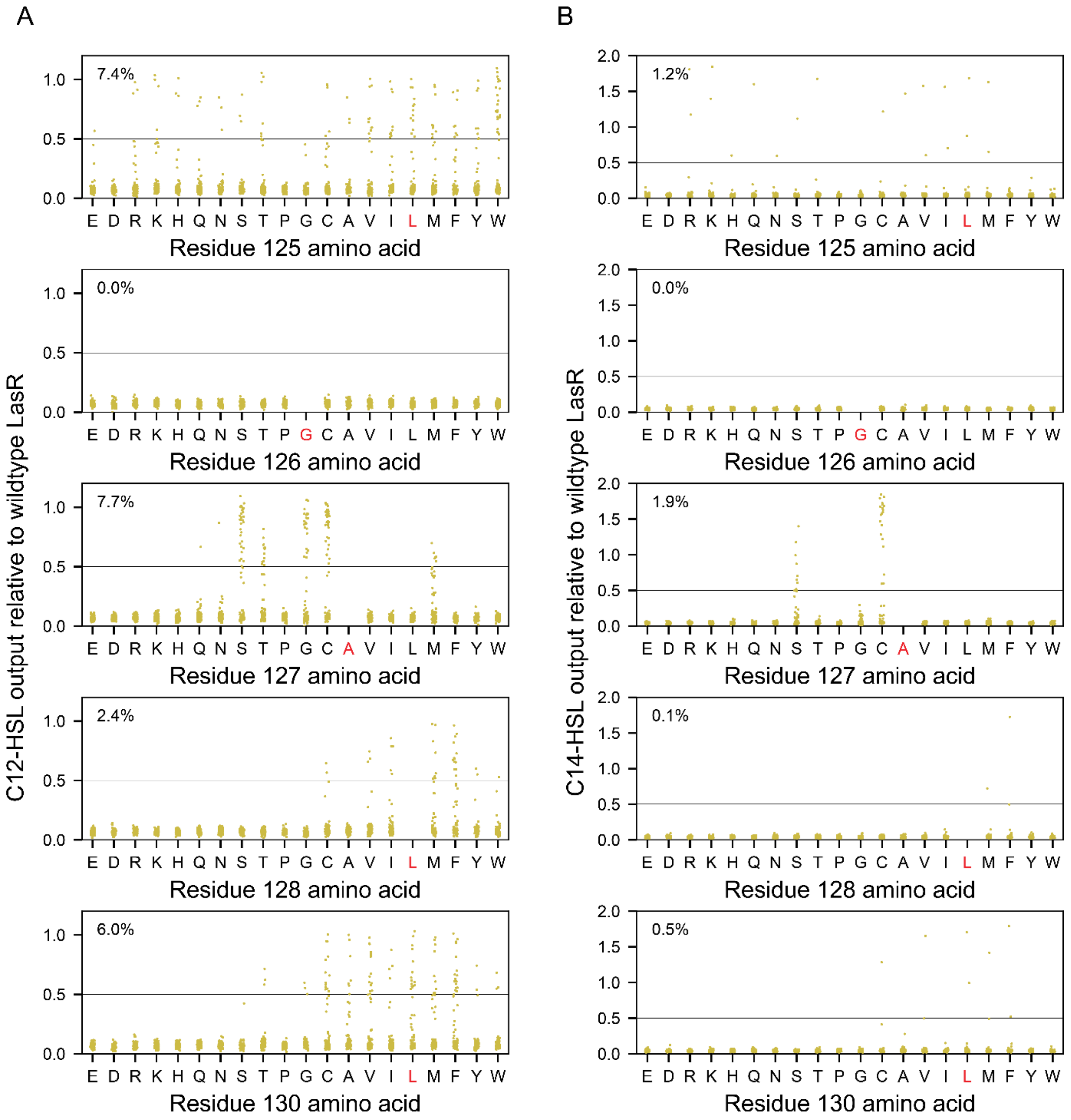


**Figure S15. Sensor output for triple mutations relative to wildtype LasR.** The average sort-seq output for each LasR design was used to calculate its relative output (output for variant / output for wildtype LasR). **(A)** Activation by C12-HSL and **(B)** C14-HSL for each triple mutation design is plotted. The percent of designs having at least 50% of the sensor output (black horizontal line, 9.85 RPU for C12-HSL and 1.22 RPU for C14-HSL) with wildtype LasR (amino acid in red font) is shown (top left).

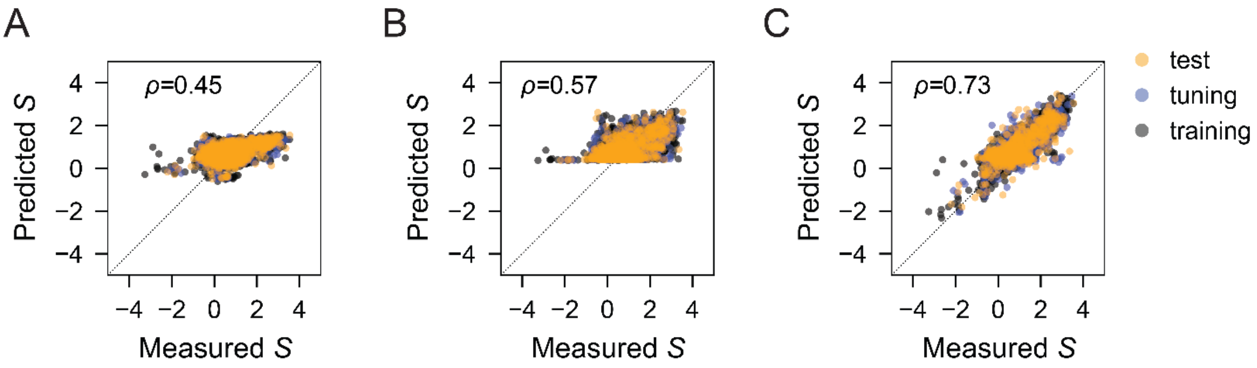


**Figure S16. Comparison of three models to predict LasR specificity from the protein sequence. (A)** Linear regression, **(B)** fully connected network, and **(C)** convolutional neural network models were constructed using a published script from other researchers (3). A randomly sampled testing set (orange, 20% of variants) remained constant for all testing sets to evaluate model predictions. To generate the models, a training set (gray, 60% of LasR variants) and tuning set (blue, 20% of LasR variants) were randomly sampled three times from the remaining sort-seq data excluding the testing set. One representative machine learning model for each is shown. Models were evaluated using the testing dataset, and the Pearson correlation coefficient ($\rho$) was determined for each model. The displayed convolutional neural network model that achieved the highest Pearson correlation coefficient ($\rho$) was utilized in designing LasR with highest specificity. Dotted line shows the 1:1 relationship.

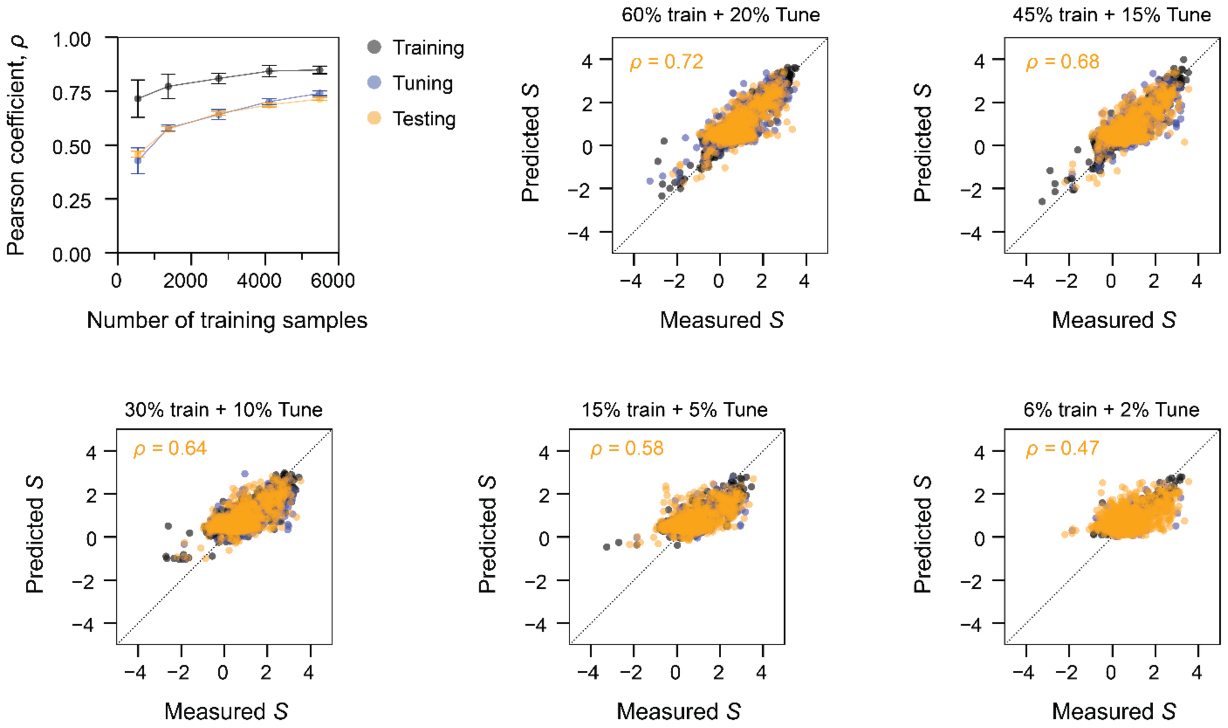


**Figure S17. Effect of varying the training set size for generating convolutional neural network models.** Convolutional neural network models were trained using different sizes of training sets and tuning sets. Pearson correlation coefficients ($\rho$) of the models were determined. The size of a tuning set (blue) was set to 1/3 of a corresponding training set size (gray). The randomly sampled testing set in Figure 5 and Figure S16, consisting of 1,828 variants (orange, 20% of variants), remained constant for all testing sets to evaluate model predictions. Training sets and tuning sets of five different sizes were randomly sampled three times from the remaining sort-seq data excluding the testing set. Pearson correlation coefficients were the average of three models trained from three training sets of the same size. Error bars are standard deviation. Representative machine learning models trained from 5 different numbers of samples are shown. Models were trained using: (i) 5,484-variant training set (60% of variants) and 1,828-variant tuning set (20% variants), (ii) 4,112-variant training set (45% of variants) and 1,371-variant tuning set (15% variants), (iii) 2,742-variant training set (30% of variants) and 914-variant tuning set (10% variants), (iv) 1,371-variant training set (15% of variants) and a 457-variant tuning set (5% variants), or (v) 548-variant training set (6% of variants) and 183-variant tuning set (2% variants). Dotted lines show the 1:1 relationship.

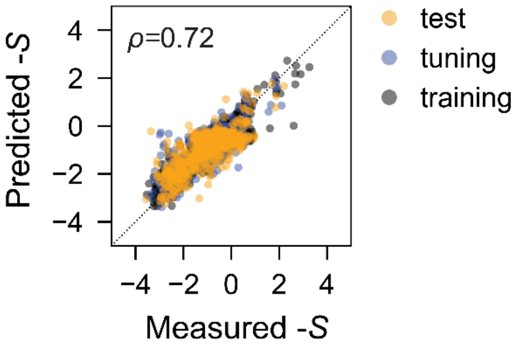


**Figure S18. Convolutional neural network model for reverse specificity.** To use the same algorithm to search for LasR variants with minimal specificity, convolutional neural network models for reverse specificity ($-1 \times S$) were constructed (3). The same testing set (orange, 20% of the variants) employed in Figure S16 was used to evaluate the model's predictions for reverse specificity. Three different neural network models were generated using three randomly sampled sets of the training set (colored in gray, representing 60% of LasR variants) and the tuning set (colored in blue, comprising 20% of LasR variants) utilized in Figure S16. A comparison of the predicted and measured reverse specificity is plotted using the model that achieved the highest Pearson correlation coefficient ($\rho$) for the testing set. This model was subsequently used for the design of LasR variants with preferential specificity to noncognate HSL. Dotted line shows the 1:1 relationship.

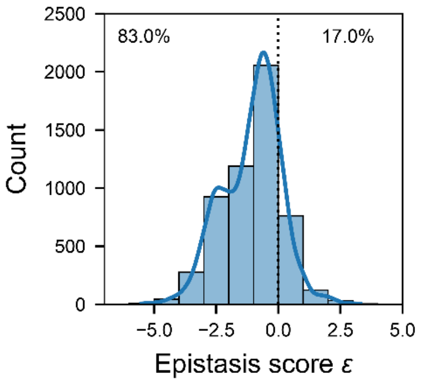


**Figure S19. Pairwise epistasis score distribution of LasR double mutation variants.** Pairwise epistasis scores of double mutation variants were calculated as described in Methods. A kernel density (blue line) was added to the histogram to visualize distribution in a continuous manner. The percentage of negative epistasis ($\varepsilon < 0$) and positive epistasis ($\varepsilon > 0$) are shown on the graph.

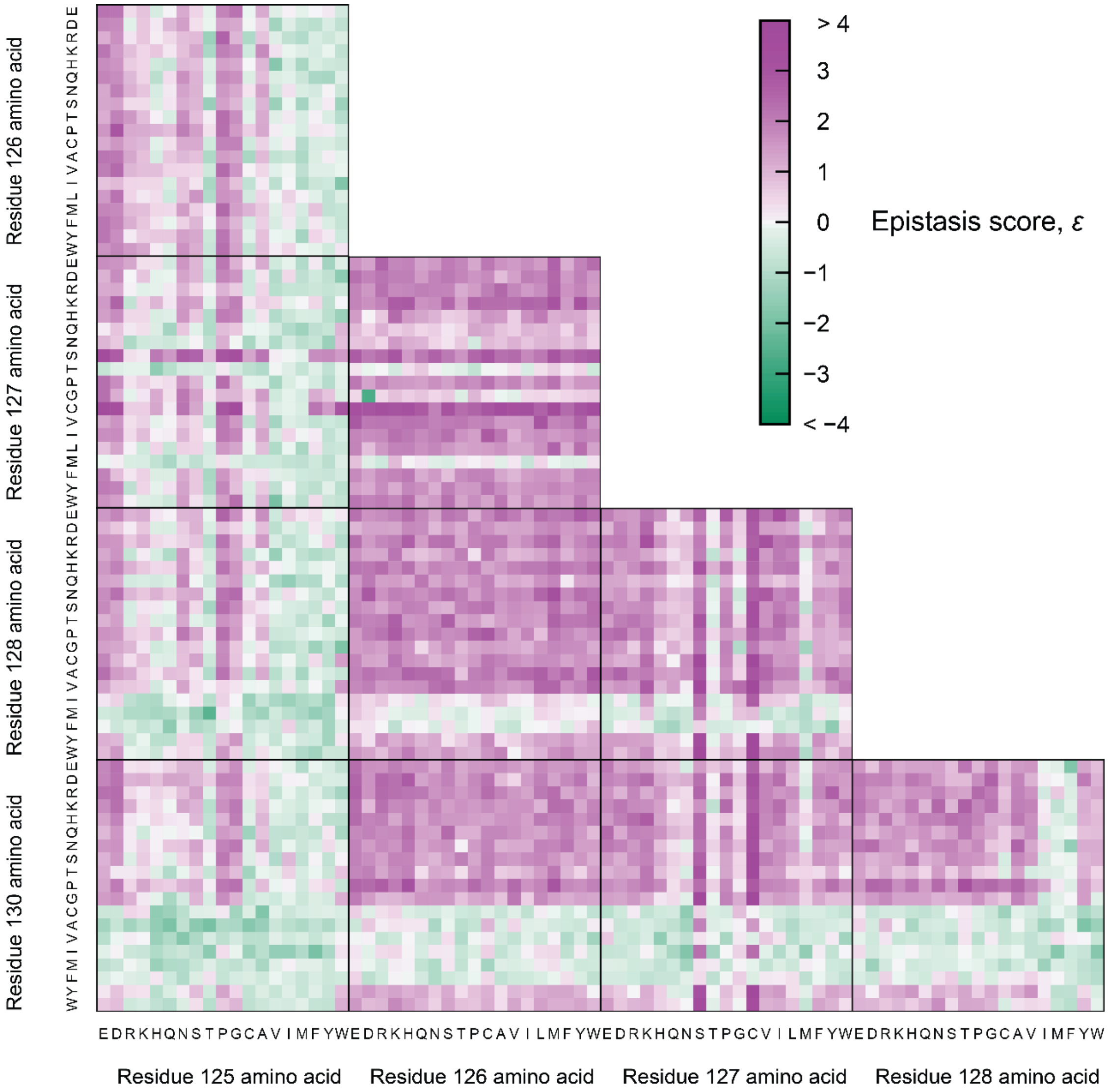


**Figure S20. Pairwise epistasis scores of LasR triple mutation variants in the S129N genetic background.** Heat maps represent pairwise epistasis scores of triple mutation variants under S129N genetic background ($\varepsilon$) mapped against the amino acid identity at each position. Pairwise epistasis under S129N genetic background was determined as defined in Methods. Position 129 is not plotted as it was fixed and present in all triple mutation designs.

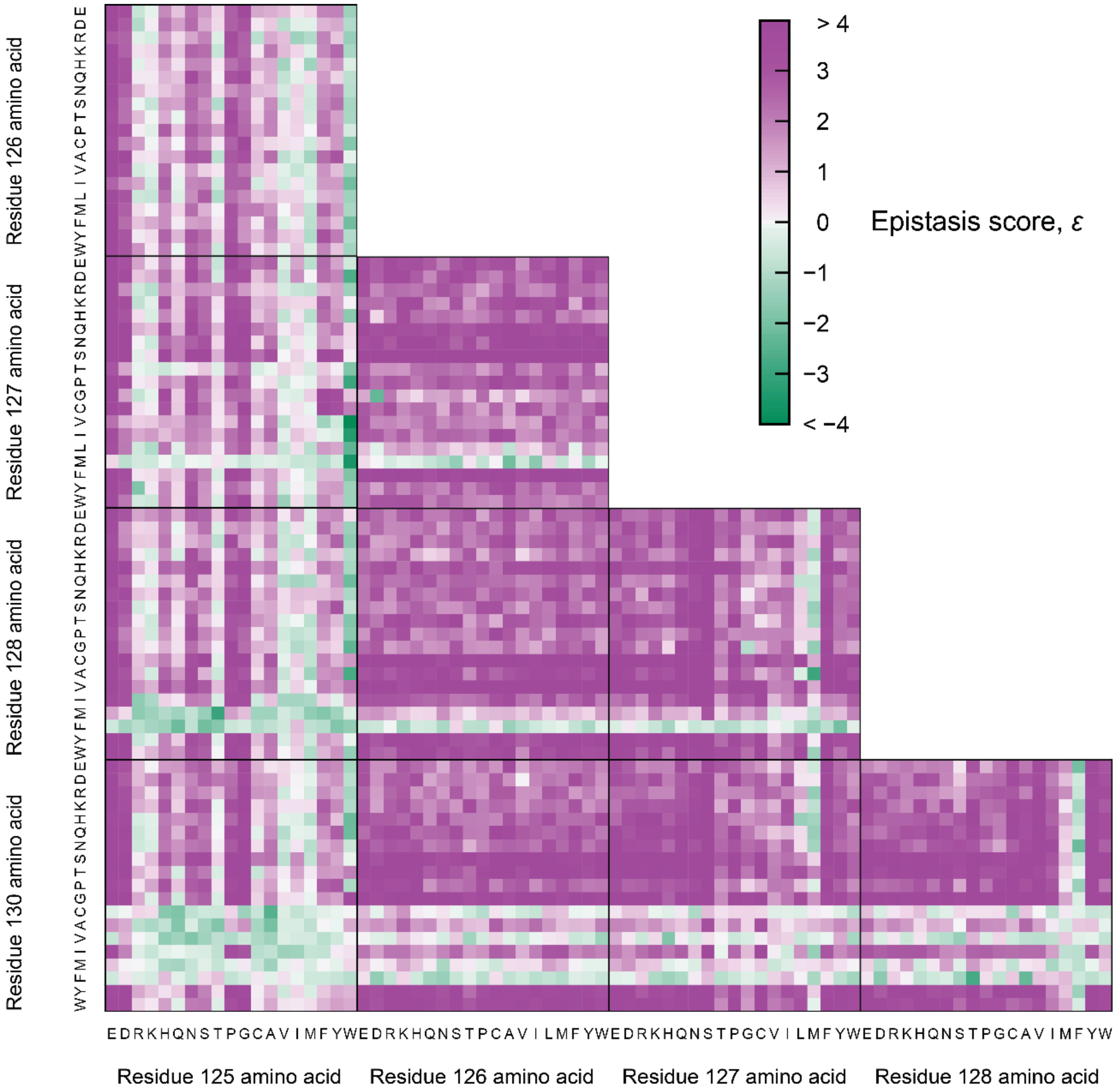


**Figure S21. Third order epistasis scores of LasR triple mutation variants.** Heat maps represent third order epistasis scores of triple mutation variants ($\varepsilon$) mapped against the amino acid identity at each position. Third order epistasis was calculated as defined in Methods. Position 129 is not plotted as it was fixed and present in all triple mutation designs.

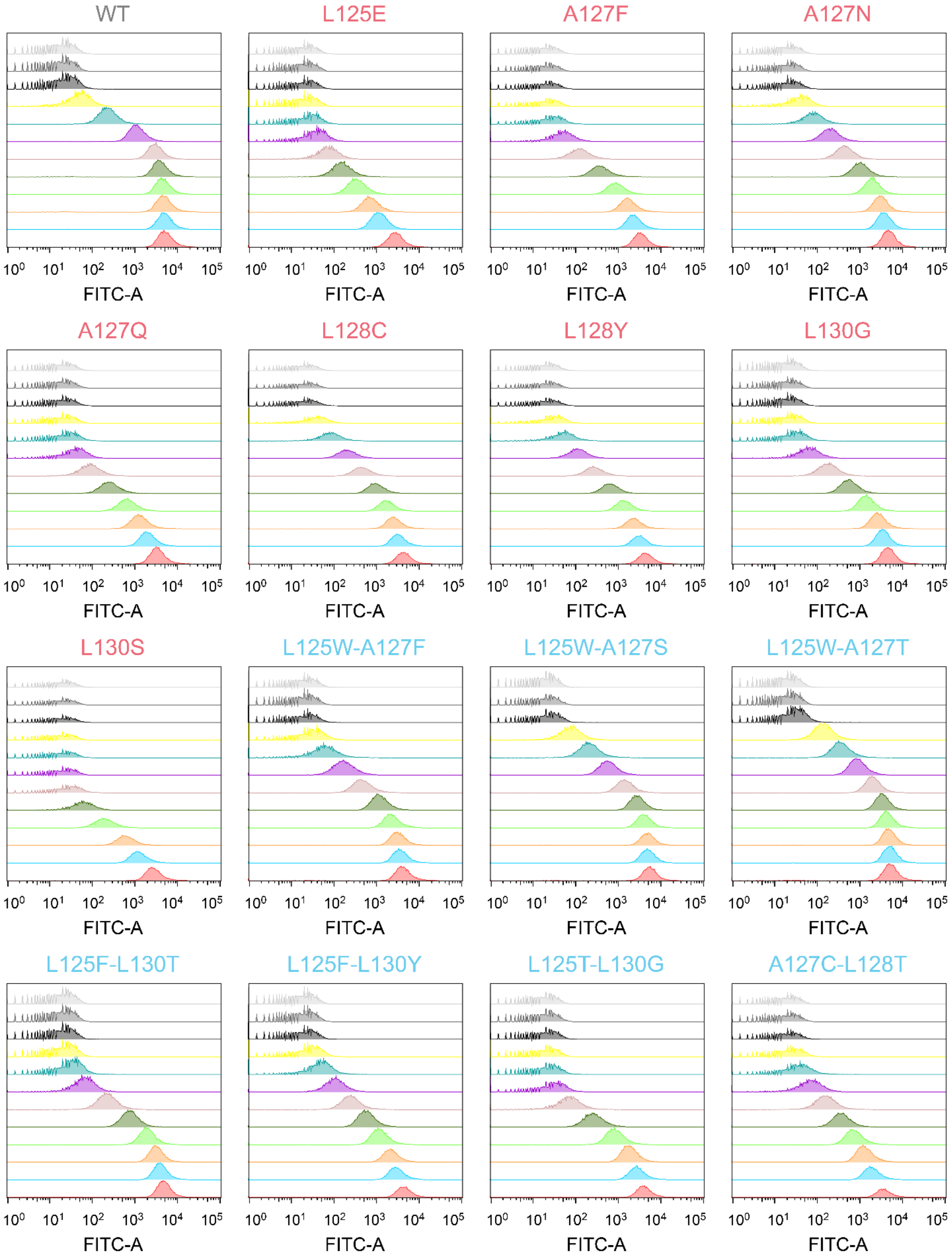
WT
L125E
A127F
A127N
A127Q
L128C
L128Y
L130G
L130S
L125W-A127F
L125W-A127S
L125W-A127T
L125F-L130T
L125F-L130Y
L125T-L130G
A127C-L128T
FITC-A
$10^0$ $10^1$ $10^2$ $10^3$ $10^4$ $10^5$

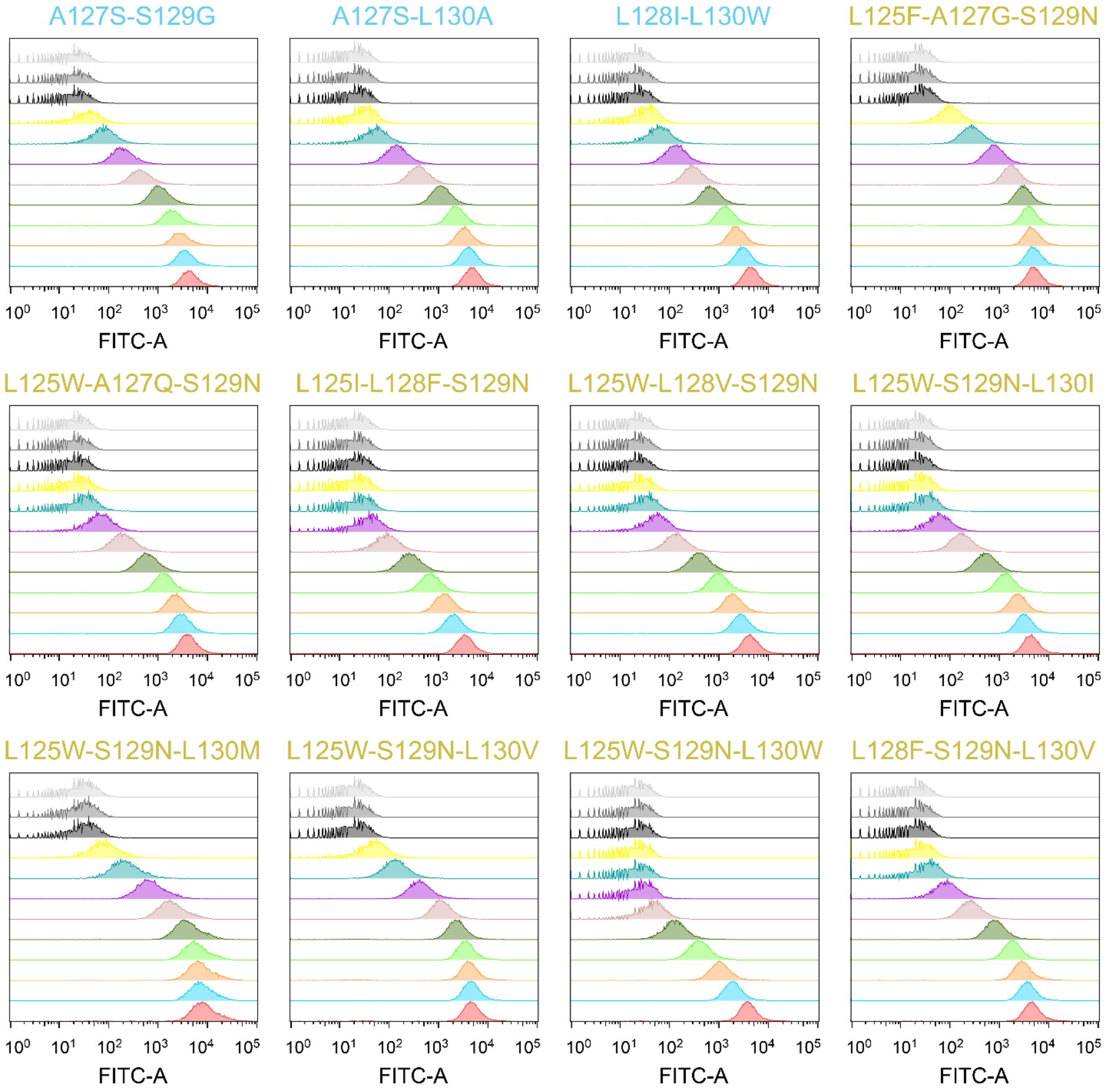


**Figure S22. Representative histograms of cell fluorescence from response function characterization.** One representative histogram from each response function characterization by flow cytometry for each variant and HSL concentration in Figure 7 is shown. The concentration of C12-HSL for WT and other variants are the same as Figure 7 and increases from top to bottom in each panel. The LasR sequence is indicated at the top of each panel. This is a 2-page figure.

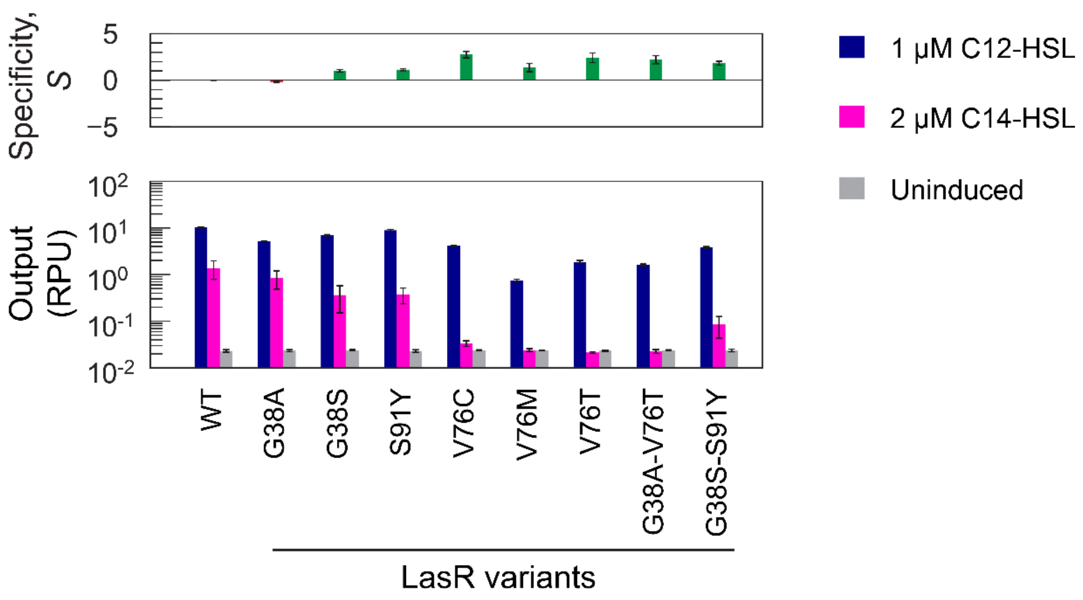


**Figure S23. Sensor characterization of LasR variants with mutations outside of LasR β5.** Site-directed mutagenesis in residues outside of the LasR β5 sheet was conducted. Selected residues were identified by covariation analysis in a previous study as being essential for specificity (2). LasR sensors were assayed without inducer (light grey), with 1 μM 3OC12-HSL (dark blue), or with 2 μM 3OHC14-HSL (magenta). The cell fluorescence was measured by flow cytometry, and the arbitrary unit was converted to standard RPU (Methods). The specificity of LasR sensor variants on each day was calculated as described in Methods. Bars indicate the mean ± s.d ($n$ = 3 biological replicates). LasR sensors containing V76C, V76T, V76M, G38A-V76T, or G38S-S91Y demonstrated significantly decreased C14-HSL output ($p$ = 0.031 – 0.036). However, those sensors also showed significantly decreased C12-HSL output ($p < 0.001$), ranging from 2.49-fold to 13.9-fold less than the wildtype (WT) LasR sensor.

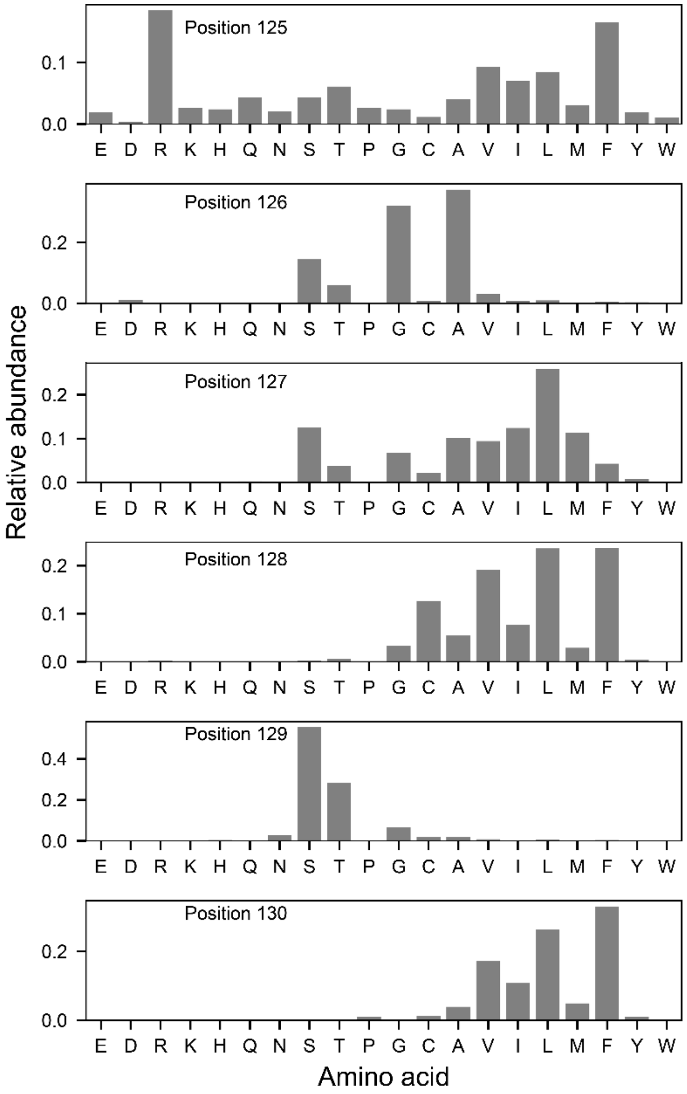


**Figure S24. Natural relative abundance of each amino acid at each position in LasR homologs.** We reduced a previously published aligned set of 6,360 natural LasR homologs (2) to 3,098 LasR homologs by removing sequences with similarity larger than 90%. Relative abundance was calculated as the frequency of an amino acid at the indicated position among the set of 3,098 LasR homologs.

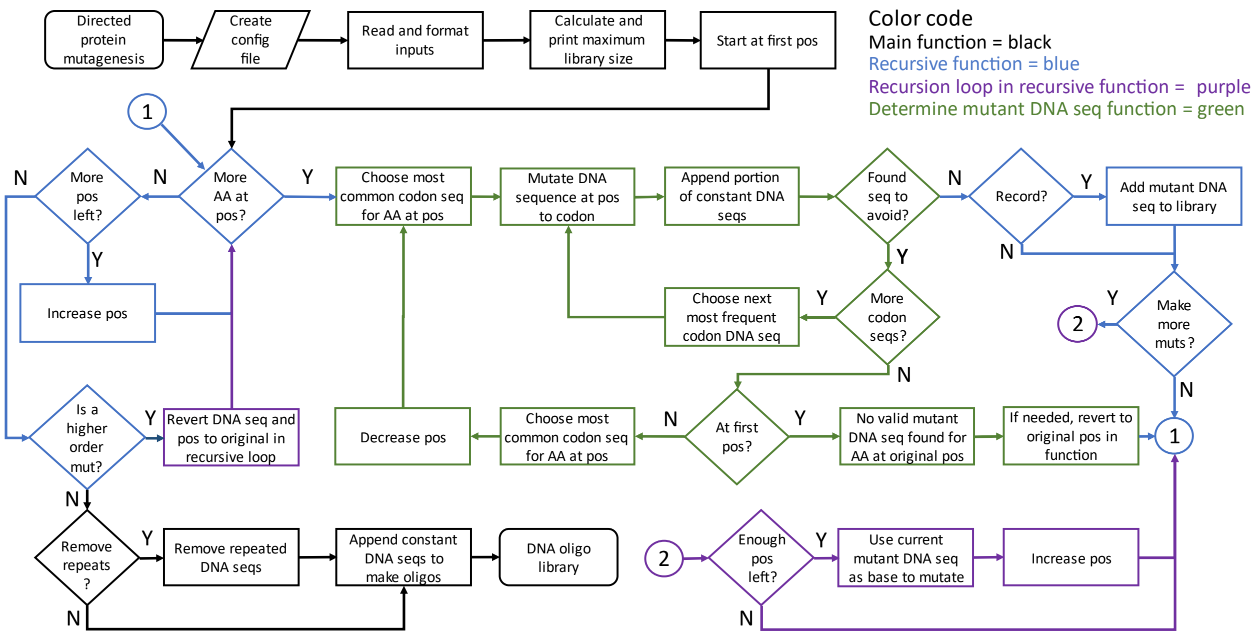


**Figure S25. Oligo design algorithm.** Flow chart of the custom Python script used to design the oligo pool for combinatorial saturation mutagenesis library is depicted. The start and end of the script (rounded rectangle), inputs (parallelogram), processes (right rectangle), decision points (diamonds), and connections to other parts of the algorithm (numbered circles) are indicated. The flow chart indicates main functions (black), recursive functions (blue), loops (purple), and DNA sequence generation (green). Config = configuration, pos = position in the protein sequence, mut = mutant, seq = DNA sequence.

**Table S1. Summary of LasR single mutation variants that reduced specificity but had positive epistasis with other mutations.**

| Variant | C12 output (RPU) | C14 output (RPU) | Specificity, $S$ |
|---|---|---|---|
| WT | 19.7 | 2.4 | 0 |
| L125R | 16.8 | 3.2 | -0.42 |
| L125K | 18.7 | 3.6 | -0.44 |
| L125W | 20.4 | 4.0 | -0.46 |
| A127C | 18.5 | 4.4 | -0.64 |
| A127V | 17.9 | 2.3 | -0.04 |
| A127L | 18.3 | 4.4 | -0.65 |
| A127M | 18.7 | 3.2 | -0.32 |
| L128F | 18.9 | 3.7 | -0.46 |
| S129G | 18.3 | 2.9 | -0.23 |
| S129W | 4.9 | 4.1 | -1.90 |
| L130F | 18.8 | 2.9 | -0.23 |

**Table S2. Epistasis scores for selected LasR double mutation variants having improved specificity**

| Variant | $S_a$ | $S_b$ | $S_{ab}$ | $\varepsilon_{ab}$ |
|---|---|---|---|---|
| L125W-A127F | -0.46 | 2.53 | 3.31 | 1.24 |
| L125W-A127S | -0.46 | 2.68 | 3.54 | 1.32 |
| L125W-A127T | -0.46 | 1.93 | 3.22 | 1.75 |
| L125F-L130T | 1.90 | 2.73 | 3.25 | -1.38 |
| L125F-L130Y | 1.90 | 2.73 | 3.53 | -1.1 |
| L125T-L130G | 0.56 | 2.91 | 3.43 | -0.04 |
| A127C-L128T | -0.64 | 0.96 | 3.33 | 3.01 |
| A127S-S129G | 2.68 | -0.23 | 3.35 | 0.9 |
| A127S-L130A | 2.68 | 1.68 | 3.23 | -1.13 |
| L128I-L130W | 2.22 | 2.80 | 3.22 | -1.8 |

**Table S3. Epistasis scores of selected LasR triple mutation variants with improved specificity ($S$).**

| Variant | $S_a$ | $S_b$ | $S_c$ | $S_{ab}$ | $S_{ac}$ | $S_{bc}$ | $S_{abc}$ | $\varepsilon_{abc}$ |
|---|---|---|---|---|---|---|---|---|
| L125F-A127G-S129N | 1.90 | 1.03 | 2.31 | -0.29 | 2.49 | 1.89 | 3.17 | 4.32 |
| L125W-A127Q-S129N | -0.46 | 2.61 | 2.31 | 2.76 | 2.90 | 1.51 | 3.07 | 0.36 |
| L125I-L128F-S129N | 0.40 | -0.46 | 2.31 | -0.38 | 2.52 | 2.42 | 3.25 | 0.94 |
| L125W-L128V-S129N | -0.46 | 2.52 | 2.31 | 1.47 | 2.90 | 0.60 | 3.04 | 2.44 |
| L125T-S129N-L130F | 0.56 | 2.31 | -0.23 | 2.74 | -0.19 | 2.60 | 3.18 | 0.67 |
| L125W-S129N-L130I | -0.46 | 2.31 | 2.20 | 2.90 | 1.72 | 2.36 | 3.15 | 0.22 |
| L125W-S129N-L130M | -0.46 | 2.31 | 0.66 | 2.90 | -0.25 | 2.67 | 3.23 | 0.42 |
| L125W-S129N-L130V | -0.46 | 2.31 | 0.31 | 2.90 | -0.17 | 2.67 | 3.46 | 0.22 |
| L125W-S129N-L130W | -0.46 | 2.31 | 2.80 | 2.90 | 2.26 | 1.35 | 3.21 | 1.35 |
| L128F-S129N-L130V | -0.46 | 2.31 | 0.31 | 2.42 | 0.12 | 2.67 | 3.14 | 0.09 |

**Table S4. Sensor response function parameters for the set of assayed LasR variants in Figure 7**

| LasR variant | $y_{min}$ | $y_{max}$ | $EC_{50}$ | $n$ |
|---|---|---|---|---|
| WT | 0.008 | 11.0 | 0.04 | 1.2 |
| L125E | 0.010 | 6.8 | 1.04 | 1.5 |
| A127F | 0.008 | 7.4 | 0.40 | 1.8 |
| A127N | 0.010 | 10.5 | 0.28 | 1.5 |
| A127Q | 0.008 | 7.3 | 0.55 | 1.7 |
| L128C | 0.009 | 9.0 | 0.25 | 1.5 |
| L128Y | 0.009 | 9.7 | 0.39 | 1.5 |
| L130G | 0.009 | 9.8 | 0.36 | 1.9 |
| L130S | 0.009 | 6.4 | 0.95 | 2.1 |
| L125W-A127F | 0.011 | 8.7 | 0.19 | 1.8 |
| L125W-A127S | 0.009 | 11.0 | 0.10 | 1.7 |
| L125W-A127T | 0.009 | 10.6 | 0.08 | 1.6 |
| L125F-L130T | 0.011 | 9.7 | 0.31 | 2.0 |
| L125F-L130Y | 0.009 | 9.4 | 0.43 | 1.5 |
| L125T-L130G | 0.010 | 8.4 | 0.43 | 2.2 |
| A127C-L128T | 0.009 | 7.1 | 0.52 | 1.5 |
| A127S-S129G | 0.009 | 9.2 | 0.23 | 1.6 |
| A127S-L130A | 0.010 | 9.7 | 0.25 | 1.8 |
| L128I-L130W | 0.009 | 9.1 | 0.40 | 1.5 |
| L125F-A127G-S129N | 0.010 | 9.6 | 0.08 | 1.7 |
| L125W-A127Q-S129N | 0.009 | 7.6 | 0.30 | 1.9 |
| L125I-L128F-S129N | 0.009 | 6.5 | 0.52 | 1.8 |
| L125W-L128V-S129N | 0.009 | 8.4 | 0.44 | 1.8 |
| L125W-S129N-L130I | 0.010 | 8.6 | 0.36 | 1.9 |
| L125W-S129N-L130M | 0.032 | 18.7 | 0.13 | 1.8 |
| L125W-S129N-L130V | 0.009 | 9.6 | 0.11 | 1.9 |
| L125W-S129N-L130W | 0.011 | 8.2 | 0.82 | 2.0 |
| L128F-S129N-L130V | 0.010 | 8.9 | 0.27 | 2.0 |

**Table S5. Plasmids used in this work**

| Plasmids Name | Description |
|---|---|
| pMZ103-rbs1 | This LasR sensor containing the wildtype LasR |
| pMZ103-rbs1-wtBsaI | This LasR sensor containing the wildtype LasR and BsaI sites on the plasmids was removed. |
| pMZ103-rbs1-backbone | The LasR sensor library backbone. A LacZ marker (yellow) for blue-white screening was placed at the LasR 125 – 130 insertion site. |
| pMZ103-rbs1-m1 | A LasR quorum sensor containing LasR A127N |
| pMZ103-rbs1-m2 | A LasR quorum sensor containing LasR L128C |
| pMZ103-rbs1-m3 | A LasR quorum sensor containing LasR L125W-A127S |
| pMZ103-rbs1-m4 | A LasR quorum sensor containing LasR L125W-A127T |
| pMZ103-rbs1-m5 | A LasR quorum sensor containing LasR S129N-L125F-A127G |
| pMZ103-rbs1-m6 | A LasR quorum sensor containing LasR S129N-L125W-L130V |
| pMZ103-rbs1-m7 | A LasR quorum sensor containing LasR L125W-A127T-L128M |
| pMZ103-rbs1-m8 | A LasR quorum sensor containing LasR L128F-S129W |
| pMZ103-rbs1-L130G | A LasR quorum sensor containing L130G |
| pMZ103-rbs1-L130W | A LasR quorum sensor containing LasR L130W |
| pMZ103-rbs1-L128Y | A LasR quorum sensor containing LasR L128Y |
| pMZ103-rbs1-L130Y | A LasR quorum sensor containing LasR L130Y |
| pMZ103-rbs1-L130T | A LasR quorum sensor containing LasR L130T |
| pMZ103-rbs1-A127S | A LasR quorum sensor containing LasR A127S |
| pMZ103-rbs1-A127Q | A LasR quorum sensor containing LasR A127Q |
| pMZ103-rbs1-A127F | A LasR quorum sensor containing LasR A127F |
| pMZ103-rbs1-L128V | A LasR quorum sensor containing LasR L128V |
| pMZ103-rbs1-L130S | A LasR quorum sensor containing LasR L130S |
| pMZ103-rbs1-L125E | A LasR quorum sensor containing LasR L125E |
| pMZ103-rbs1-L128I | A LasR quorum sensor containing LasR L128I |
| pMZ103-rbs1-L128W | A LasR quorum sensor containing LasR L128W |
| pMZ103-rbs1-L125W | A LasR quorum sensor containing LasR L125W |
| pMZ103-rbs1-L125F-L130Y | A LasR quorum sensor containing LasR L125F-L130Y |
| pMZ103-rbs1-L125T-L130G | A LasR quorum sensor containing LasR L125T-L130G |
| pMZ103-rbs1-A127S-S129G | A LasR quorum sensor containing LasR A127S-S129G |
| pMZ103-rbs1-A127C-L128T | A LasR quorum sensor containing LasR A127C-L128T |
| pMZ103-rbs1-L125W-A127F | A LasR quorum sensor containing LasR L125W-A127F |
| pMZ103-rbs1-L125F-L130T | A LasR quorum sensor containing LasR L125F-L130T |
| pMZ103-rbs1-A127S-L130A | A LasR quorum sensor containing LasR A127S-L130A |
| pMZ103-rbs1-L128I-L130W | A LasR quorum sensor containing LasR L128I-L130W |
| pMZ103-rbs1-S129N-L125I-L128F | A LasR quorum sensor containing LasR S129N-L125I-L128F |
| pMZ103-rbs1-S129N-L125W-L130M | A LasR quorum sensor containing LasR S129N-L125W-L130M |
| pMZ103-rbs1-S129N-L125W-L130W | A LasR quorum sensor containing LasR S129N-L125W-L130W |
| pMZ103-rbs1-S129N-L125T-L130F | A LasR quorum sensor containing LasR S129N-L125T-L130F |

| | |
|---|---|
| pMZ103-rbs1-S129N-L125W-L130I | A LasR quorum sensor containing LasR S129N-L125W-L130I |
| pMZ103-rbs1-S129N-L128F-L130V | A LasR quorum sensor containing LasR S129N-L128F-L130V |
| pMZ103-rbs1-S129N-L125W-A127Q | A LasR quorum sensor containing LasR S129N-L125W-A127Q |
| pMZ103-rbs1-S129N-L125W-L128V | A LasR quorum sensor containing LasR S129N-L125W-L128V |
| pMZ103-rbs1-L125I-S129W | A LasR quorum sensor containing LasR L125I-S129W |
| pMZ103-rbs1-L125R-S129W | A LasR quorum sensor containing LasR L125R-S129W |
| pMZ103-rbs1-S129W-L130M | A LasR quorum sensor containing LasR S129W-L130M |
| pMZ103-rbs1-A127G-S129A | A LasR quorum sensor containing LasR A127G-S129A |
| pMZ103-rbs1-L125W-A127S-L128C-L130W | A LasR quorum sensor containing LasR L125W-A127S-L128C-L130W |
| pMZ103-rbs1-L125W-A127T-L128C-S129G-L130V | A LasR quorum sensor containing LasR L125W-A127T-L128C-S129G-L130V |
| pMZ103-rbs1-L125R-L128F-S129W | A LasR quorum sensor containing LasR L125R-L128F-S129W |
| pMZ103-rbs1-L125R-L128F-S129W-L130M | A LasR quorum sensor containing LasR L125R-L128F-S129W-L130M |
| pMZ103-rbs1-L130I | A LasR quorum sensor containing LasR L130I |
| pMZ103-rbs1-S129N-L130I | A LasR quorum sensor containing LasR S129N-L130I |
| pMZ103-rbs1-G38A | A LasR quorum sensor containing LasR G38A |
| pMZ103-rbs1-G38S | A LasR quorum sensor containing LasR G38S |
| pMZ103-rbs1-S91Y | A LasR quorum sensor containing LasR S91Y |
| pMZ103-rbs1-V76C | A LasR quorum sensor containing LasR V76C |
| pMZ103-rbs1-V76M | A LasR quorum sensor containing LasR V76M |
| pMZ103-rbs1-V76T | A LasR quorum sensor containing LasR V76T |
| pMZ103-rbs1-G38A-V76T | A LasR quorum sensor containing LasR G38A-V76T |
| pMZ103-rbs1-G38S-S91Y | A LasR quorum sensor containing LasR G38S-S91Y |

**Table S6. Genetic part sequence used in this work**

| Part Name | Type | DNA sequence | Reference |
|---|---|---|---|
| $P_{Las}$ | Promoter | TTCGAGCCTAGCAAGGGTCCGGGTTCACCGAAATCTA TCTCATTTGCTAGTTATAAAATTATGAAATTTGCGTAAA TTCTTCA | (4) |
| RiboJ | Insulator | AGCTGTCACCGGATGTGCTTTCCGGTCTGATGAGTCC GTGAGGACGAAACAGCCTCTACAAATAATTTTGTTTAA | (5) |
| BBa_B0034 | Synthetic RBS | AAAGAGGAGAAA | (6) |
| *rbs1* | Synthetic RBS | TCTAATTTAACAAAAGGTTTTCCAATTT | This study |
| *yfp* | Gene | ATGGTGAGCAAGGGCGAGGAGCTGTTCACCGGGGTG GTGCCCATCCTGGTCGAGCTGGACGGCGACGTAAAC GGCCACAAGTTCAGCGTGTCCGGCGAGGGCGAGGGC GATGCCACCTACGGCAAGCTGACCCTGAAGTTCATCT GCACCACAGGCAAGCTGCCCGTGCCCTGGCCCACCC TCGTGACCACCTTCGGCTACGGCCTGCAATGCTTCGC CCGCTACCCCGACCACATGAAGCTGCACGACTTCTTC AAGTCCGCCATGCCCGAAGGCTACGTCCAGGAGCGC ACCATCTTCTTCAAGGACGACGGCAACTACAAGACCC GCGCCGAGGTGAAGTTCGAGGGCGACACCCTGGTGA ACCGCATCGAGCTGAAGGGCATCGACTTCAAGGAGGA CGGCAACATCCTGGGGCACAAGCTGGAGTACAACTAC AACAGCCACAACGTCTATATCATGGCCGACAAGCAGA AGAACGGCATCAAGGTGAACTTCAAGATCCGCCACAA CATCGAGGACGGCAGCGTGCAGCTCGCCGACCACTA CCAGCAGAACACCCCAATCGGCGACGGCCCCGTGCT GCTGCCCGACAACCACTACCTTAGCTACCAGTCCGCC CTGAGCAAAGACCCCAACGAGAAGCGCGATCACATGG TCCTGCTGGAGTTCGTGACCGCCGCCGGGATCACTCT CGGCATGGACGAGCTGTACAAGTAA | (7) |
| *lasR** | gene | ATGGCCTTGGTTGACGGTTTTCTTGAGCTGGAACGCT CAAGTGGAAAATTGGAGTGGAGCGCCATCCTGCAGAA GATGGCGAGCGACCTTGGATTCTCGAAGATCCTGTTC GGCCTGTTGCCTAAGGACAGCCAGGACTACGAGAACG CCTTCATCGTCGGCAACTACCCGGCCGCCTGGCGCGA GCATTACGACCGGGCTGGCTACGCGCGGGTCGACCC GACGGTCAGTCACTGTACCCAGAGCGTACTGCCGATT TTCTGGGAACCGTCCATCTACCAGACGCGAAAGCAGC ACGAGTTCTTCGAGGAAGCCTCGGCCGCCGGCCTGG TGTATGGGCTGACCATGCCGCTGCATGGTGCTCGCGG CGAACTCGGCGCGCTGAGCCTCAGCGTGGAAGCGGA AAACCGGGCCGAGGCCAACCGTTTCATGGAGTCGGTC CTGCCGACCCTGTGGATGCTCAAGGACTACGCACTGC AGAGCGGTGCCGGACTGGCCTTCGAACATCCGGTCA GCAAACCGGTGGTTCTGACCAGCCGGGAGAAGGAAG TGTTGCAGTGGTGCGCCATCGGCAAGACCAGTTGGGA GATATCGGTTATCTGCAACTGCTCGGAAGCCAATGTGA ACTTCCATATGGGAAATATTCGGCGGAAGTTCGGTGT GACCTCCCGCCGCGTAGCGGCCATTATGGCCGTTAAT TTGGGTCTTATTACTCTCTGA | (4) |
| *lacI* | Gene | ATGAAACCAGTAACGTTATACGATGTCGCAGAGTATGC CGGTGTCTCTTATCAGACCGTTTCCCGCGTGGTGAAC CAGGCCAGCCACGTTTCTGCGAAAACGCGGGAAAAAG TGGAAGCGGCGATGGCGGAGCTGAATTACATTCCCAA | (8) |

| | | | |
|---|---|---|---|
| | | CCGCGTGGCACAACAACTGGCGGGCAAACAGTCGTTG<br>CTGATTGGCGTTGCCACCTCCAGTCTGGCCCTGCACG<br>CGCCGTCGCAAATTGTCGCGGCGATTAAATCTCGCGC<br>CGATCAACTGGGTGCCAGCGTGGTGGTGTCGATGGTA<br>GAACGAAGCGGCGTCGAAGCCTGTAAAGCGGCGGTG<br>CACAATCTTCTCGCGCAACGCGTCAGTGGGCTGATCA<br>TTAACTATCCGCTGGATGACCAGGATGCCATTGCTGTG<br>GAAGCTGCCTGCACTAATGTTCCGGCGTTATTTCTTGA<br>TGTCTCTGACCAGACACCCATCAACAGTATTATTTTCT<br>CCCATGAGGACGGTACGCGACTGGGCGTGGAGCATC<br>TGGTCGCATTGGGTCACCAGCAAATCGCGCTGTTAGC<br>GGGCCCATTAAGTTCTGTCTCGGCGCGTCTGCGTCTG<br>GCTGGCTGGCATAAATATCTCACTCGCAATCAAATTCA<br>GCCGATAGCGGAACGGGAAGGCGACTGGAGTGCCAT<br>GTCCGGTTTTCAACAAACCATGCAAATGCTGAATGAGG<br>GCATCGTTCCCACTGCGATGCTGGTTGCCAACGATCA<br>GATGGCGCTGGGCGCAATGCGCGCCATTACCGAGTC<br>CGGGCTGCGCGTTGGTGCGGATATCTCGGTAGTGGG<br>ATACGACGATACCGAAGATAGCTCATGTTATATCCCGC<br>CGTTAACCACCATCAAACAGGATTTTCGCCTGCTGGG<br>GCAAACCAGCGTGGACCGCTTGCTGCAACTCTCTCAG<br>GGCCAGGCGGTGAAGGGCAATCAGCTGTTGCCAGTCT<br>CACTGGTGAAAAGAAAAACCACCCTGGCGCCCAATAC<br>GCAAACCGCCTCTCCCCGCGCGTTGGCCGATTCATTA<br>ATGCAGCTGGCACGACAGGTTTCCCGACTGGAAAGCG<br>GGCAGTGA | |
| *tetR* | Gene | ATGTCCAGATTAGATAAAAGTAAAGTGATTAACAGCGC<br>ATTAGAGCTGCTTAATGAGGTCGGAATCGAAGGTTTAA<br>CAACCCGTAAACTCGCCCAGAAGCTAGGTGTAGAGCA<br>GCCTACATTGTATTGGCATGTAAAAAATAAGCGGGCTT<br>TGCTCGACGCCTTAGCCATTGAGATGTTAGATAGGCA<br>CCATACTCACTTTTGCCCTTTAGAAGGGGAAAGCTGGC<br>AAGATTTTTTACGTAATAACGCTAAAAGTTTTAGATGTG<br>CTTTACTAAGTCATCGCGATGGAGCAAAAGTACATTTA<br>GGTACACGGCCTACAGAAAAACAGTATGAAACTCTCG<br>AAAATCAATTAGCCTTTTTATGCCAACAAGGTTTTTCAC<br>TAGAGAATGCATTATATGCACTCAGCGCTGTGGGGCA<br>TTTTACTTTAGGTTGCGTATTGGAAGATCAAGAGCATC<br>AAGTCGCTAAAGAAGAAAGGGAAACACCTACTACTGAT<br>AGTATGCCGCCATTATTACGACAAGCTATCGAATTATT<br>TGATCACCAAGGTGCAGAGCCAGCCTTCTTATTCGGC<br>CTTGAATTGATCATATGCGGATTAGAAAAACAACTTAAA<br>TGTGAAAGTGGGTCCTAA | (8) |
| *araC* | Gene | ATGGCTGAAGCGCAAAATGATCCCCTGCTGCCGGGAT<br>ACTCGTTTAATGCCCATCTGGTGGCGGGTTTAACGCC<br>GATTGAGGCCAACGGTTATCTCGATTTTTTTATCGACC<br>GACCGCTGGGAATGAAAGGTTATATTCTCAATCTCACC<br>ATTCGCGGTCAGGGGGTGGTGAAAAATCAGGGACGA<br>GAATTTGTTTGCCGACCGGGTGATATTTTGCTGTTCCC<br>GCCAGGAGAGATTCATCACTACGGTCGTCATCCGGAG<br>GCTCGCGAATGGTATCACCAGTGGGTTTACTTTCGTCC<br>GCGCGCCTACTGGCATGAATGGCTTAACTGGCCGTCA<br>ATATTTGCCAATACGGGGTTCTTTCGCCCGGATGAAGC<br>GCACCAGCCGCATTTCAGCGACCTGTTTGGGCAAATC<br>ATTAACGCCGGGCAAGGGGAAGGGCGCTATTCGGAG<br>CTGCTGGCGATAAATCTGCTTGAGCAATTGTTACTGCG | (9) |

| | | | |
|---|---|---|---|
| | | GCGCATGGAAGCGATTAACGAGTCGCTCCATCCACCG<br>ATGGATAATCGGGTACGCGAGGCTTGTCAGTACATCA<br>GCGATCACCTGGCAGACAGCAATTTTGATATCGCCAG<br>CGTCGCACAGCATGTTTGCTTGTCGCCGTCGCGTCTG<br>TCACATCTTTTCCGCCAGCAGTTAGGGATTAGCGTCTT<br>AAGCTGGCGCGAGGACCAACGTATCAGCCAGGCGAA<br>GCTGCTTTTGAGCACCACCCGGATGCCTATCGCCACC<br>GTCGGTCGCAATGTTGGTTTTGACGATCAACTCTATTT<br>CTCGCGGGTATTTAAAAAATGCACCGGGGCCAGCCCG<br>AGCGAGTTCCGTGCCGGTTAA | |
| L3S2P21 | Terminator | CTCGGTACCAAATTCCAGAAAAGAGGCCTCCCGAAAG<br>GGGGGCCTTTTTTCGTTTTGGTCC | (10) |

* 9,485 LasR variants from the combinatorial mutagenesis library and 8 LasR variants outside of β5 sheet are not listed.

**Table S7. Primers designed for amplicon amplification in Next-gen sequencing.**

| Name | Sequence |
|---|---|
| LasR_NGS_FW1 | GAAGCTGGTGTATGGGCTGACCA |
| LasR_NGS_FW2 | GTAGCTGGTGTATGGGCTGACCA |
| LasR_NGS_FW3 | CTAGCTGGTGTATGGGCTGACCA |
| LasR_NGS_FW4 | AGTGCTGGTGTATGGGCTGACCA |
| LasR_NGS_FW5 | ACTGCTGGTGTATGGGCTGACCA |
| LasR_NGS_FW6 | TGTGCTGGTGTATGGGCTGACCA |
| LasR_NGS_FW7 | TGAGCTGGTGTATGGGCTGACCA |
| LasR_NGS_FW8 | TTGGCTGGTGTATGGGCTGACCA |
| LasR_NGS_FW9 | CATGCTGGTGTATGGGCTGACCA |
| LasR_NGS_FW10 | AACGGCTGGTGTATGGGCTGACCA |
| LasR_NGS_FW11 | ATCGGCTGGTGTATGGGCTGACCA |
| LasR_NGS_FW12 | ACGAGCTGGTGTATGGGCTGACCA |
| LasR_NGS_FW13 | TACGGCTGGTGTATGGGCTGACCA |
| LasR_NGS_FW14 | GTAAGCTGGTGTATGGGCTGACCA |
| LasR_NGS_FW15 | TCACGCTGGTGTATGGGCTGACCA |
| LasR_NGS_FW16 | ATCTGCTGGTGTATGGGCTGACCA |
| LasR_NGS_FW17 | TAGAGCTGGTGTATGGGCTGACCA |
| LasR_NGS_FW18 | GAAGGCTGGTGTATGGGCTGACCA |
| LasR_NGS_RV1 | GCAAAGTGAAACGGTTGGCCTCGG |
| LasR_NGS_RV2 | ATACACTGAAACGGTTGGCCTCGG |
| LasR_NGS_RV3 | AACGAATGAAACGGTTGGCCTCGG |
| LasR_NGS_RV4 | TACCTGTGAAACGGTTGGCCTCGG |
| LasR_NGS_RV5 | AAGGTATGAAACGGTTGGCCTCGG |
| LasR_NGS_RV6 | CCTCTGTGAAACGGTTGGCCTCGG |
| LasR_NGS_RV7 | GCTCTATGAAACGGTTGGCCTCGG |
| LasR_NGS_RV8 | GGTACATGAAACGGTTGGCCTCGG |
| LasR_NGS_RV9 | TGTGCATGAAACGGTTGGCCTCGG |
| LasR_NGS_RV10 | TAGACTGAAACGGTTGGCCTCGG |
| LasR_NGS_RV11 | ATGCTTGAAACGGTTGGCCTCGG |
| LasR_NGS_RV12 | GCTGATGAAACGGTTGGCCTCGG |
| LasR_NGS_RV13 | TCTACTGAAACGGTTGGCCTCGG |
| LasR_NGS_RV14 | CTGACTGAAACGGTTGGCCTCGG |
| LasR_NGS_RV15 | GAACTTGAAACGGTTGGCCTCGG |
| LasR_NGS_RV16 | AGTCTTGAAACGGTTGGCCTCGG |
| LasR_NGS_RV17 | CACAATGAAACGGTTGGCCTCGG |
| LasR_NGS_RV18 | GCAAATGAAACGGTTGGCCTCGG |

**Table S8. Coverage correction factor of all samples.**

| Sample | Number of sorted cells | Pool ratio | Adjusted pool ratio | Coverage correction factor, $f_i$ |
|---|---|---|---|---|
| 3OC12-HSL Replicate 1 - bin 1 | 2,000,000 | 0.121118904 | 0.1 | 1.211189037 |
| 3OC12-HSL Replicate 1 - bin 2 | 495,948 | 0.030034339 | 0.033034339 | 0.909185409 |
| 3OC12-HSL Replicate 1 - bin 3 | 581,304 | 0.035203452 | 0.035203452 | 1 |
| 3OC12-HSL Replicate 1 - bin 4 | 390,870 | 0.023670873 | 0.044789777 | 0.528488301 |
| 3OC12-HSL Replicate 2 - bin 1 | 2,000,000 | 0.121118904 | 0.1 | 1.211189037 |
| 3OC12-HSL Replicate 2 - bin 2 | 705,053 | 0.042697623 | 0.042697623 | 1 |
| 3OC12-HSL Replicate 2 - bin 3 | 786,267 | 0.047615899 | 0.047615899 | 1 |
| 3OC12-HSL Replicate 2 - bin 4 | 500,752 | 0.030325267 | 0.05144417 | 0.589479166 |
| 3OHC14-HSL Replicate 1 - bin 1 | 2,000,000 | 0.121118904 | 0.121118904 | 1 |
| 3OHC14-HSL Replicate 1 - bin 2 | 581,325 | 0.035204723 | 0.035204723 | 1 |
| 3OHC14-HSL Replicate 1 - bin 3 | 530,747 | 0.032141747 | 0.032141747 | 1 |
| 3OHC14-HSL Replicate 1 - bin 4 | 427,597 | 0.02589504 | 0.034968478 | 0.74052522 |
| 3OHC14-HSL Replicate 2 - bin 1 | 2,000,000 | 0.121118904 | 0.121118904 | 1 |
| 3OHC14-HSL Replicate 2 - bin 2 | 565,332 | 0.034236196 | 0.034236196 | 1 |
| 3OHC14-HSL Replicate 2 - bin 3 | 491,980 | 0.029794039 | 0.031794039 | 0.937095127 |
| 3OHC14-HSL Replicate 2 - bin 4 | 405,524 | 0.024558311 | 0.034631749 | 0.709127077 |
| Unsorted Pool Replicate 1 | 1,025,000* | 0.062 | 0.05 | |
| Unsorted Pool Replicate 2 | 1,025,000* | 0.062 | 0.05 | |

*The cell counts of unsorted library samples number were determined from the total colonies on agar plates when preparing the library (Methods).

**Table S9. Correction factor for total count calculated from the cell count in bin 1.**

| Sample | Number of sorted cells | % bin unsorted | Calculated total sorted cells | Scaling factor $F_{1j}$ |
|---|---|---|---|---|
| 3OC12-HSL Replicate 1 - bin 1 | 2,000,000 | 0.691 | 2894356 | - |
| 3OC12-HSL Replicate 1 - bin 2 | 495,948 | 0.093 | 5332774 | 1.842473484 |
| 3OC12-HSL Replicate 1 - bin 3 | 581,304 | 0.086 | 6759349 | 2.335355023 |
| 3OC12-HSL Replicate 1 - bin 4 | 390,870 | 0.066 | 5922273 | 2.046145227 |
| 3OC12-HSL Replicate 2 - bin 1 | 2,000,000 | 0.674 | 2967359 | - |
| 3OC12-HSL Replicate 2 - bin 2 | 705,053 | 0.091 | 7747835 | 2.611020451 |
| 3OC12-HSL Replicate 2 - bin 3 | 786,267 | 0.089 | 8834461 | 2.977213247 |
| 3OC12-HSL Replicate 2 - bin 4 | 500,752 | 0.069 | 7257275 | 2.445701797 |
| 3OHC14-HSL Replicate 1 - bin 1 | 2,000,000 | 0.845 | 2366864 | - |
| 3OHC14-HSL Replicate 1 - bin 2 | 581,325 | 0.046 | 12637500 | 5.33934375 |
| 3OHC14-HSL Replicate 1 - bin 3 | 530,747 | 0.042 | 12636833 | 5.339062083 |
| 3OHC14-HSL Replicate 1 - bin 4 | 427,597 | 0.033 | 12957485 | 5.474537348 |
| 3OHC14-HSL Replicate 2 - bin 1 | 2,000,000 | 0.839 | 2383790 | - |
| 3OHC14-HSL Replicate 2 - bin 2 | 565,332 | 0.046 | 12289826 | 5.155582043 |
| 3OHC14-HSL Replicate 2 - bin 3 | 491,980 | 0.043 | 11441395 | 4.799665349 |
| 3OHC14-HSL Replicate 2 - bin 4 | 405,524 | 0.035 | 11586400 | 4.8604948 |

**Table S10. Summary of parameters for NGS data analysis.**

| | 3OC12-HSL replicate 1 | 3OC12-HSL replicate 2 | 3OHC14-HSL replicate 1 | 3OHC14-HSL replicate 2 |
|---|---|---|---|---|
| $\bar{F}$ | 2.075 | 2.678 | 5.384 | 4.939 |
| $f_{bin\ 1}$ | 1.211 | 1.211 | 1.0 | 1.0 |
| $f_{bin\ 2}$ | 0.9092 | 1.0 | 1.0 | 1.0 |
| $f_{bin\ 3}$ | 1.0 | 1.0 | 1.0 | 0.9371 |
| $f_{bin\ 4}$ | 0.5285 | 0.5895 | 0.7405 | 0.7091 |
| $FL_1$ | 3 | 777 | 10285 | 26473 |
| $FL_2$ | 6 | 754 | 10624 | 26059 |
| $FL_3$ | 22 | 278 | 1498 | 4523 |
| $FL_4$ | 18 | 299 | 1537 | 5270 |

### Note S1. Electrocompetent cell preparation

*E. coli* NEB 10-beta was streaked on plate and incubated overnight at 37 °C. A colony was inoculated into 2 mL of LB and cultured overnight at 37°C, 250 rpm. The $OD_{600}$ of the overnight culture was measured and diluted to an $OD_{600}$ of 0.01 using 100 ml fresh LB. Diluted culture was incubated at 37 °C 250 rpm for 2.5 h. Then, the $OD_{600}$ was checked periodically until it reached 0.55 – 0.65. The culture was cooled down in ice/water bath for 15 min to slow growth. The culture was subsequently centrifuged at 8000 rpm, 4 °C for 10 min. The supernatant was discarded, and the remaining cell pellet was washed and resuspended using 100-ml precooled sterilized MQ water on ice/water bath. The resuspended culture was kept on ice for 5 min. The step of centrifugation, discarding the supernatant, and resuspending was repeated once. The supernatant was discarded, and the cell pellet was washed and resuspended using precooled 10% glycerol. The culture was chilled in ice/water bath for 5 min. Then the culture was centrifuged at maximum speed of a centrifuge, 4 °C for 10 min. The supernatant was discarded, and 25 mL 10% glycerol stocks were added to resuspend and aliquot into PCR tubes. The cells were then stored at -70 °C freezer. The temperature of this preparation should be kept low by using ice/water bath, prechilled solutions, and prechilled centrifuge.
